# Whole cell response to receptor stimulation involves many deep and distributed subcellular processes

**DOI:** 10.1101/462200

**Authors:** Jens Hansen, Arjun Singh Yadaw, Mustafa M. Siddiq, Rosa Tolentino, Vera Rabinovich, Gomathi Jayaraman, Mohit Jain, Tong Liu, Hong Li, Joseph Goldfarb, Ravi Iyengar

## Abstract

Neurite outgrowth is an integrated whole cell response regulated by cannabinoid-1 receptor. To understand underlying mechanisms, we identified subcellular processes (SCPs) and their interactions required for the response. Differentially expressed genes and proteins upon receptor stimulation of neuronal cells were used informatically to build networks of SCPs and their interactions. From SCP networks we identified additional genes, which when ablated validated the SCP involvement in neurite outgrowth. The experiments and informatics analyses identified diverse SCPs such as those involved in pyrimidine metabolism, lipid biosynthesis, mRNA splicing and stability along with membrane vesicle and microtubule dynamics. We find that SCPs required for neurite outgrowth are widely distributed among constitutive cellular functions. Several of these SCPs are deep since they are distal to cell signaling pathways and proximal SCPs involved in microtubule growth and membrane vesicle dynamics. We conclude that receptor regulation of SCPs for neurite outgrowth is distributed and deep.

## Introduction

Neuronal differentiation that results in highly connected circuits responsible for brain function starts with the formation of multiple projections called neurites. Subsequently, these projections undergo further differentiation to become either axons or dendrites (Arimura and Kaibuchi 2007, Bradke, Fawcett and Spira 2012). Neurite outgrowth is considered to be among the earliest steps in neuronal differentiation. It can be triggered by wide range of extracellular signals including netrins, growth factors and ligands that regulate GPCRs (Crutcher 1986, Kozlowski et al. 1989, Ziegler and Unsicker 1981). Regulation of neurite outgrowth by GPCRs is often routed through the G_o/i_ pathway. We have studied cannabinoid 1 receptor (Cb1r) regulated neurite outgrowth as a model system for G protein regulated whole cell responses, and have shown that signal flow from G_o_ through Rap to Src and STAT3 control is required (He et al. 2005). A detailed analysis of the upstream signaling network in our laboratory demonstrated the key role of transcriptional regulation and the involvement of multiple transcription factors in Cb1r triggered neurite outgrowth (Bromberg et al. 2008). We sought to identify the downstream subcellular mechanisms by which transcriptional control triggers neurite outgrowth. Identifying the subcellular pathways involved could help elucidate the underlying subcellular mechanisms. To identify the transcriptionally regulated genes involved in early neurite outgrowth we conducted RNA-Seq and proteomics experiments and identified the differentially expressed genes and proteins as the starting point for mapping downstream mechanisms.

The goal of this study was to obtain a genome-wide near-comprehensive description of the regulatory pathways involved in Cb1r regulated neurite outgrowth in Neuro-2A cells and rat primary cortical neurons. We had hypothesized that we might be able to track the relationships between the signaling network we had previously described (Bromberg et al. 2008) and the key cellular processes involved in neurite outgrowth: microtubule growth and vesicle transport and fusion at the growth cone (Tsaneva-Atanasova et al. 2009). However, inspection of the differentially expressed genes and proteins by transcriptomics and proteomics analyses indicated that many diverse subcellular processes (SCPs) were likely to be involved. SCPs are pathways consisting of multiple gene products, with a pathway having an input node and an output effector function that might be biochemical or physiological. There are many signaling pathways, metabolic pathways and pathways with biophysical outputs in a cell. To identify SCPs and relationships between SCPs we used standard cell biological knowledge found in textbooks such as Molecular Biology of the Cell (Alberts et al. 2015), Biochemistry (Berg et al. 2015) and Medical Biochemistry (Rosenthal and Glew 2009) to develop a cell biological ontology called Molecular Biology of the Cell Ontology (MBCO) (Hansen et al. 2017). This ontology allows us to associate genes/proteins with specific SCPs and specify the relationships among SCPs required for a whole cell function such as neurite outgrowth. We analyzed the differentially expressed genes using MBCO and found disparate SCPs are likely to be involved, indicating that receptor regulation of neurite outgrowth was likely to be a distributed phenomenon within the cell. We also found that that many of the genes and proteins that were differentially expressed belonged to SCPs that could be considered both generally required for cell function and often viewed as part of constitutive processes. Since such SCPs are distal from the signaling networks and the proximal functions such as microtubule growth and membrane vesicle transport, we labeled them deep SCPs. We used a combination of bioinformatics analyses and dynamical modeling to understand the relationships among the SCPs involved in neurite outgrowth. To experimentally test the computational predictions regarding the SCPs involved and consequently the underlying mechanisms, we conducted siRNA ablation studies. We selected representative genes of the identified SCPs even if the selected genes themselves were not differentially expressed. Together these experimental and computational analyses show that SCPs related to many diverse subcellular functions come together to mount an integrated whole cell response: neurite outgrowth in response to Cb1r stimulation. The organization of the interacting SCPs indicate that mechanism of neurite outgrowth will be deep and distributed.

## Results

### Chronological activation of distributed interdependent SCPs

To characterize regulatory mechanisms that form the basis for neurite outgrowth (NOG), we stimulated the neuroblastoma cell line N2A with HU210, a potent activator of the Cb1r. Cb1r stimulation increases NOG, thereby offering a model system where a well-defined receptor can trigger a whole cell response. We stimulated N2A cells for 2h, 4h, 6h and 8h with HU210 and investigated gene expression changes by bulk RNASeq and for 5h, 10h and 18h with HU210 and investigated protein expression changes by discovery proteomics (figure 1A, Supplementary table 1).

**Figure 1:**
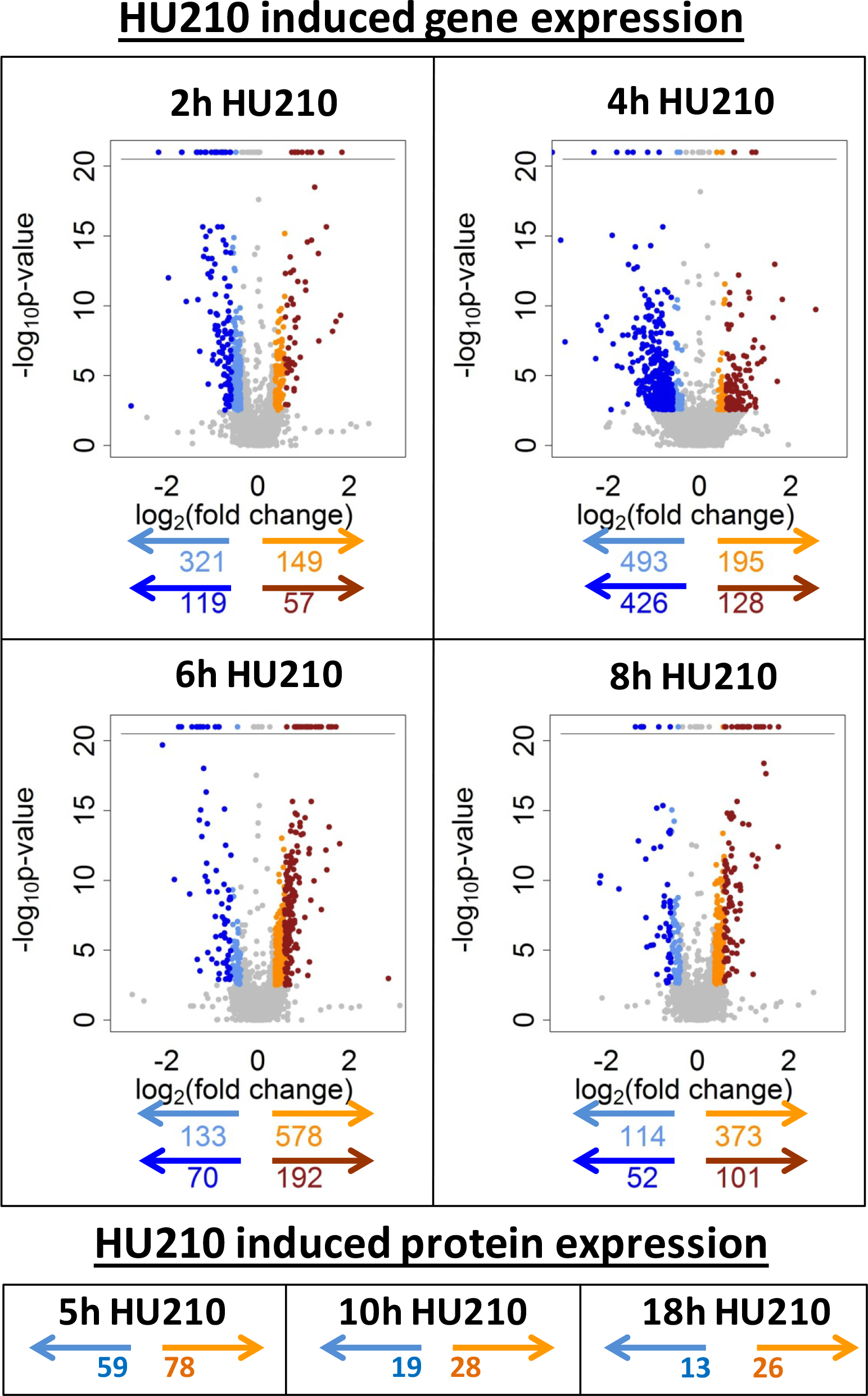
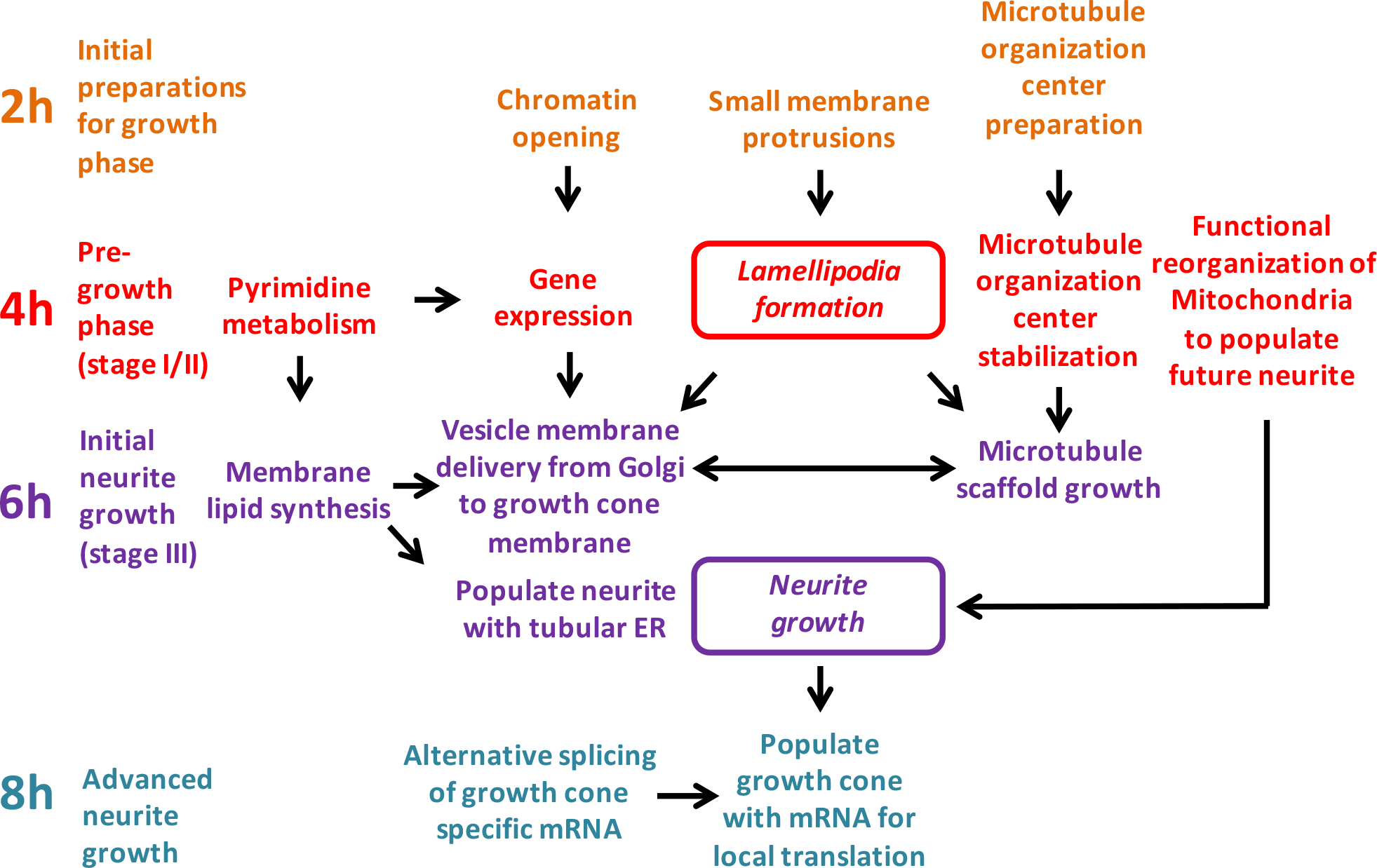
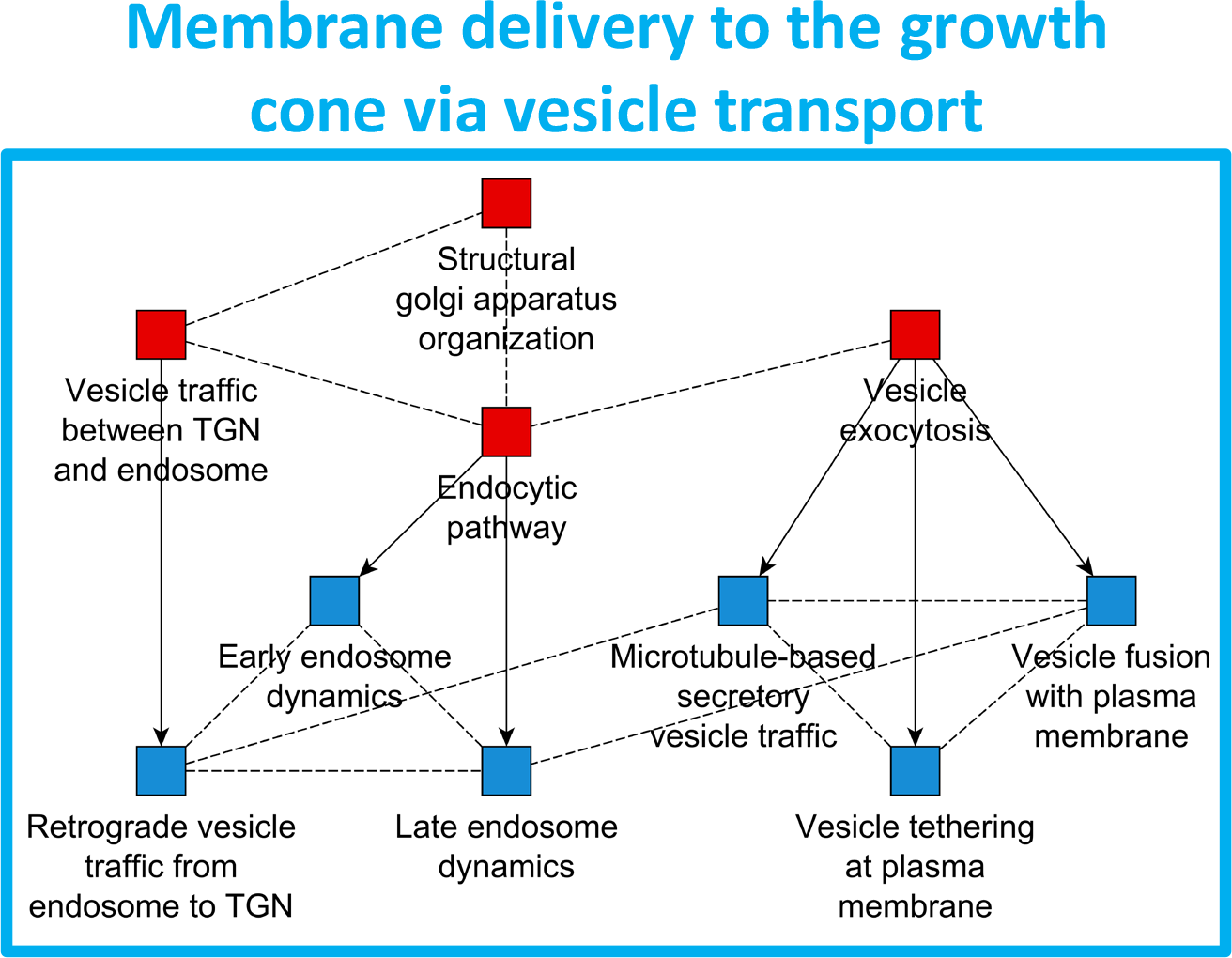
Chronological activation of distributed interdependent SCPs during neurite outgrowth (NOG) matches their dependencies. (A) Neuro2A cells were stimulated with the Cb1R agonist HU210 for indicated time periods, followed by identification of differentially expressed genes (DEGs). Volcano plots show DEGs (FDR 5%). Up- and downregulated genes are in orange and light blue, respectively, if the absolute log_2_(fold change) is between log_2_(1.3) and log_2_(1.5) or in red and blue, if the absolute log_2_(fold change) is larger than log_2_(1.5). Up- and downregulated protein counts are in orange and blue, respectively (nominal p-value 5% and minimum absolute log_2_(fold change) = log_2_(1.1). (B) DEGs were subjected to standard and dynamic enrichment analysis using the Molecular Biology of the Cell Ontology (MBCO) to characterize sub-cellular processes (SCPs) and SCP-networks that are involved in NOG. Predicted SCPs describe multiple sub-cellular functions that are activated in a chronological order that matches their dependencies. Framed SCPs describe functions that can be associated with morphological changes at neurite outgrowth stages II and III. (C) Example SCP-network that was predicted based on up-regulated genes after 6h HU210. Dynamic enrichment analysis identified level-2 and level-3 networks that both describe microtubule vesicle trafficking between the *trans* Golgi network (TGN), endosomes and plasma membrane, suggesting that after 6h HU210, cellular synthesis focuses on proteins involved in membrane delivery from the TGN to the growth cone. The SCP-network also contains an SCP that describes retrograde vesicle traffic from the endosome to the TGN, an initially counter intuitive finding, since membrane back transport should decrease outgrowth velocity. Red nodes: level-2 SCPs, Blue nodes: level-3 SCPs, arrows: hierarchical relationships between parent and child SCPs, dashed lines: horizontal relationships between SCPs of the same level. The predicted SCPs ‘Intracellular bridge abscission’ and ‘Classical complement pathway’ were removed. See suppl. figure 1E for complete dynamic enrichment results.

To identify regulatory SCPs that enable NOG we subjected the differentially expressed genes (DEGs) and proteins (DEPs) to pathway enrichment analysis using the Molecular Biology of the Cell Ontology (MBCO) (Hansen et al. 2017). MBCO has a strict focus on cell biology and represents it in an intuitive way familiar to cell biologists, since it was designed using standard cell biology and biochemistry books as templates. SCPs are hierarchically organized and assigned to levels. SCPs of the same level describe cell biological function of similar detail. Higher level SCPs (i.e. SCPs with lower level numbers) describe more general sub-cellular functions, lower level SCPs (with higher level numbers) more detailed sub-cellular functions. To construct networks of SCPs, the hierarchical tree is enriched using a unique MBCO algorithm that predicts weighted horizontal interactions between SCPs of the same level that lie within and beyond the annotated hierarchy. These interactions enable the dynamic characterization of functionally dependent SCP-networks from experimentally obtained gene lists that underlie the integrated whole cell response. This enrichment algorithm is called dynamic enrichment analysis and serves as a valuable addition to standard enrichment analysis using Fisher’s Exact Test. Briefly, dynamic enrichment analysis searches for all level-2 or level-3 SCPs that contain at least one input gene, and combines them to SCP-unions if they are horizontally connected with a minimum connection weight. The unions are added as function-specific SCPs to the original set of SCPs and the new function-specific MBCO is used for enrichment analysis of the gene list. Results are ranked by p-value and the top 3 level-2 or top 5 level-3 predictions (that are either single SCPs or SCP-unions) are connected with each other based on all MBCO horizontal interactions to generate SCP-networks.

We analyzed up- and down-regulated genes and proteins at each time point after CB1 receptor activation of N2A cells using MBCO and standard as well as dynamic enrichment analysis (Figure 1B-C, Suppl. figure 1). Since we were interested in obtaining near comprehensive maps of downstream mechanisms we looked for all identifiable cellular mechanisms that can drive cellular state change and used two different minimum fold change cutoffs for the DEGs submitted to enrichment analysis. Up- and down-regulated genes and proteins were subjected to standard and dynamic enrichment analysis. Since changes in levels of gene expression were small we used a minimum log_2_(fold change) of +/− log_2_(1.3) to identify DEGs for standard enrichment analysis that tracks SCPs within an annotated category of function. However to identify SCP interactions that arise from genes that might belong to multiple categories by dynamic enrichment we used a more stringent cutoff of a minimum log_2_(fold change) = +/− log_2_(1.5) to identify DEGs (Figure 1A). Enrichment results predicted multiple SCPs and SCP networks that *suggest* the activation of different sub-cellular functions in a chronological order that matches their dependencies (Figure 1B) (suppl. table 2 and 3). SCP-networks at 2h contain microtubule and centrosomal SCPs. Such early expression changes in centrosomal genes could be associated with the localization of the centrosome to the hillock of the future axon, an event that occurs at an early stage of NOG (de Anda et al. 2010, Higginbotham and Gleeson 2007, Tang and Marshall 2012) (suppl. figure 1A/B). Genes associated with nuclear chromatin condensation are down-regulated, suggesting that at this stage the cell prepares for increased gene expression by increasing chromatin accessibility for transcription. Concordantly, up- and down-regulated SCP-networks at 4 hours suggest an increase of gene expression, mainly via up-regulation of ribosomal proteins and down-regulation of CCR4-NOT complex subunits that are involved in bulk mRNA degradation and polymerase II suppression (suppl. figure 1C). Lamellipodia formation is up-regulated (suppl. figure 1D), in agreement with morphological observations that neurites emerge from lamellipodia that form at the earliest stages of NOG (Govek, Newey and Van Aelst 2005). Activity changes in mitochondrial processes indicate mitochondria also prepare for outgrowth (suppl. figure 1D), in agreement with the observation that mitochondria accumulate at the hillock of the future axon before its major growth phase (Bradke and Dotti 1997, Mattson and Partin 1999). Up-regulated pyrimidine synthesis and salvage can support gene expression because they generate the building blocks for mRNA synthesis and suggest that the cell is preparing for an increase in lipid membrane production, since the pyrimidine nucleoside triphosphates CTP and UTP are cofactors during membrane lipid synthesis (Garavito, Narváez-Ortiz and Zimmermann 2015). The up-regulated SCP-network at 6 hours describes microtubule-based vesicle trafficking between the Golgi apparatus and the plasma membrane (figure 1C, suppl. figure 1E), indicating a cellular focus on membrane delivery from the lipid synthesis site in the cell body to the growth cone where new membrane is added to the growing neurite shaft (Pfenninger 2009, Nakazawa et al. 2012). In agreement, standard enrichment analysis (suppl. figure 1F) predicted up-regulation of membrane lipid metabolism. Up-regulated microtubule plus-end tracking (suppl. figure 1F) further supports that DEGs at 6h refer to the main neurite growth phase that is characterized by parallel and interdependent growth of the neurite shaft and microtubule neurite scaffold. 8h DEGs enrich for multiple SCPs and SCP-networks, including the upregulation of splicing activities and further modification of vesicle trafficking (including retrograde vesicle trafficking) and membrane lipid synthesis (suppl. figures 1G/H).

DNA repair processes are predicted to be down-regulated after 2h, 4h, and 6h HU210 treatment (Suppl. figures 1A/C/E) and up-regulated after 6h HU210 treatment (Suppl. figure 2). Such SCP activities could be involved in the regulation of gene expression (Madabhushi et al. 2015).

Downstream analysis of DEPs confirmed many of the SCPs identified based on gene expression changes. Dynamic and standard enrichment analysis of DEPs after 5h HU210 treatment predicted the involvement of both up- and downregulated proteins in ribonucleoprotein synthesis, mRNA translation, mRNA splicing and nuclear protein translocation (suppl. figures 1I and 3, supplementary tables 4 and 5). Additionally, upregulated proteins were part of SCPs involved in microtubule growth and proteasomal degradation and downregulated proteins were part of SCPs that can be related to the microtubule organization center. At later timepoints (10h, 18h) only a few proteins were differentially expressed (figure 1A) that were involved in multiple SCPs among which are SCPs related to centrosome organization, microtubule growth, mRNA metabolism and protein processing during the biosynthetic pathway (supplementary figures 1J and 1K and 3, supplementary tables 4 and 5).

### Neurite outgrowth requires many metabolic processes

To document the involvement of the predicted SCPs in neurite outgrowth we selected representative genes for siRNA knock down. Our aim was not to analyze if a certain gene is involved in a particular SCP, but if a particular SCP is involved in NOG. Therefore, when possible, we selected genes whose knock down has been shown to influence the corresponding SCP in other cell types, independent of its transcriptional regulation during NOG. Indeed, most of the genes selected for siRNA knockdown were not identified as DEGs or DEPs (Table 1).

**Table 1:**
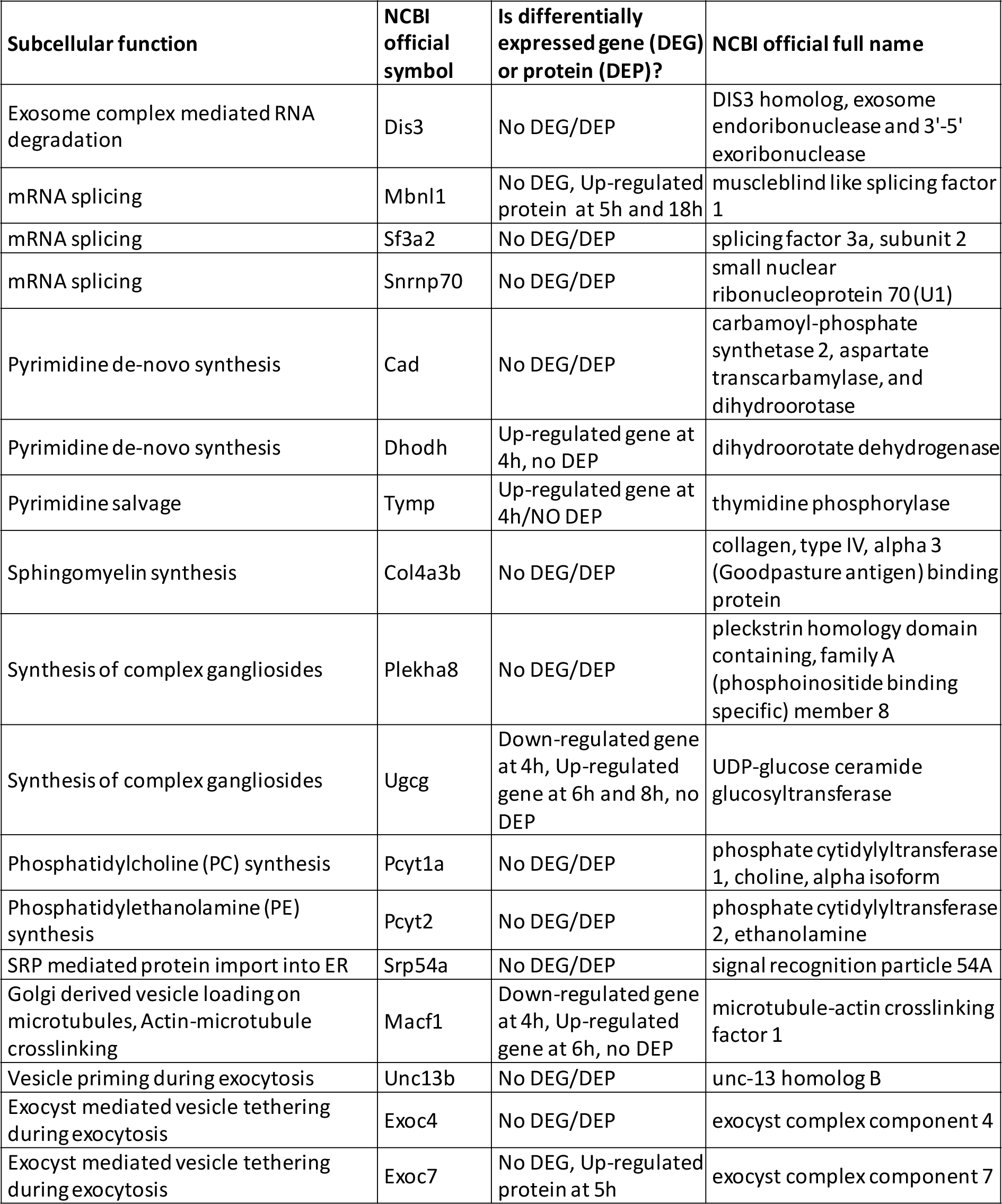
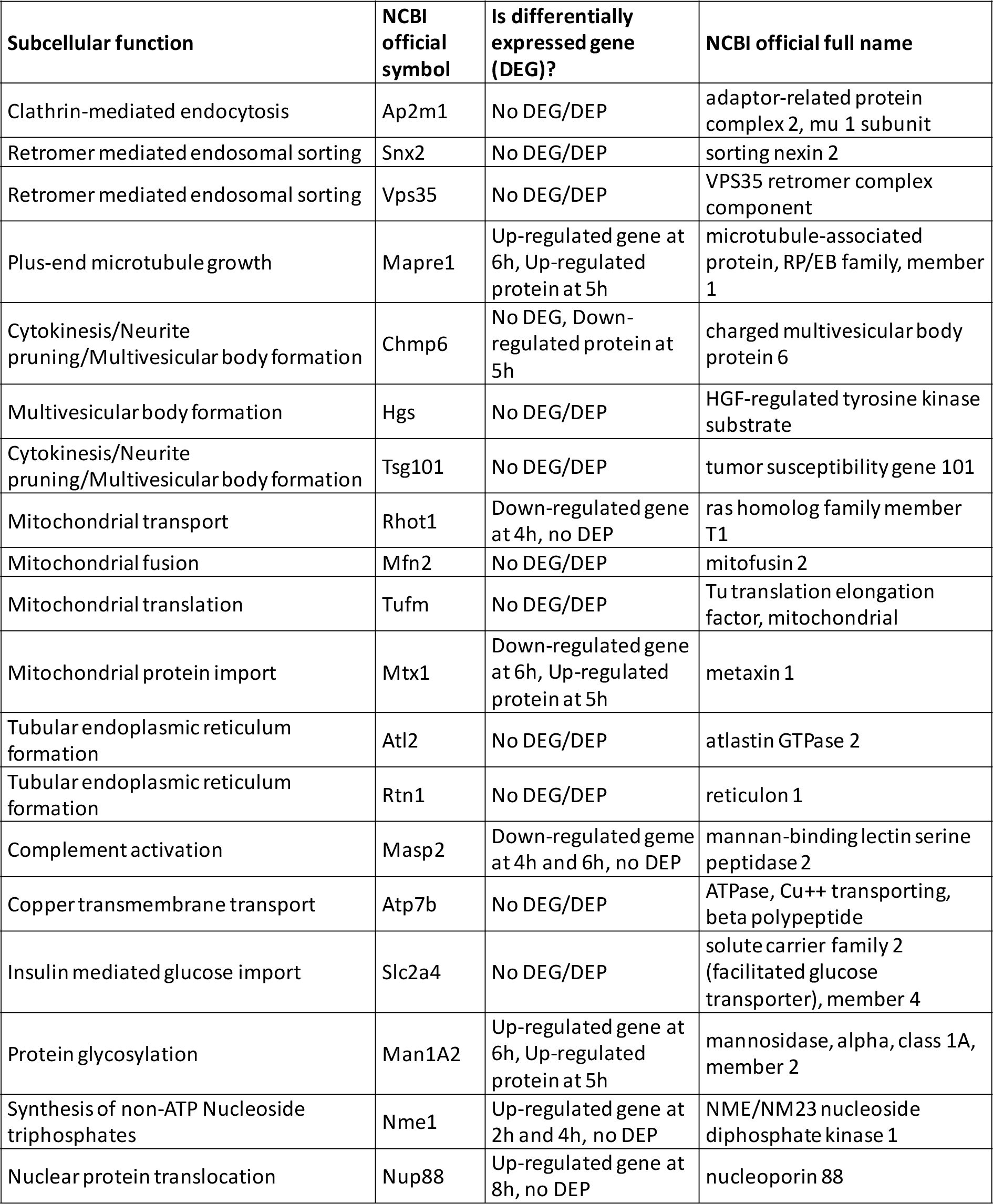
Selected genes for siRNA knock down screening. We selected representative genes for the indicated sub-cellular functions to test the effect of their knock down on neurite outgrowth. Most of the selected genes were not among the differentially expressed genes (No DEG) or differentially expressed proteins (No DEP) after HU210 treatment of N2A cells.

Primary rat cortical neurons were seeded on microfluidic chambers that contain two sides that are separated by a microgroove sidewall (figure 2A). Plated on the left side of the chamber, the cell bodies cannot cross to the right side, but their neurites can grow through small slits in the sidewall into the right side, allowing an easy quantification of outgrowth length. After siRNA transfection we waited for 48h to allow siRNA mediated gene silencing and performed axotomy to reset neurite growth length in all chambers to 0 (i.e. to the microfludic sidewall). Neurite outgrowth was documented after another 48h. After quality control (suppl. Figure 4) we quantified average growth densities at any distance from the microgroove wall. Average growth density showed a steady decrease along the outgrowth direction. Neurite outgrowth densities of each experiment were normalized towards control density at 100 μm which was set to 100%. Since NOG densities differed between different experiments, we identified those distances within each experiment at which control NOG density had fallen to 75%, 50% and 25%. NOG densities of siRNA treated samples were determined at these reference distances and analyzed for significant difference from 75%, 50% or 25% (i.e. control values). Any gene knock down that showed a nominally significant change of growth density (alpha = 0.05) of at least one of these 3 distances was defined to significantly regulate NOG. In a few cases where significance was closely missed (p-value > 0.05 and ≤ 01) we stated that the gene may influence NOG.

**Figure 2:**
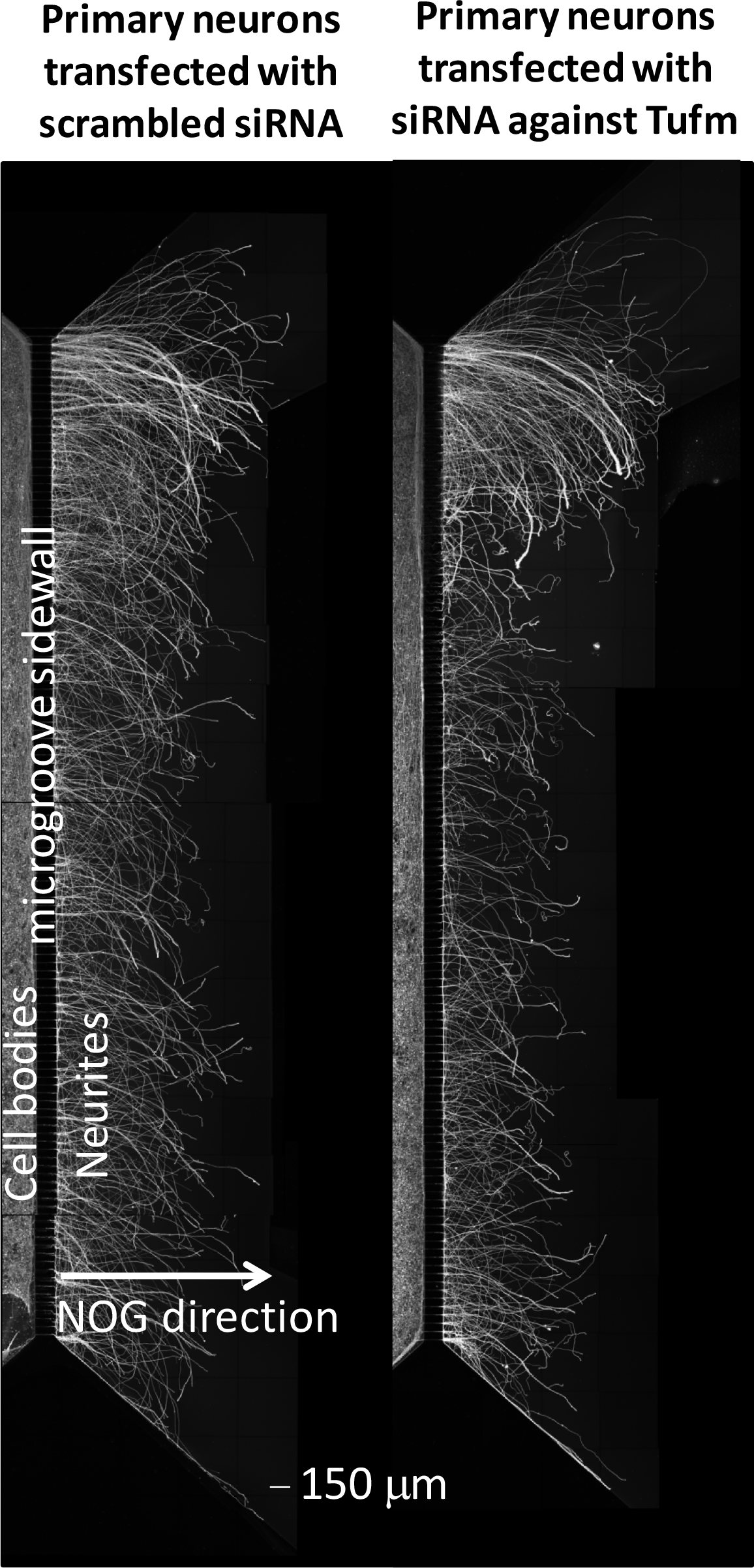
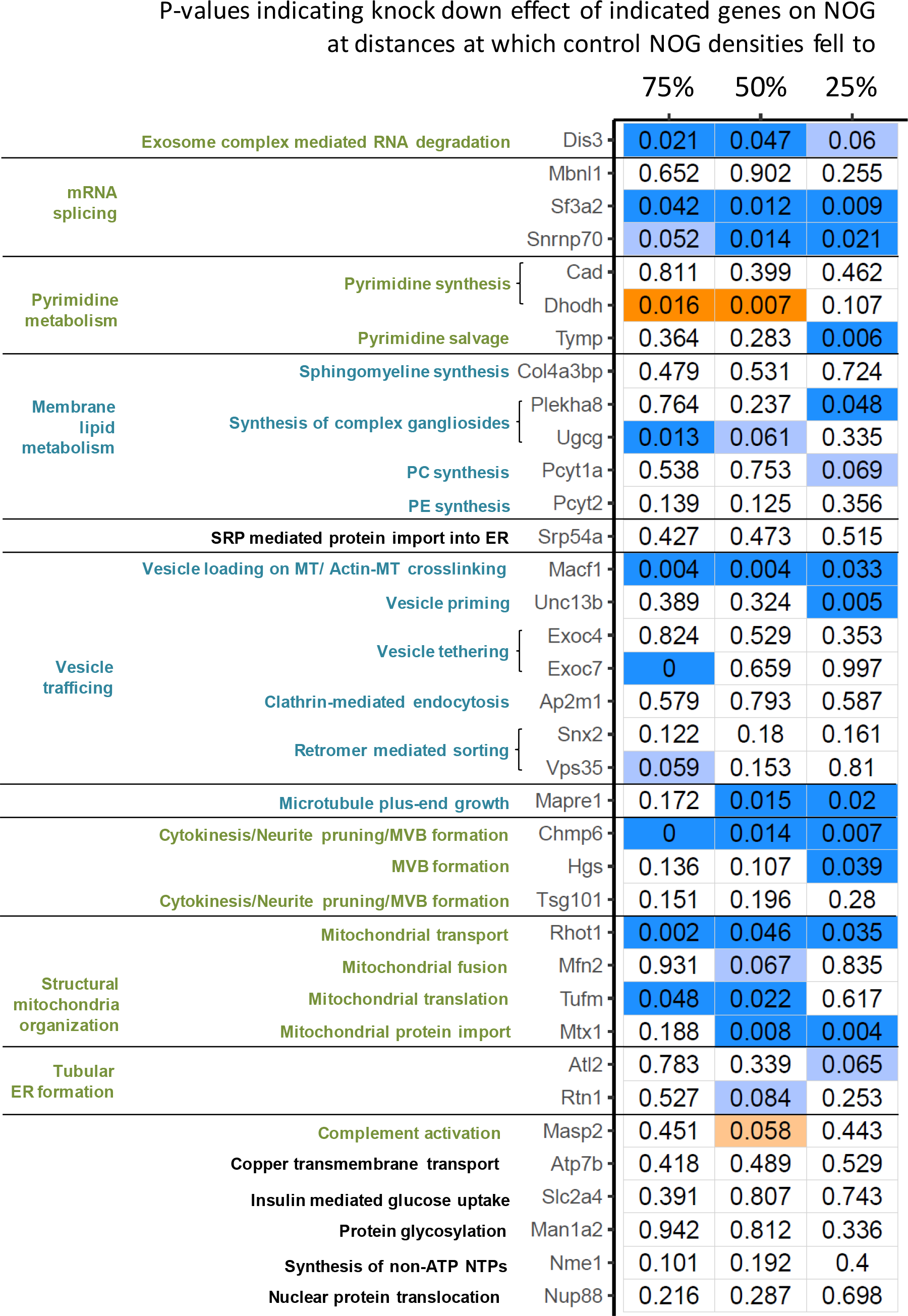
siRNA knock down experiments document involvement of multiple SCPs. (A) P1 rat cortical neurons were transfected with siRNA against selected representative pathway genes (shown: Tufm) or scrambled siRNAs. To analyze neurite growth under gene silencing conditions we waited for 48h after transfection, before resetting neurite growth to the edge of the microgroove via axotonomy. Another 48h later we stained neurites and cell bodies for β-III tubulin to document siRNA effect on neurite outgrowth. Each shown image is a composite of three different overlapping images that showed the top, middle and bottom part of the microchamber and were manually stitched. (B) Neurite outgrowth fluorescence intensities were quantified and the average intensity at each distance from the microgroove sidewall calculated. All intensities of one experiment were normalized towards control average intensity at 100 μm that was set to 100% (Multiple controls within one experiment were averaged before normalization). Since outgrowth results showed great variations between the different experiments, we searched for those distances at which control average intensities had fallen to 75%, 50% and 25%. We determined the outgrowth intensity of each siRNA treated sample at these three distances, removed outliers via Dixons-Q-test and used a one-sample two-tailed t-test to analyze, if the outgrowth intensity after knock down of a particular gene is significantly different from the reference control densities. Nominal p-values are shown and labeled dark blue or dark orange, if knock down significantly inhibited or stimulated neurite outgrowth (p-value <= 0.05), respectively, and light blue or light orange, if knock down may inhibit or stimulat NOG (p-value > 0.05, <= 0.1), respectively. SCPs that are regulated by the indicate genes are added and colored in blue, if they can be directly associated with neurite shaft and scaffold growth and in green, if they describe deep more basal regulatory functions. Black SCPs belong to genes that did not show an effect on neurite outgrowth. PC: Phosphatidylcholine, PE: Phosphatidylethanolamine, SRP: Signal recognition particle, ER: Endoplasmic reticulum, MT: Microtubule, MVB: Multi vesicular bodies. See supplementary figure 5 for detailed results.

We could verify the involvement of multiple SCPs that we identified in our RNASeq screening approach (Figure 2B, Suppl. Figure 5). Among these were SCPs that can be directly linked to neurite shaft growth such as membrane lipid production and membrane delivery to the growth cone via vesicle trafficking, as well as SCPs that describe deep regulatory functions that contribute to NOG on a lower more basal level (e.g. mRNA splicing).

#### Pyrimidine Biosynthesis Pathways

Pyrimidine synthesis and salvage was predicted to be up-regulated after 4h HU210 treatment. Pyrimidines are substrates for multiple sub-cellular pathways such as DNA and RNA synthesis and are co-factors during the synthesis of phosphatidylethanolamine (PE), phosphatidylcholine (PC) and complex gangliosides (Figure 3A) (Garavito et al. 2015). The up-regulation of pyrimidine synthesis and salvage could, therefore, support increased gene expression by delivering the building blocks for mRNA synthesis. Additionally, an increased pool of the pyrimidine nucleoside triphosphates CTP and UTP could support phospholipid and ganglioside synthesis, SCPs that are up-regulated after 6h HU210 treatment. CTP is a co-factor for neuronal phosphatidylethanolamine and -choline synthesis via the Kennedy pathway (Vincenzetti et al. 2016) and uridine induces intracellular CTP increase and neurite outgrowth (Pooler et al. 2005). UTP and CTP are co-factors for ganglioside synthesis (Sandhoff and Sandhoff 2018). Though pyrimidine de novo synthesis was also up-regulated, we could only document the dependence of NOG on pyrimidine salvage (Tymp), but not on pyrimidine de-novo synthesis (Cad knock down showed no effect, Dhodh knock down increased NOG) (Figure 2B, Suppl. Figure 5). These results are in agreement with a mechanism that favors pyrimidine recycling. De novo synthesis of pyrimidines is associated with high energy cost, so pyrimidine recycling is favored to maintain a sufficient pool for pyrimidine dependent pathways (Vincenzetti et al. 2016). The source for recycled substrates might either be pyrimidine metabolites that were generated in the same cell (e.g. after DNA/RNA degradation) or metabolites that were produced by astrocytes, since we used astrocyte conditioned media, to generate an environment that resembles the in-vivo situation in the CNS.

**Figure 3:**
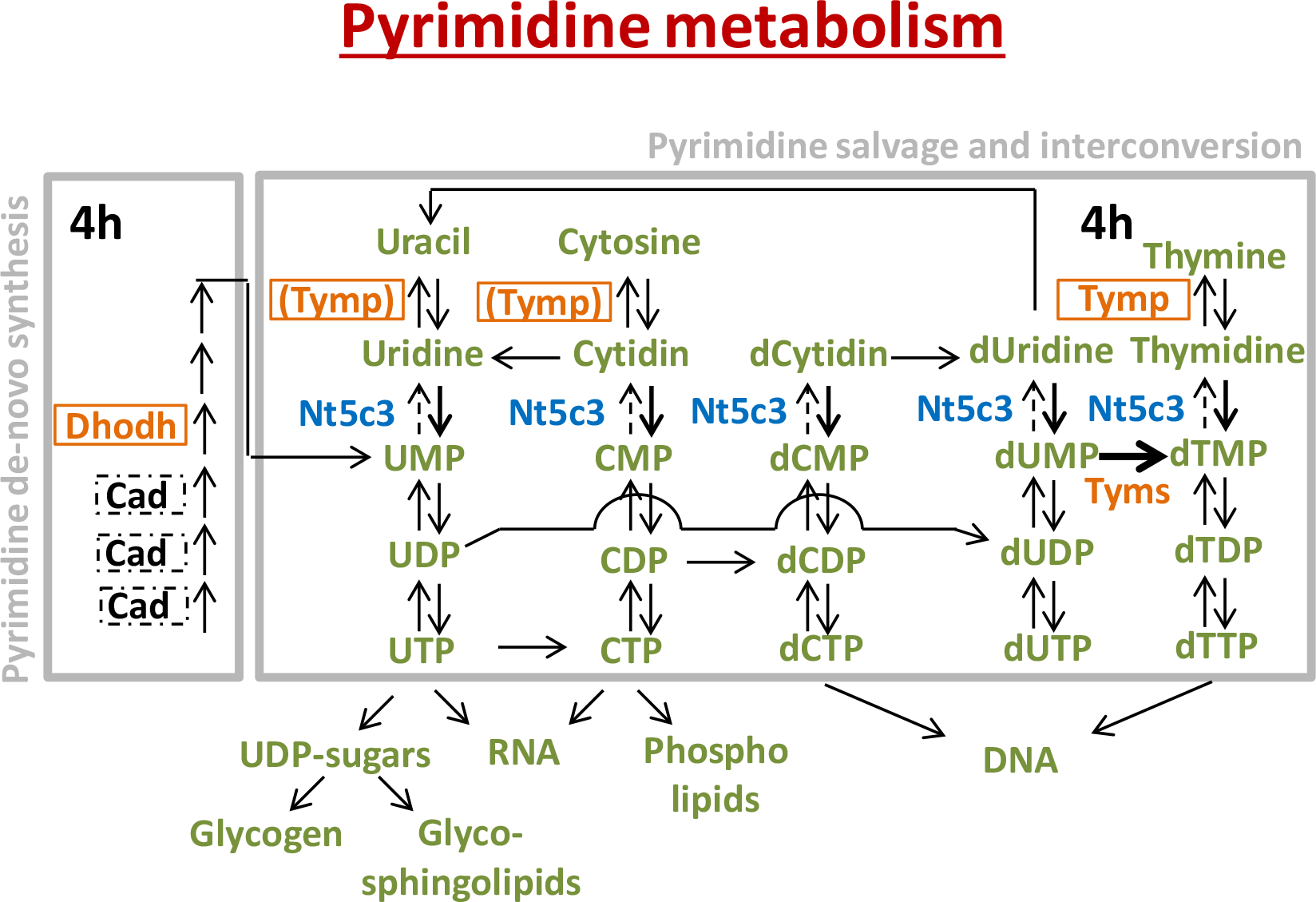
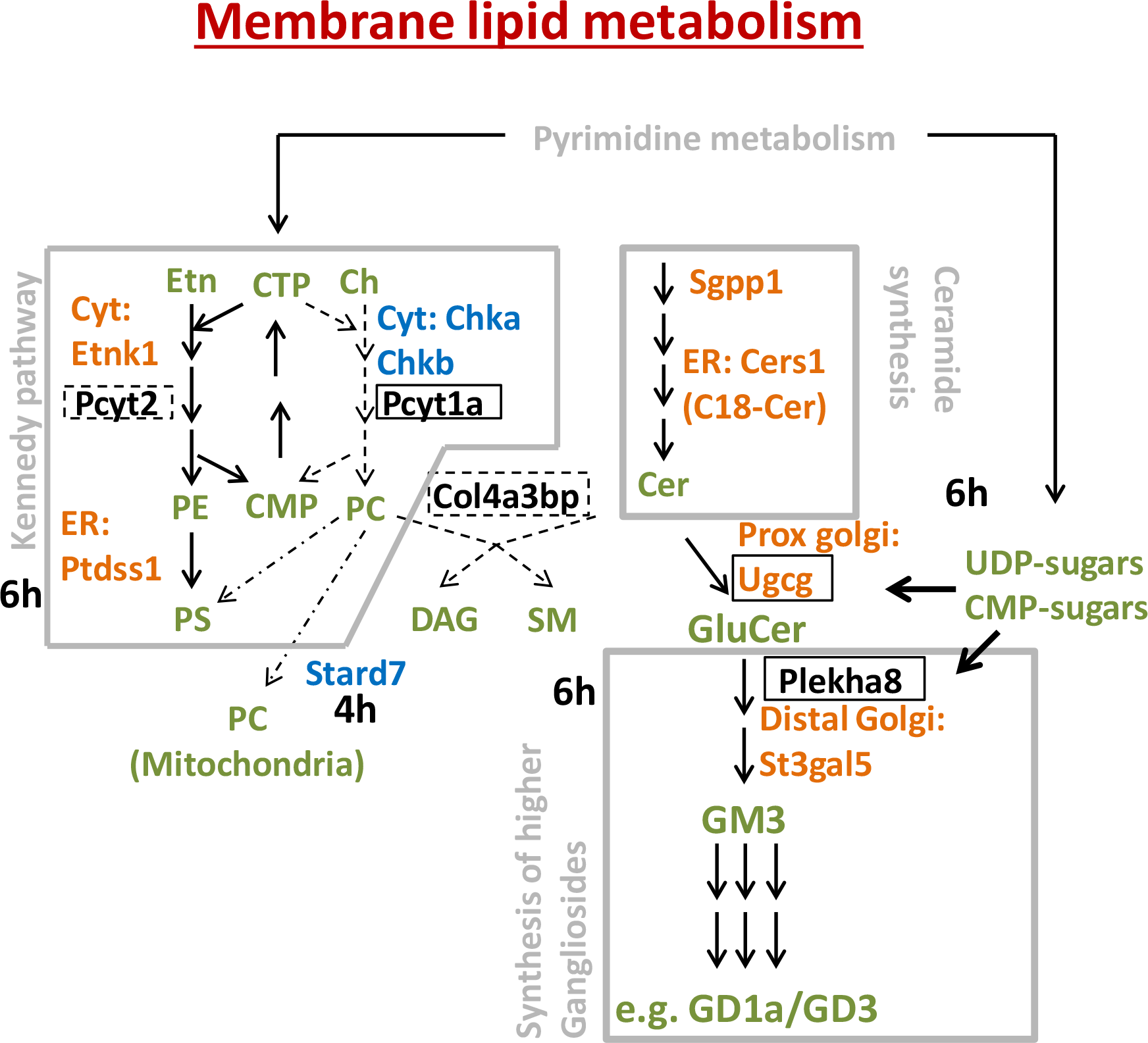
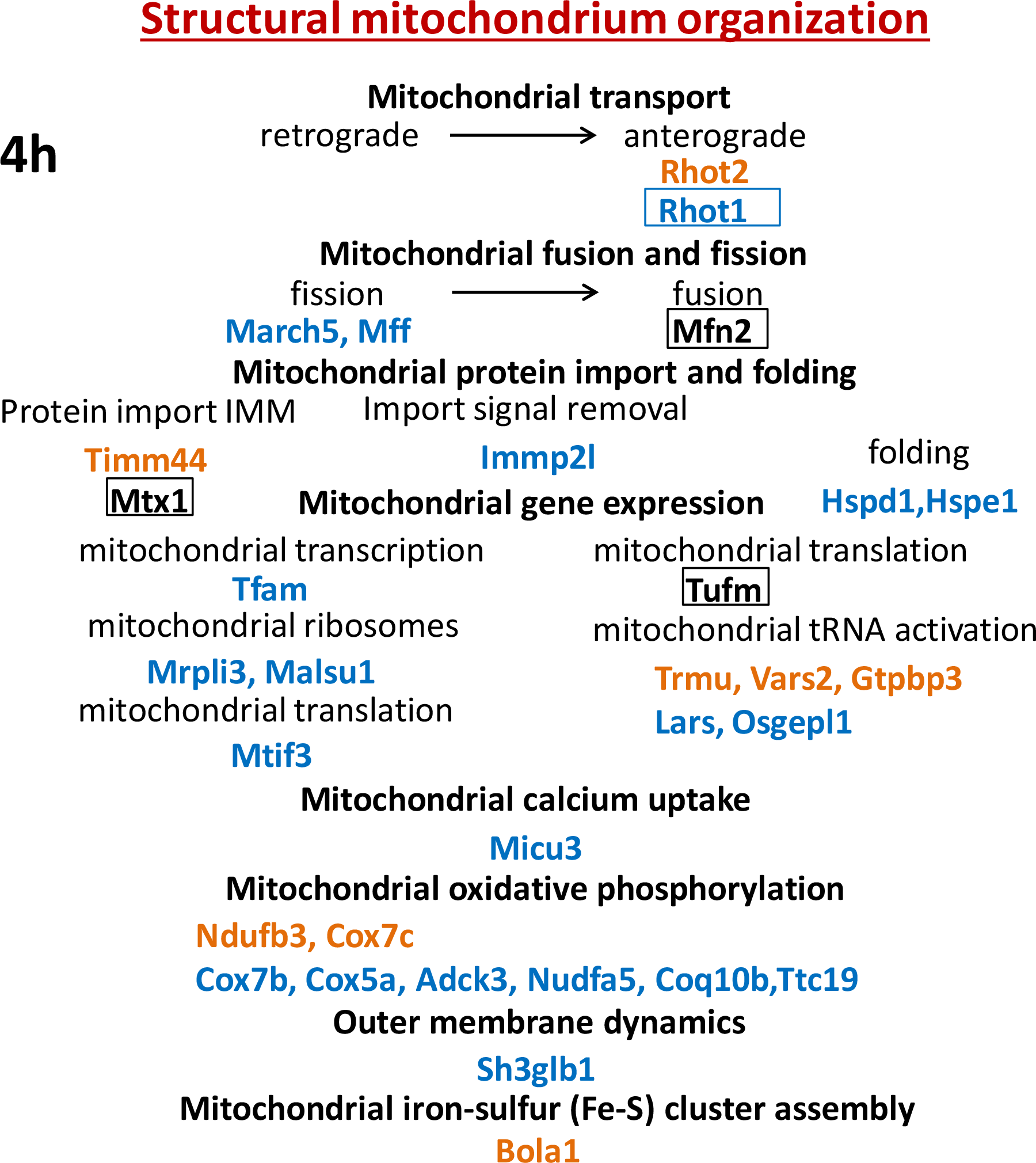
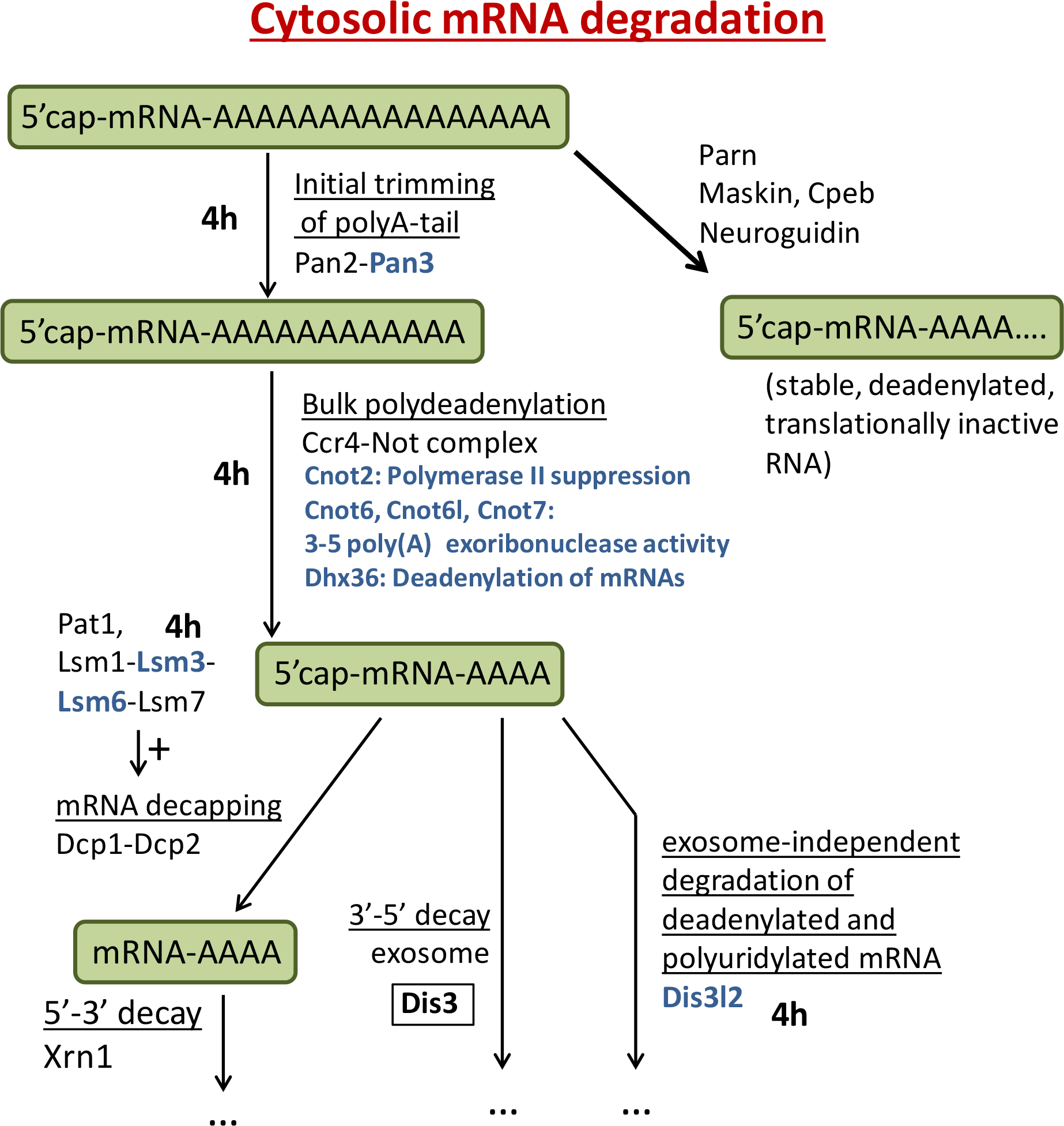
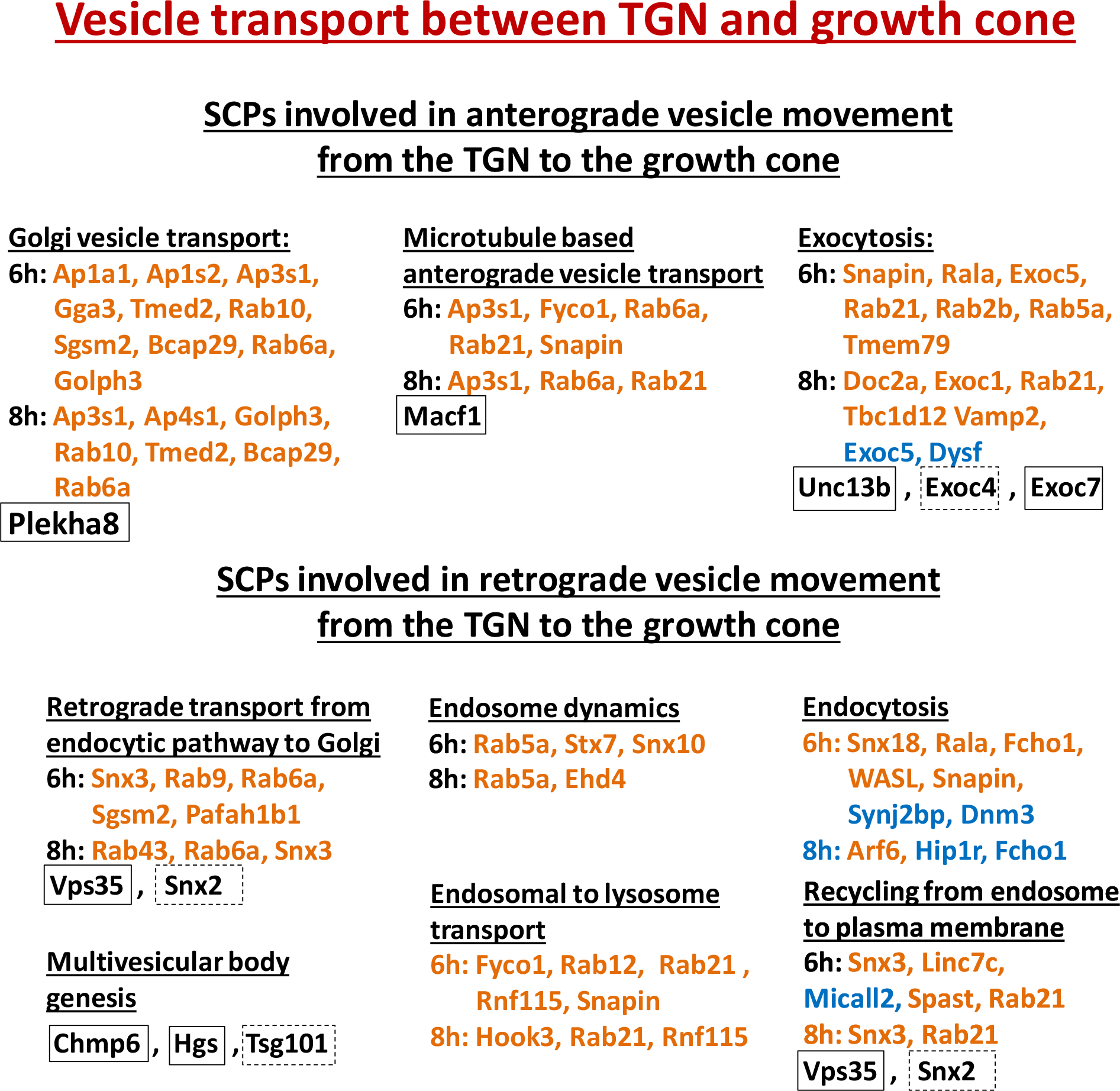
Enzymatic reactions and gene products involved in basic cellular metabolism, structural mitochondrium organization and vesicle transport. (A) Enzymatic reactions involved in pyrimidine de-novo synthesis and pyrimidine salvage and interconversion (adapted from (Garavito et al. 2015) with modifications from (Vincenzetti et al. 2016)). Up- (orange) and down-regulated (blue) genes suggest a net flux of pyrimidine metabolites to pyrimidines that can be used to generate complex molecules. siRNA knock down results suggest the dependence of NOG on pyrimidine salvage, but not on pyrimidine de novo synthesis. Orange/blue: up-/down-regulated genes (based on a minimum log_2_(fold change) of +/− log_2_(1.3)). Green: metabolites. Genes surrounded by solid rectangles: siRNA targets with an effect on NOG. Genes surrounded by dashed rectangles: siRNA targets with no effect on NOG. (B) Enzymatic reactions and transport proteins involved in phospholipid, ceramide and ganglioside synthesis. Up- and down regulated genes as well as siRNA knock down results suggest a shift of phospholipid production from PC to PE/PS and an increase in ganglioside production. Etn: Ethanolamine, Ch: Choline, PE: Phosphatidylethanolamine, PS: Phosphatidylserine, PC: Phosphatidylcholine, Cer: Ceramide GM3/GD1a/GD3: GM3/GD1a/GD3 ganglioside, GluCer: Glucosylceramide, SM: Sphingomyelin, DAG: Diacylglycerol (C) Up- and down-regulated genes and siRNA targets involved in Structural mitochondria organization. (D) Up- and down-regulated genes in cytosolic mRNA degradation (Norbury 2013). (E) Up- and down-regulated genes in vesicle transport between the TGN and growth cone.

#### Lipid Biosynthesis Pathways

The distribution of up- and down-regulated genes within membrane lipid synthesis pathways suggests an increase in the synthesis of C18 ceramide and complex gangliosides (Figure 3B) and a shift in phospholipid synthesis from phosphatidylcholine (PC) to phosphatidylethanolamine (PE) and phosphatidylserine (PS). Genes involved in lipid synthesis pathways are up-regulated in parallel to genes involved in vesicle transport, suggesting the functional coupling of both pathways for membrane synthesis and delivery to the growth cone.

Complex gangliosides contribute 10 to 20% of lipids in neuronal membranes (Posse de Chaves and Sipione 2010) and are synthesized using ceramide as a substrate (Yu et al. 2004). They are primarily localized in the outer leaflet of the plasma membrane (Yu et al. 2011), suggesting that ceramide and ganglioside synthesis are up-regulated to generate the building blocks of the outer leaflet of the growing neurite. In agreement with these observations, we find that knock down of genes that are involved in ganglioside synthesis, either as an enzyme (Ugcg) (Daniotti and Iglesias-Bartolomé 2011) or as a lipid transport protein (Plekha8) that directs glycosylceramide from the cis-Golgi to the synthesis site of complex gangliosides at the trans-Golgi (D’Angelo et al. 2007, Yamaji et al. 2008, Lannert et al. 1998), inhibit neurite outgrowth (Figure 2B, Suppl. Figure 5). In contrast, the knock down of Col4a3bp, a lipid transport protein that directs glycosylceramide to sphingomyelin synthesis sites at the Golgi (Hanada et al. 2003, Hanada et al. 2009, Yamaji et al. 2008) did not influence NOG (Figure 2B, Suppl. Figure 5). Our results are in agreement with experimental data suggesting neuronal sphingomyelin synthesis at a non-Golgi location (Sadeghlar, Sandhoff and van Echten-Deckert 2000, van Echten et al. 1990).

PE and PS mainly localize to the inner leaflet of the plasma membrane (Fadeel and Xue 2009), suggesting that the up-regulation of enzymes involved in PE and PS synthesis via the Kennedy pathway increases the production of the building blocks of the inner leaflet of the growing neurite. PC, the most abundant phospholipid in cell membranes, is largely localized at the ER (Tracey et al. 2018) and serves as a storage reservoir for choline that is used for synthesis of the neurotransmitter acetylcholine (Blusztajn et al. 1987a, Blusztajn, Liscovitch and Richardson 1987b). The two different arms of the Kennedy pathway that synthesize PE (that can be converted into PS) and PC both utilize diacylglycerol (DAG) as a substrate and the pyrimidine nucleoside triphosphate CTP as a co-factor (Vincenzetti et al. 2016). The predicted down-regulation of PC synthesis might further favor the production of PE and PS by increasing the availability of both molecules for PE synthesis. Knock down of Pcyt2, an enzyme involved in PE synthesis via the Kennedy pathway, may decrease NOG, since we could document reductions of NOG in three independent experiments, though the p-value was not significant due to large experimental variations (Figure 2B, Suppl. Figure 5). Similarly, the knock down of Pcyt1a, an enzyme involved in PC synthesis via the Kennedy pathway, may also decrease NOG (with a p-value close to significance). Although our DEGs suggest a shift from PC to PE and PS synthesis to increase neurite membrane production, NOG may still depend on a sufficient supply of PC as the most abundant phospholipid in cell membranes (Carter et al. 2008).

#### Mitochondrial Functions

Differential expression of genes at 4h HU210 that are involved in structural mitochondrial organization suggest a shift from Rhot1 to Rhot2 mediated anterograde mitochondrial transport, a shift from mitochondrial fission to mitochondrial fusion, the down-regulation of mitochondrial gene expression, and activity changes in SCPs that are involved in mitochondrial protein import, mitochondrial gene expression and oxidative phosphorylation (Figure 3C). Knock down of 3 mitochondrial genes (Rhot1, Tufm and Mtx1) significantly inhibited neurite outgrowth, while knock down of another gene (Mfn2) may inhibit NOG (based on a p-value close to significance) (Figure 2B, Suppl. Figure 5), documenting the importance of the SCPs mitochondrial anterograde transport, fusion, translation and protein import. Mitochondria populate the growing neurite where they increase their length through fusion (Voccoli and Colombaioni 2009). Our results suggest that mitochondria are first prepared for populating the neurite before the neurite enters its major growth phase, since mitochondrial genes are up-regulated before the up-regulation of genes involved in membrane synthesis and vesicle trafficking (figures 1B and 1C, Suppl. figures 1D, 1E and 1F). This interpretation complements morphological observations showing mitochondria, peroxisomes and Golgi vesicles accumulation at the base of that neurite that becomes the future axon before it enters its major growth phase (Mattson and Partin 1999, Bradke and Dotti 1997).

#### Regulation of mRNAs

In parallel to the up-regulation of ribosomal genes after 4h HU210 treatment, we observed multiple downregulated genes that participate in bulk mRNA degradation (figure 3D). Our results suggest that an increase in gene expression that is needed for continuous synthesis of neurite components for the following neurite growth phase (6h) is facilitated by an increase in ribosomal translation and mRNA lifetime. Down-regulated genes after 4h HU210 treatment are part of multiple SCPs that participate in mRNA degradation at multiple consecutive steps (Norbury 2013): initial mRNA-polyA tail trimming, bulk mRNA polydeadenylation by the CCR4-NOT complex, mRNA decapping and exosome-independent degradation of deadenylated and polyuridylated mRNA (De Almeida et al. 2018). The CCR4-NOT complex regulates numerous steps during mRNA biogenesis including transcription, mRNA export and mRNA degradation (Collart 2016, Doidge et al. 2012, Ukleja et al. 2016, Villanyi and Collart 2015). Three of the five down-regulated CCR4-NOT complex subunits have 3’-5’ poly(A) exoribonuclease activity (Cnot6, Cnot6L, Cnot7) (Doidge et al. 2012), suggesting that down-regulation of these complex components increases mRNA lifetime.

siRNA knock down of the putative catalytic component Dis3 of the exosome complex (Kilchert, Wittmann and Vasiljeva 2016, Siwaszek, Ukleja and Dziembowski 2014) that is involved in another degradation pathway of polydeadenylated mRNA decreases NOG (Figure 2B, Suppl. Figure 5), indicating that mRNA degradation, possibly of mRNAs that contain errors, is still a necessary function to secure neurite outgrowth.

Components of the splicing machinery are down-regulated after 4h HU210 treatment, i.e. at the timepoint where other enrichment results suggest an increase in gene expression, and are up-regulated after 8h HU210 treatment, i.e. after the up-regulation of the machinery for neurite shaft and scaffold growth at 6h. This chronological order suggests that down-regulation of the splicing machinery after 4h HU210 treatment contributes to the increase of gene expression to ensure continuous production of proteins at the cell body for the following initial neurite growth phase. The late timepoint of the up-regulation of the splicing machinery at 8h suggests that a higher concentration of splicing machinery components is needed to contribute to the advanced neurite growth phase. Axonal growth cones contain a reservoir of translationally silent mRNAs (Kim and Jung 2015) and the machinery for local translation, protein folding and posttranslational modification (Estrada-Bernal et al. 2012). More than half of the neurite-localized proteome might have been generated by local mRNA translation (Zappulo et al. 2017). Alternative splicing of 3’ UTRs localizes neuronal mRNA molecules to neurites (Taliaferro et al. 2016). Restriction of spliceosomal components alters splicing of a subset of mammalian mRNAs with weak 5’ splice sites (Wickramasinghe et al. 2015). Our results suggest that low concentrations of the splicing machinery might favor splicing to produce mRNA for protein production in the cell body, while high concentrations increase alternative splicing to generate mRNAs that are transported into the neurite for local protein production.

The involvement of splicing in neurite outgrowth was documented by our knock down results. Knock down of the spliceosomal components Sf3a2 and Snrnp70 significantly inhibited NOG (Figure 2B, Suppl. Figure 5).

#### Membrane Vesicle Transport

DEGs related to vesicle transport processes are related to multiple steps in vesicle transport between the plasma membrane, endosome and TGN (Figure 3E). Our siRNA screening supported this observation, since we could document that multiple SCPs ranging from vesicle loading on the microtubule after budding at the TGN (Burgo et al. 2012), vesicle exocytosis and retromer based endosomal sorting (Gallon and Cullen 2015, Zhang et al. 2018) are involved in NOG (Figure 2B, Suppl. Figure 5). The gene Vps35 is a major component of the retromer that is involved in protein sorting from the endosome to the plasma membrane and to the Golgi. Its knock down inhibited NOG (Figure 2B, Suppl. Figure 5). Knock down of SNX2 had no effect on NOG. Within the retromer SNX2 shows functional redundancy with SNX1 (Rojas et al. 2007), offering an explanation for the missing effect of its knock down. In combination with the observed up-regulation of genes involved in retrograde transport from the endosome to the Golgi (Figure 1C, Suppl. Figures 1E and 1H), these results indicate that an increase in back transport from the growth cone to the TGN is needed to increase NOG outgrowth velocity.

To further investigate this counter-intuitive finding we used our multicompartment ODE model that integrates membrane production, membrane delivery to the growth cone via vesicle transport and microtubule growth during NOG (Yadaw et al. 2017). The simulations were focused on understanding how membrane back-transport regulates NOG. In this model we start with the postulate that individual SCPs need to be in balance to achieve neurite outgrowth without violation of predefined model constraints. The model simulates anterograde (forward) and retrograde (backward) vesicle trafficking between the TGN and the growth cone (Figure 4A). Anterograde vesicles bud at the TGN and move through the neurite shaft to the growth cone cytoplasm via kinesin-mediated transport along the microtubules. Kinesin is attached to the vesicles via kinesin receptors that are integral parts of the vesicle membrane. Anterograde vesicles within the growth cone cytoplasm fuse with the growth cone plasma membrane after SNARE complex formation between vesicle(v)-SNAREs V and target(t)-SNAREs at the plasma membrane. Any additional membrane at the growth cone is added to the neurite shaft, causing its growth. Retrograde vesicles are generated via endocytosis at the growth cone membrane, transported backwards along the microtubules via dynein mediated vesicle transport into the cell body cytoplasm and fuse with the TGN. Since some but not all of kinetic parameters are known, part of our NOG model is an analytical solution that predicts kinetic parameter sets that allow NOG growth at a certain velocity (an experimentally measured high level constraint) without violation of SCP model constraints.

**Figure 4:**
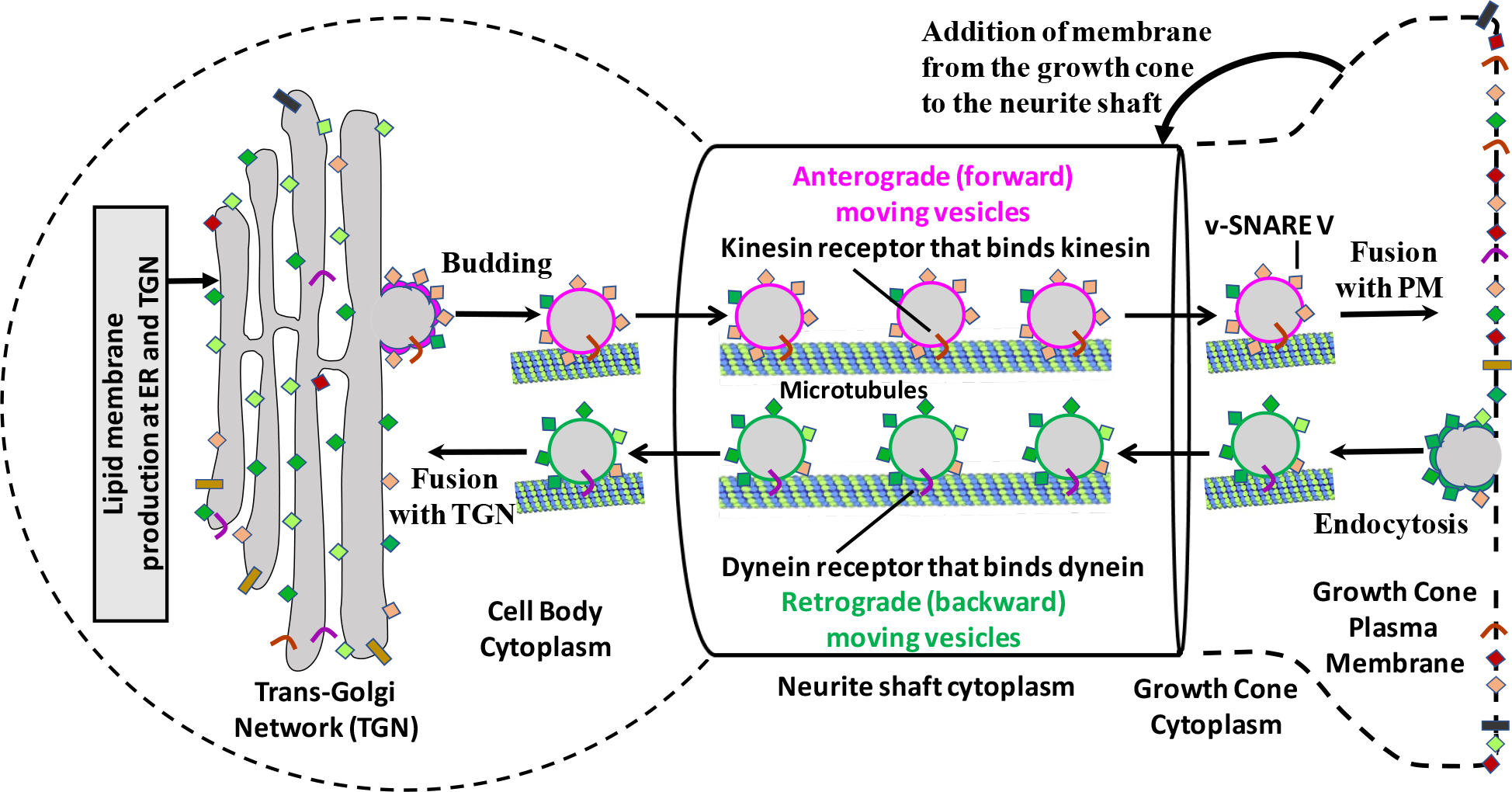
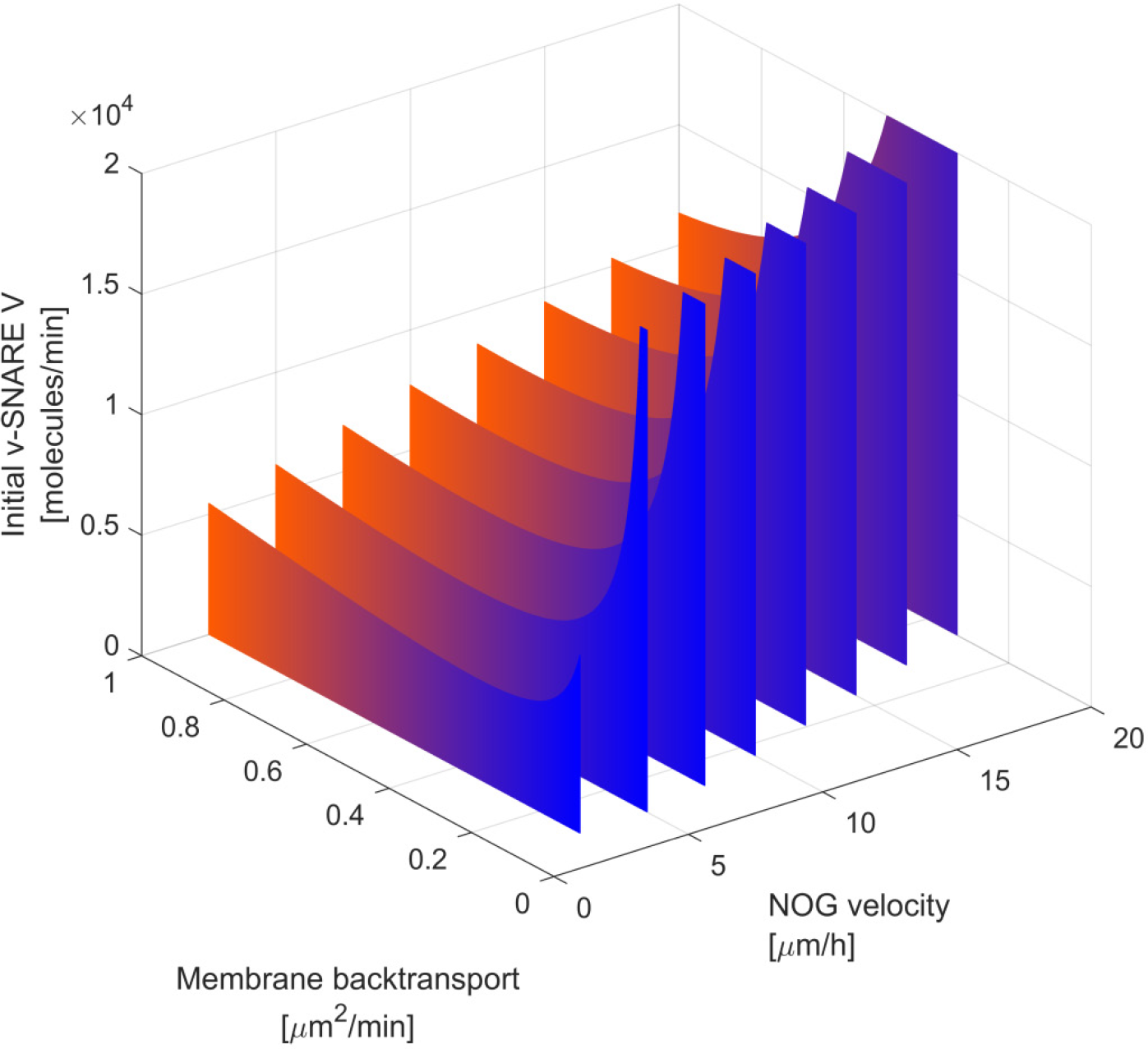
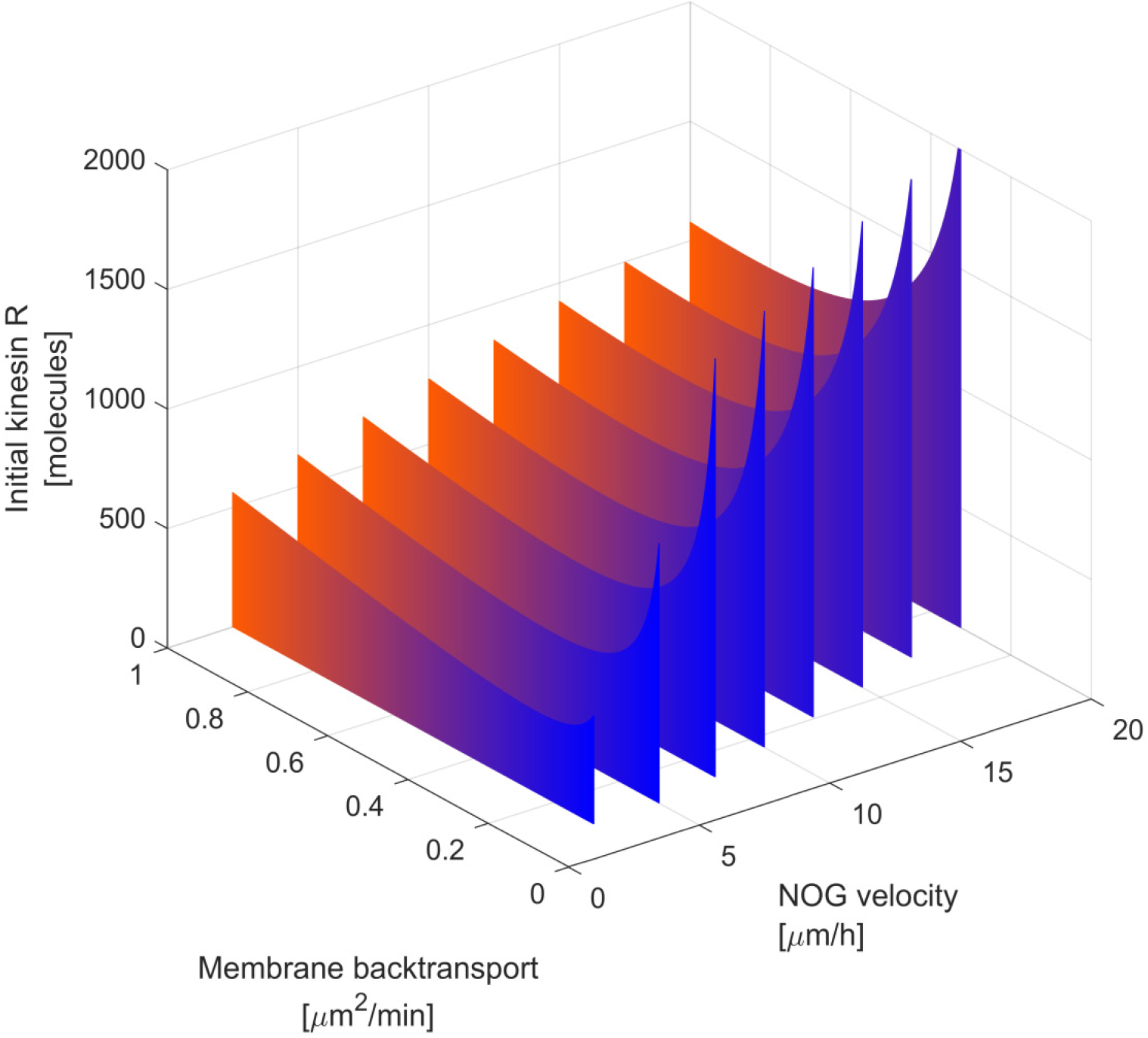
Dynamic model of vesicular transport predicts that an increase in vesicle backtransport is necessary for sufficient back transport of v-SNAREs and kinesin receptors from the growth cone to the TGN. **(A)** We used our compartmental ODE model that simulates membrane production, vesicle membrane transport to the growth cone and microtubule growth to analyze how membrane back transport influences NOG (See text for details). Part of our NOG model is an analytical solution that predicts kinetic parameter sets that allow NOG growth under a certain velocity without violation of SCP model constraints. Figure adapted from (Yadaw et al. 2017). (B/C) Our analytical solution predicts that for each velocity there is a threshold for the membrane back transport rate below which the amount of v-SNAREs (B) and kinesin receptors (C) rapidly increase. Below a second threshold NOG is no longer possible without violation of the model constraints (lack of entries past the right edge of each wall). Both thresholds shift to higher rates with increasing NOG velocity. Such constellations are the consequence of the limited number of binding spots for SNAREs and kinesin receptors on backward moving vesicles. The more forward moving vesicles bring membrane from the TGN to the growth cone, the more v-SNAREs and kinesin receptors need to be brought back from the growth cone to the TGN to be available for the next set of forward moving vesicles. Consequently, the number of backward moving vesicles has to be increased accordingly. (One of our model constraints postulates that 90% of anterograde vesicles are part of a stationary vesicle reservoir within the neurite shaft. Consequently, the elongating neurite shaft acts as a sink for antereograde vesicles and their membrane protein components. Such loss of v-SNAREs and kinesin receptors needs to be compensated by de-novo production of new proteins, increasing the total amount of v-SNAREs and kinesin receptors. (B) and (C) show the initial v-SNARE and kinesin receptor counts, before the neurite starts growing.)

Using our dynamic model, we could predict that the total number of vesicle(v)-SNAREs and vesicle kinesin receptors in the neuronal cell increases with increasing NOG velocity (Figure 4B/C). The higher the outgrowth velocity the more vesicles need to be transported to the growth cone. Consequently, higher velocities demand more v-SNAREs and kinesin receptors to equip the anterograde moving vesicles. Our model predicts that the amount of needed v-SNAREs and kinesin receptors rapidly increases, if the back transport rate is below a certain threshold. Below a second lower threshold, it is no longer possible to achieve NOG without violation of the model constraints (on simulation output past the right edges of the walls in figure 4B and 4C). Any v-SNAREs and kinesin receptors that are brought to the growth membrane need to be back transported to the TGN to be available for the next set of anterograde moving vesicles. The binding sites for v-SNAREs and kinesin receptors on a vesicle are limited. Consequently, to secure the back transport of a sufficient number of v-SNAREs and kinesin receptors at higher outgrowth velocities, more vesicles have to move backwards, increasing the total membrane back transport rate, as indicated by the differentially expressed genes (Snx3, Rab9, Rab6a, Sgsm2, Pafah1b1, Rab43) and knock down (Vps35) results. Thus using dynamical models we can demonstrate qualitative mechanistic agreement between changes in expression and dynamics of whole cell response.

## Discussion

Several previous studies in mice (Sekine et al. 2018, Tedeschi et al. 2016) and in worms (Chisholm et al. 2016) have shown that during axonal regeneration after injury, DEGs encode components of numerous cell signaling pathways including those involved in synaptic plasticity as well as SCPs involved in vesicle transport. A genome wide screen during regeneration identified a distinct set of genes including members of the SOCS family, inhibitors of signaling through Stat3. This finding is in agreement with our previous finding of an essential role for Stat3 in neurite outgrowth (Bromberg et al. 2008). Another prominent gene identified in the genome wide screen was Rab27 a protein involved in vesicle transport (Sekine et al. 2018). We started with a different approach, where our goal was to identify genes regulated by a GPCR, the Cb1 receptor, so that we could identify the SCPs involved and how they connected with one another to produce a whole cell function. So it was not entirely unexpected that these DEGs and DEPs are somewhat different from those previously reported. In looking through the SCPs and the DEGs and DEPs involved it became clear that Cb1 receptor regulation of neurite outgrowth is both deep and distributed (Figure 5). By distributed we mean that a wide range of SCPs are needed. By deep we mean that SCPs involved are often part of general cellular function and not necessarily selective for neurite outgrowth or even neuronal functions. This includes regulation of pyrimidine metabolism, mitochondrial transport and regulation of RNA stability. Regulation of these SCPs can affect many cellular responses in many different cell types and hence, it is possible that such deep regulation may be a common feature of cell state change. Future experiments in other cell types will be needed to determine if these same SCPs are involved in other cell state change events.

**Figure 5:**
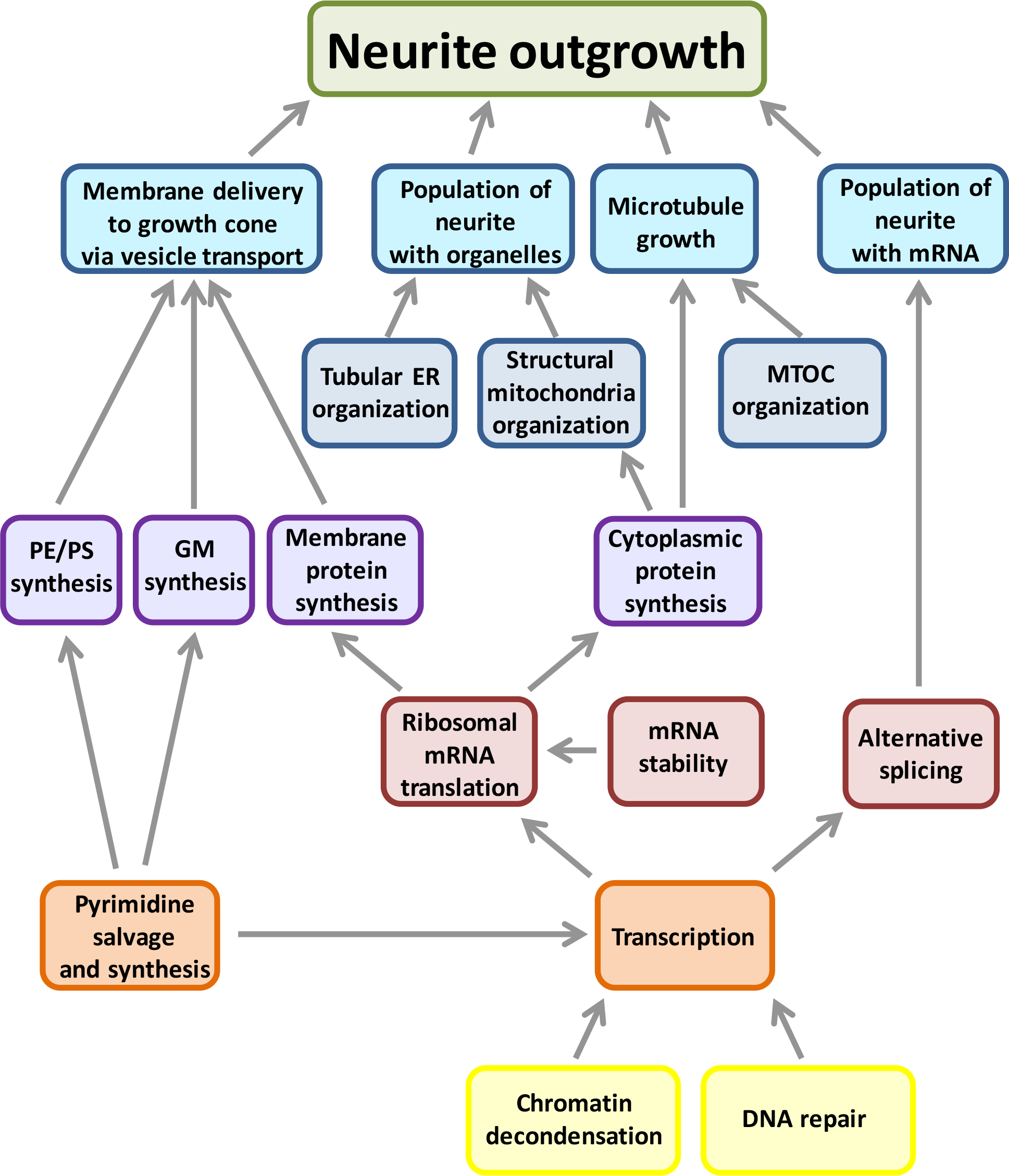
Neurite outgrowth depends on deep and distributed regulation. Multiple SCPs interact with each other to enable neurite outgrowth. The SCPs describe sub-cellular functions that can either be directly associated with neurite shaft and neurite scaffold growth, e.g. ‘Membrane lipid delivery to the growth cone’ and ‘Microtubule growth’, or describe deeper more basal sub-cellular functions, e.g. ‘Pyrimidine salvage and synthesis’. The network proposes SCP interactions that constitute the deep and distributed regulation of NOG.

In order to validate the functional role of the SCPs inferred from differentially expressed genes we used an orthogonal strategy that had three parts. First, we conducted the siRNA based knockdown studies using rat cortical neurons in primary culture. Second, we studied neurite outgrowth in microfluidic chambers where we could sever initially growing neurites and then measure neurite growth in a standardized way. Although it was not our intent to study the SCPs involved in axonal regeneration after injury, our experimental approach allows us to reasonably suggest that the SCPs we identified could play a role in axonal regeneration. Third, we did not knock down only genes that were differentially expressed, but also genes, whose expression levels did not change but were within SCPs that we computationally inferred to be relevant for neurite outgrowth. This approach allowed us to focus on SCPs rather than on individual genes. Overall the siRNA based knockdown experiments allowed us to identify a diverse set of SCPs required for neurite outgrowth. It should be noted that the magnitude of the effects are modest, both for the differentially expressed genes as well as the effect of knockdown on neurite outgrowth. This is not surprising given the wide range of SCPs involved. As documented for multifactorial diseases such as diabetes mellitus type II (Morris et al. 2012) or abdominal aortic aneurysm (Li et al. 2018), individual genes only have small to moderate effects, but interact with each other to enable the whole cell response.

For many of the deep SCPs we were also able to find previous studies that had individually implicated a role for the SCPs in neurite outgrowth. These include phospholipid biosynthesis (Carter et al. 2008) and mitochondrial functions (Cheng, Hou and Mattson 2010). Nevertheless, our findings cannot be interpreted as definitive proof that the SCPs we describe here are involved in neurite outgrowth in all conditions and in all species. Such proof will have come from future in depth small-scale experiments that analyzes the role of the SCP of interest in terms of the different genes involved. Even with this cautious interpretation we have in this study described how a receptor can trigger a whole cell response by regulating a diverse set of SCPs and the relationship between the SCPs in enabling the whole cell response - neurite outgrowth.

## Methods

### Cell culture

mRNA was prepared from 2 to 3 pooled independent experiments to achieve a sufficiently high amount of mRNA. Briefly, for each experiment 1 million Neuro2A cells were seeded on each of eight 15cm dishes and cultured for about 24h in minimum essential medium (MEM) (supplemented with 10% fetal bovine serum (FBS), 1 mM sodium pyruvate and 1% Penicillin/Streptomycin). Low cell counts ensured low confluence leaving sufficient space for neurite outgrowth. After 16h starvation in serum free MEM supplemented with 0.1% Bovine Serum Albumin (BSA), we added HU210 (dissolved in DMSO) to the media in a final concentration of 2 μM. Control cells were treated with equal volumes of drug free DMSO. Cells were incubated for 2, 4, 6 and 8h before replacement of the media by 8ml TRIZOL, followed by collection and freezing of the cell suspension.

Cell pellets for proteomic analysis were prepared from 2 independent experiments. For each experiment we plated 1 million Neuro 2A on each of 7 15cm dishes and cultured them for about 24h, followed by 1h starvation. Media conditions were the same as described above. HU210 or DMSO was added as described above and cells were incubated for 5h, 10h and 18h, before stopping of the experiment by putting the cells on ice and replacing the media with ice-cold PBS. Cells were harvest, centrifuged (500g, 10min) and cell pellets were shock frozen using liquid nitrogen. Untreated cell pellets were harvest directly after starvation.

### RNASeq - sample preparation and sequencing

RNA was prepared using the standard TRIZOL protocol (TRIzol™ Reagent, invitrogene). RNA was subjected to quality control via an Agilent Bioanalyzer. All samples had an RNA integrity score of 10. 40 μg of total RNA was used for each sample. mRNA was purified from the RNA using a Dynabeads mRNA Purification Kit and fragmented via sonication using the Covaris E210. cDNA libraries were prepared as described previously (Nagalakshmi, Waern and Snyder 2010, Quail, Swerdlow and Turner 2009, Quail et al. 2008). Fragments of 200 to 300 base pairs length were selected via gel size selection. cDNA libraries were subjected to sequencing using an Illumina HiSeq 2000 machine. Each sample was subjected to a single lane, yielding ~100 million reads per sample.

### Identification of differentially expressed genes (DEGs)

Raw reads were aligned to the mouse reference genome mm9 using Tophat 2.0.8., samtools-0.1.7 and bowtie 2.1.0. Tophat was used with the ensemble GTF file as a gene annotation reference and the option ‘no-novel-juncs’. Output BAM files were directly subjected to differential gene expression analysis using Cufflinks with the options ‘multi-read-correct’, ‘upper-quartile norm’ and ‘frag-bias-correct’ against the mm9 genome. DEGs were identified based on adjusted p-value (FDR 5%) and a minimum fold change (log_2_[(FPKM_HU210_+1) / (FPKM_DMSO_ + 1)] >= ± log_2_(1.5) or ± log_2_(1.3)).

### Proteomics – sample preparation and mass spectrometry

Collected cell pellets were lysed in extraction buffer (8 M Urea, 1% N-octyl glucoside, 100 mM TEAB (Triethylammonium bicarbonate), 1X Protease inhibitor, 1X Phosphatase Inhibitors) and clarified by centrifugation. Proteins were then quantified using BioRad Bradford Protein Assay Kit with BSA as standard. One hundred microgram of proteins from each sample was digested overnight with Trypsin at 37 C. Samples were split into two different groups that were processed independently (group 1: 2× untreated cells, 2× 10h HU210 and 2× 10h DMSO treated cells and 2× 5h HU210 treated cells, group 2: 2× untreated cells, 2× 18h HU210, 2× 18h DMSO and 2× 5h DMSO treated cells). Both groups contained an aliquot of the same untreated cell samples of both experiments. Resulted peptides from each sample were labeled with an iTRAQ reagent. After labeling, peptides were mixed and subjected to Strong cation exchange chromatography and separated into 30 fractions. Peptides in each fraction were desalted and concentrated using C18 spin column. Based upon the quantity, fractions with low peptide yield were pooled which resulted into total of 12 fractions. Peptides in these fractions were further resolved on reverse phase chromatography and then analyzed on Orbitrap Velos Mass Spectrometer.

### Identification of differentially expressed proteins

Mass spectra were searched against rat protein sequences from SwissProt protein database using Mascot, X!Tandem and Sequest search engines. Results obtained from these search engines were further combined using Scaffold 3 Q+. Total of 4336 proteins were identified and quantified with 0.5% false discovery rate (FDR) at protein level and 0.1% FDR at peptide level. Each of proteins is identified with at least 1 unique peptide having >= 95% identification probability. To allow identification of differentially expressed proteins after 5h HU210 versus 5h DMSO treatment, we normalized each protein expression value of each of the two 5h samples within each run by dividing it by the average expression value of that protein in both untreated samples of that same run.

Only proteins that were identified based on at least two different peptides were kept for further analysis. Differentially expressed proteins were determined via student’s two sided t-test under the assumption of equal variance (p-value: 0.05) and a minimum average fold change (+/− log_2_(1.1)).

### Standard and dynamic enrichment analysis

Up- and down-regulated genes and proteins were subjected to pathway enrichment analysis using Fisher’s Exact Test and the Molecular Biology of the Cell Ontology (Hansen et al. 2017) (https://github.com/SBCNY/Molecular-Biology-of-the-Cell). MBCO is a strictly cell biological ontology that was designed based on standard cell biology books such as ‘Molecular Biology of the Cell’ (Alberts et al. 2015) and populated by a combination of PubMed abstract text mining and statistical enrichment analysis. Since mitochondria SCPs are distributed over various MBCO branches (i.e. they are children of multiple level-1 SCPs such as ‘Organelle organization’, ‘Energy generation and metabolism of cellular monomers’ and ‘Transmembrane protein import and translocation’) we generated a new level-1 mitochondria SCP (‘Structural mitochondrium organization’) and defined it to be the parent of all mitochondria related level-1 and level-2 SCPs. It was populated with all genes of its children SCPs. MBCO is populated with the human gene products. We used the Jackson Laboratories mouse informatics database(Mouse Genome Informatics [MGI], http://www.informatics.jax.org) and the National Center for Biotechnology Information homologene (http://www.ncbi.nlm.nih.gov/homologene/) database to replace human by their mouse orthologes. For statistical accuracy we kept only those gene products in the MBCO that had a chance of being identified as differentially expressed (i.e. only gene products whose genes are ensemble genes on the mm9 genome or only gene products whose proteins are annotated in the SwissProt protein database). Similarly, all differentially expressed genes or proteins that are not part of the MBCO background gene product list (that contains all 13,762 gene products that were identified in at least one abstract during the MBCO text mining) were also removed.

Significantly up- or down-regulated genes with a minimum log_2_ fold change of ± log_2_(1.5) were separately subjected to dynamic enrichment analysis, using MBCO level-2 and level-3 sub-cellular processes (SCPs). MBCO contains predicted horizontal interactions between SCPs of the same level that can be ranked by their interaction strengths and complement the hierarchical vertical relationships between parent and child SCPs. Dynamic enrichment analysis uses these horizontal interactions to predict SCP networks that underlie whole cell function. Briefly, dynamic enrichment analysis identified those level-2 or level-3 SCPs that contain at least one input gene (e.g. an up-regulated gene) and combines 2 or 3 of the identified SCPs to SCP unions, if they are strongly connected. To generate level-2 SCP-networks we use the top 20% of horizontal (within the same level) SCP interactions, to generate level-3 SCP networks the top 25% of interactions. Generated SCP unions are added as new function specific SCPs to the ontology and the new function specific MBCO is used for enrichment analysis of the input gene list via Fisher’s Exact Test. Results are ranked by p-value and the top 3 level-2 or top 5 level-3 predictions (that are either SCPs or SCP-unions) are connected with each other based on all MBCO horizontal interactions to generate SCP-networks. See (Hansen et al. 2017) for details.

Significantly, up- or down-regulated genes with a minimum log_2_ fold change of ± log_2_(1.3) were subjected to standard enrichment analysis using MBCO and Fisher’s Exact Test. All SCPs with a nominal significance of at least 5% at a particular time point were considered as significant and investigated for biological relevance.

### siRNA testing in neurite outgrowth assay

All protocols comply with the IACUC at the Icahn School of Medicine at Mount Sinai. Briefly, primary cortical neurons were seeded on one side of micro fluidity chambers using Neurobasal Media supplemented with B27, L-660 glutamine and Penicillin-streptomycin (NB+++). The two sides of the chambers are separated by a 150 μm microgroove sidewall that contains slits that are too small for whole cells to pass, but large enough to allow neurite growth into the other side of the chamber. Consequently, neurites will grow across the neurite side, allowing an easy quantification of outgrowth length. 1.5h to 2h after plating we transfected the primary neurons with a pool of four different siRNAs (Dharmacon™ Accell™siRNA delivery SMART Pools). The next day transfection media was exchanged for NB+++ media that contained 25% astrocyte conditioned media. To generate astrocyte conditioned NB+++ we incubated primary astrocytes for at least 12h with NB+++, followed by steril filtration of the used NB+++. Astrocyte conditioned media contained astrocyte secreted metabolites and growth factors that support survival of primary cortical neurons. To allow the gene silencing effect to get established we continued to incubate the cells until 48h after siRNA transfection. Basal outgrowth not attributable to the siRNAs was removed by axotomy afterwards. Another 48h later (total of 96 hrs after transfection) we stopped the experiments by exchanging the media for PBS containing 4% Para formaldehyde and 4% sucrose. Cells and neurites were stained with anti-βIII tubulin (Tuj1; Covance). To quantify NOG we imaged the slides on a LSM880 confocal microscope (10× or 20×, zoom ins: 31× or 63×). For details, see (Yadaw et al. 2017). Images were obtained with the cell body side of the chamber on the left and the neurite side on the right side of the image.

### Analysis of neurite outgrowth

Before analysis of neurite outgrowth effects of the different siRNAs, we submitted all images to a computer assisted quality control step. In a small number of samples, the axotomy damaged the cell body site, so that the neurite outgrowth results were not interpretable. An axotomy that damages the cell body site, removes cell bodies from the area right next to the microgroove wall. Consequently, tubulin fluorescence intensity is less at this area and the intense line of tubulin fluorescence is retracted from the microgroove. Based on these criteria we generated a computational pipeline to automatically identify samples with damaged cell bodies. We used Image J software and the Plot Profile function to analyze the cell body site, starting at the left edge of the microgroove. This function calculates the average intensity of all pixels at a certain distance from the microgroove. All distances were converted to 0.1 μm steps, assuming a linear increase or decrease between any two measured adjacent intensity values. In case of multiple images of one micro fluidity chamber, we calculated the average intensity for each image (without double quantification of overlapping areas in the images) and generated the average intensities for the whole chamber based on the intensities of the individual images. The intention of our algorithm was to identify those samples where the intense line of tubulin labeling was too far away from the microgroove. For this, we focused on an area of up to 100 μm left of the microgrove, i.e. of an area that is 2/3 as wide as the microgrove itself. Average cell body intensities were normalized towards the highest intensity within this area of each sample. We screened for intensity peaks within this area and determined the highest peak within the first 50 μm left of the microgroove. A peak was defined as the highest point in the middle of 5 μm interval. If the normalized intensity of this peak was not at least 85%, our algorithm concluded that the intense line of tublin staining is too far away from the microgroove wall and the cell body site of this sample was labeled damaged. Computational proposals were manually confirmed or – in a few cases – corrected.

Manual investigation also identified a few samples that did not pass our quality control for reasons other than a cell body site that was damaged during axotomy (see supplementary figure 4C for rejection reasons). All samples that did not pass our quality control steps were removed from any further analyses.

To quantify the effect of gene knock down on NOG, we quantified the average fluorescence intensity at distances to the right of the microgroove wall. As described for the cell body sites, we converted all distance to distances separated by 0.1 μm (starting with 0 μm) and merged the results of multiple images of the same sample. Each experiment contained one to three controls. In case of multiple controls we calculated the mean average intensities that were used for any further analyses. To allow comparison of the results across different experiments, we normalized all average intensities of each siRNA or control of one experiment towards control average expression at 100 μm. Quantified neurite outgrowth results showed inter-experimental variation. To compare the results across different experiments, we identified those distances from the microgroove at which the control intensity had fallen to 75%, 50%, and 25%. The intensities at these distances of each siRNA treated sample were identified and compared for each gene across the different experiments. Outliers at each distance were identified for genes with at least 4 samples using Dixon’s Q-test (Confidence interval: 0.9) and removed from further analysis. One-sample two-sided t-test (equal variance) was used to determine whether the sample outgrowth was significantly different from 75%, 50% or 25%. Any gene knock down with a nominal p-value of less than 0.05 for at least one distance, was considered as significantly influencing neurite outgrowth. Similarly, any gene knock down with a nominal p-value of less than 0.05-0.1 was considered as a possible influence on neurite outgrowth.

## Acknowledgements

This research was supported by NIH grants R01GM54508 and Systems Biology Center grant P50GM071558.

## Declaration of interests

The authors declare no competing financial interests.

## Author contributions

R.I. and J.H. conceived the project. J.H. generated and analyzed RNASeq and proteomics data and predicted siRNA targets. M.J, T.L. and H.L. conducted the proteomic experiments and analyzed proteomic data. R.T., V.R. and G.J. maintained cell cultures. M.S., R.T. and V.R. tested the siRNAs and generated the microscopic images. J.H., A.Y., M.S. and G.M. analyzed the microscopic images. J.H. developed the computational pipeline for image analysis. J.H. and A.Y. conducted the dynamical models. J.H., J.G. and R.I. analyzed and organized the data and wrote the manuscript. All authors commented on the manuscript.

