## Supplementary figures for "Whole cell response to receptor stimulation involves many deep and distributed subcellular processes"

### Suppl. figure 1A

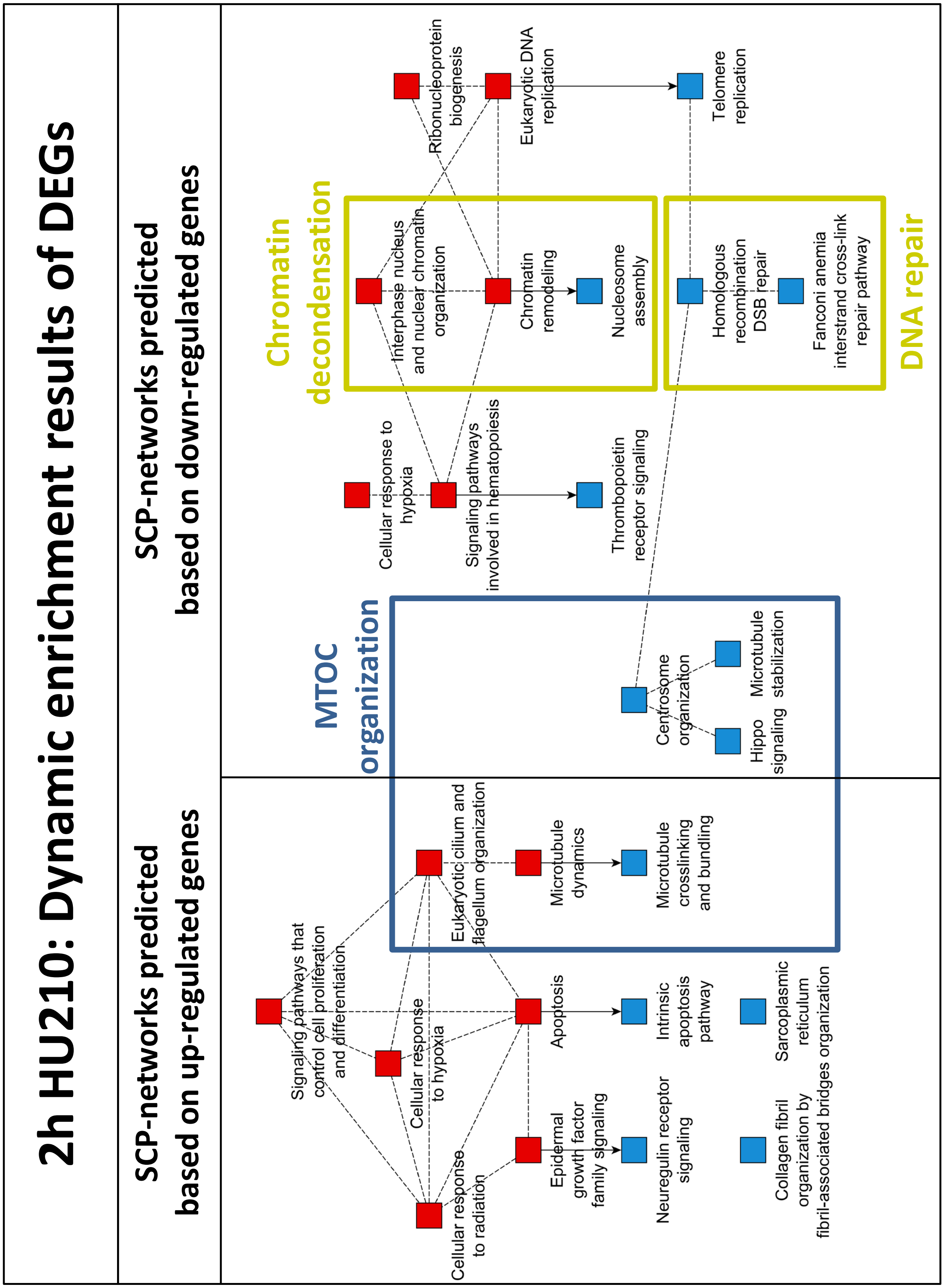

**Supplementary figure 1A: Dynamic enrichment analysis of DEGS at 2h.** Up- and down-regulated genes after 2h HU210 (minimum $log_2$(fold change) of +/- $log_2$(1.5)) were submitted to dynamic enrichment enrichment analysis using the Molecular Biology of the Cell Ontology (MBCO). Dynamic enrichment analysis considers dependencies between SCPs based on computationally predicted horizontal SCP relationships that lie within and beyond the annotated hierarchy to predict SCP-networks that describe whole cell function. Briefly, the algorithm searches for all level-2 or level-3 SCPs that contain at least one input gene and combines them to SCP-unions, if they are strongly connected. The unions are added as function-specific SCPs to the original set of SCPs and the new function-specific MBCO is used for enrichment analysis of the input genes using Fisher's Exact Test. Top three level-2 and top five level-3 predicted SCP-unions or SCPs were used to generate indicated networks. Red rectangles: level-2 SCPs, blue rectangles: level-3 SCPs. Dashed lines: horizontal SCP interactions, arrows: hierarchical annotated interactions between level-2 parent SCPs and level-3 child SCPs. MTOC: Microtubule Organization Center.

### Suppl. figure 1B

## 2h HU210:
#### Standard enrichment results of DEGs

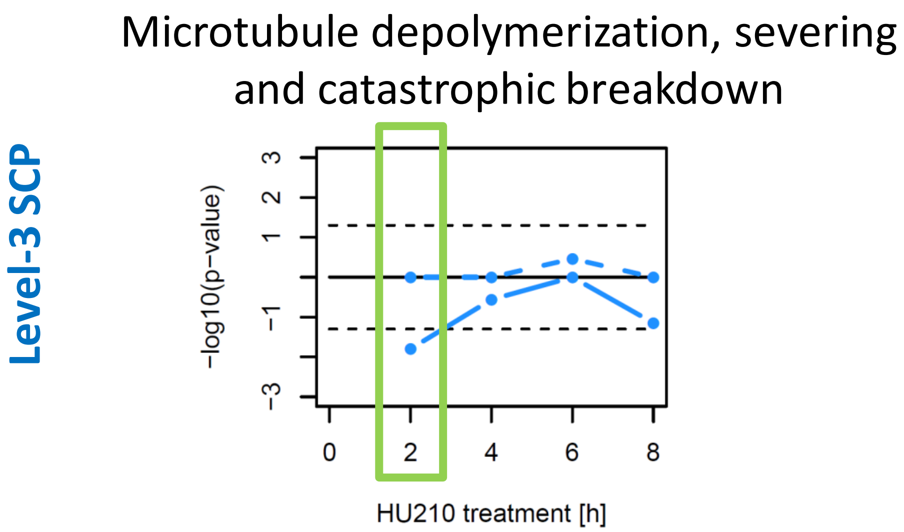


**Supplementary figure 1B: Selected SCP that was predicted by standard enrichment analysis of DEGs at 2h.** Up- and down-regulated genes of each timepoint (2h, 4h, 6h, 8h) with a minimum $log_2$(fold change) of +/- $log_2$(1.3) were submitted to standard (regular) enrichment analysis using the Molecular Biology of the Cell Ontology (MBCO) and Fisher's Exact test. Significance was defined by a maximum nominal p-value of 5%. Minus $log_{10}$(p-values) of the indicated SCP were plotted above 0, if they are based on enrichment results of up-regulated genes, and below 0, if they are based on enrichment results of down-regulated genes. If up- or down-regulated genes of at least one timepoint significantly enrich for that particular SCP, the minus log10(p-values) are connected by a solid line and by a dashed line otherwise. Horizontal black solid line indicates minus $log_{10}$(p-value) of 0 (p-value=1), black dashed line indicates nominal significance threshold (minus $log_{10}$(0.05)). Green rectangle shows the minus $log_{10}$(p-values) of the current timepoint. See supplementary figure 2 for all predicted SCPs.

### Suppl. figure 1C

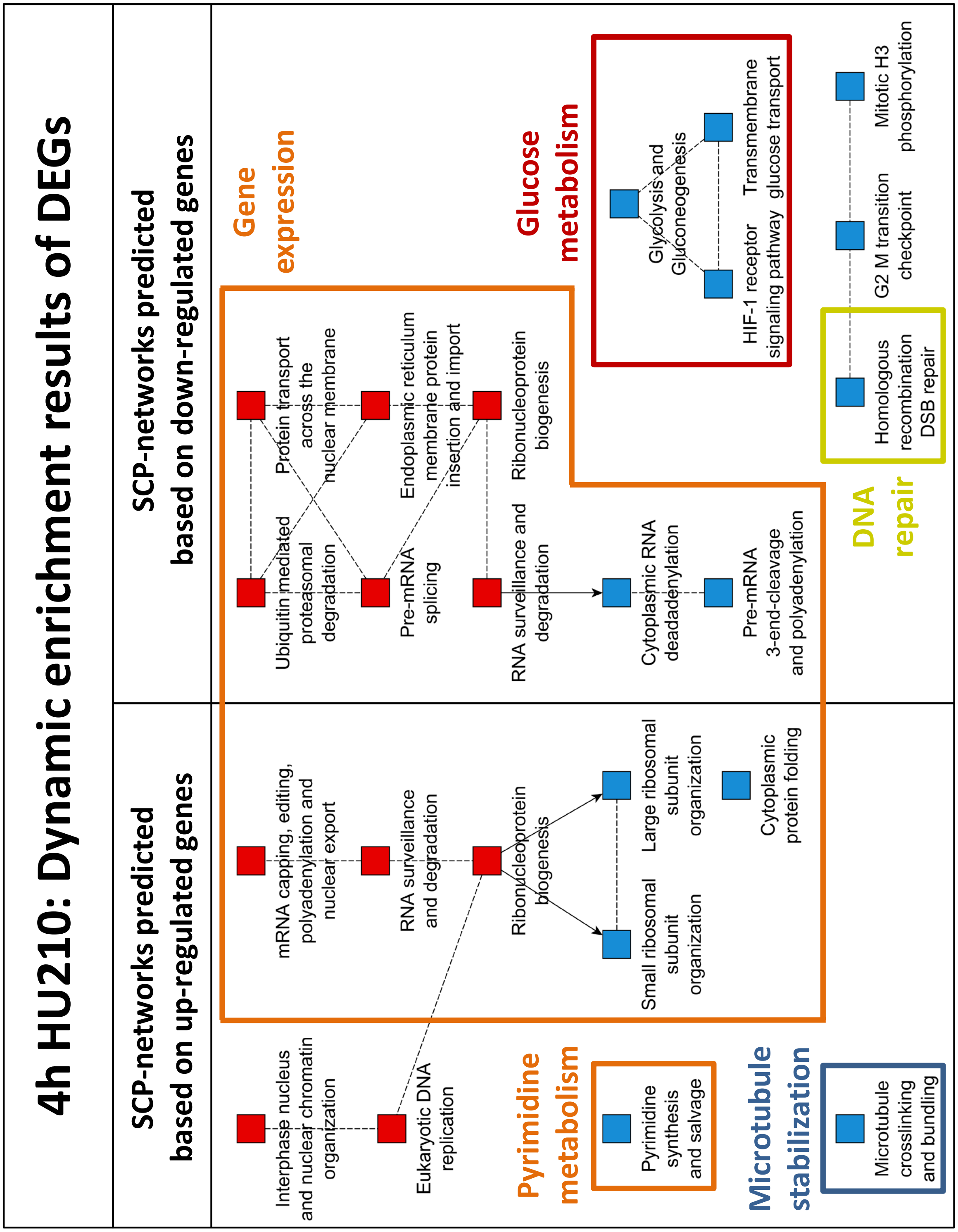


**Supplementary figure 1C: Dynamic enrichment analysis of DEGs at 4h.**

### Suppl. figure 1D

## 4h HU210:
#### Standard enrichment results of DEGs

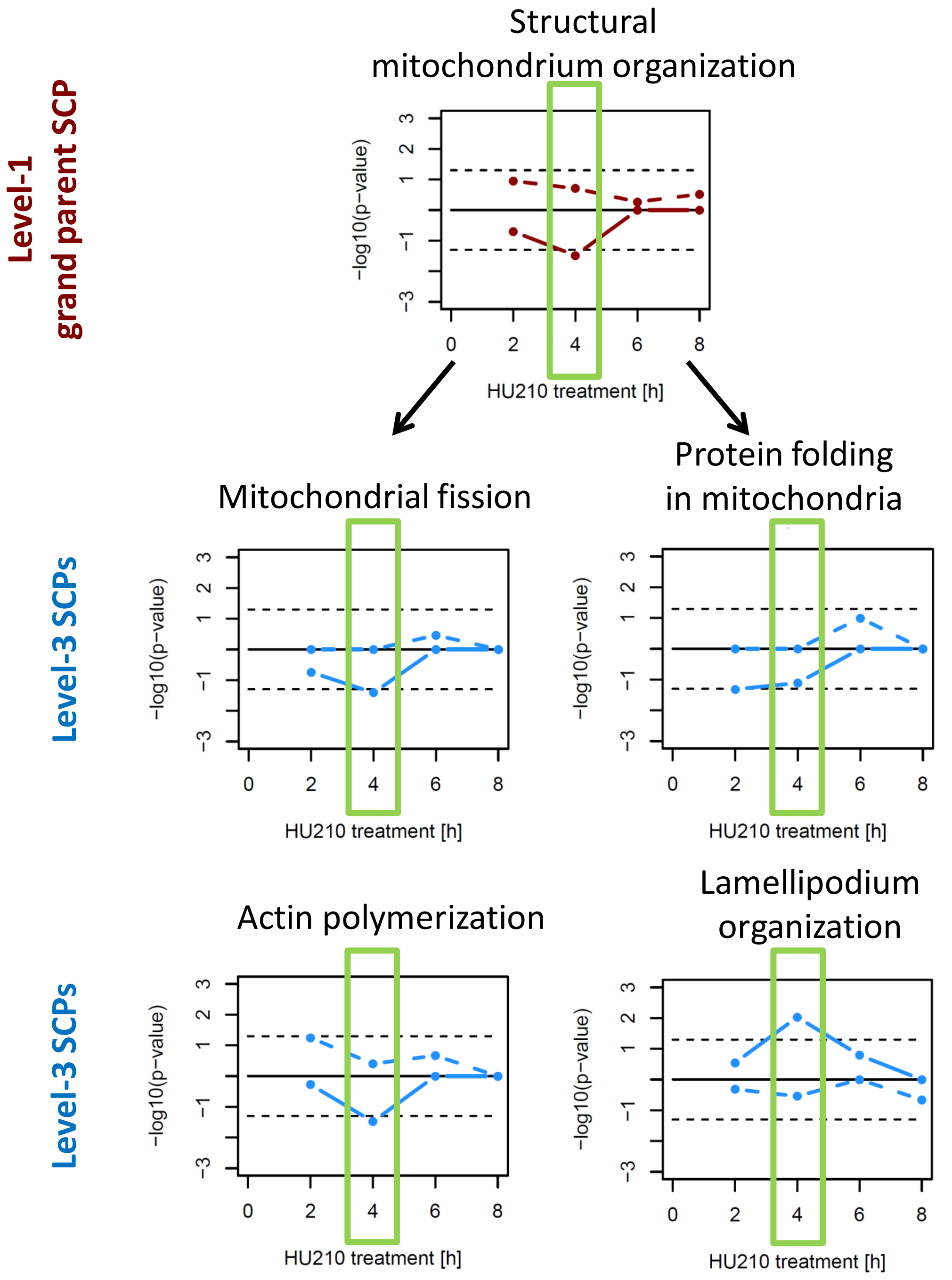


**Supplementary figure 1D: Selected SCPs that were predicted by standard enrichment analysis of DEGs at 4h.**

### Suppl. figure 1E

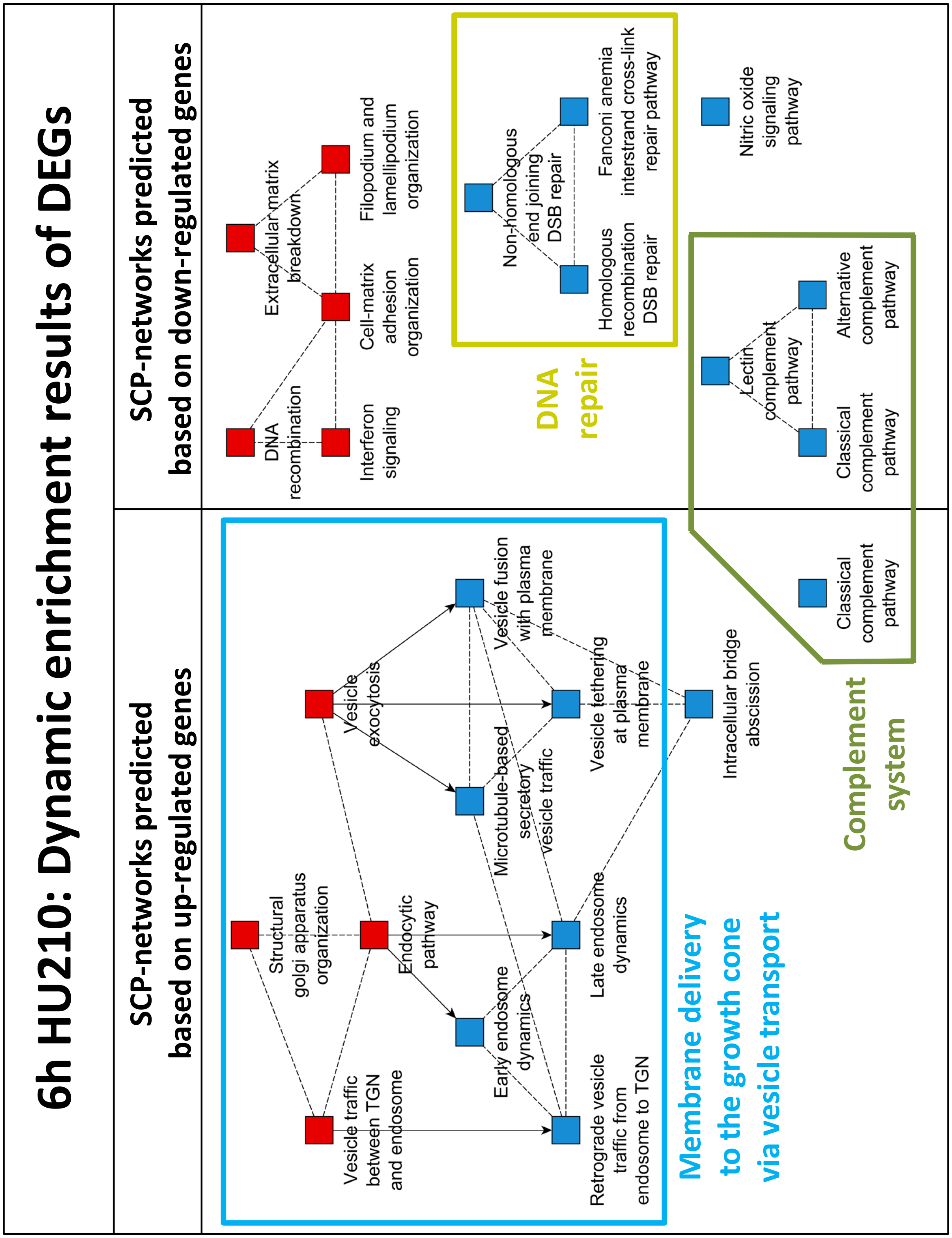


**Supplementary figure 1E: Dynamic enrichment analysis of DEGs at 6h.**

### Suppl. figure 1F

## 6h HU210:
#### Standard enrichment results of DEGs

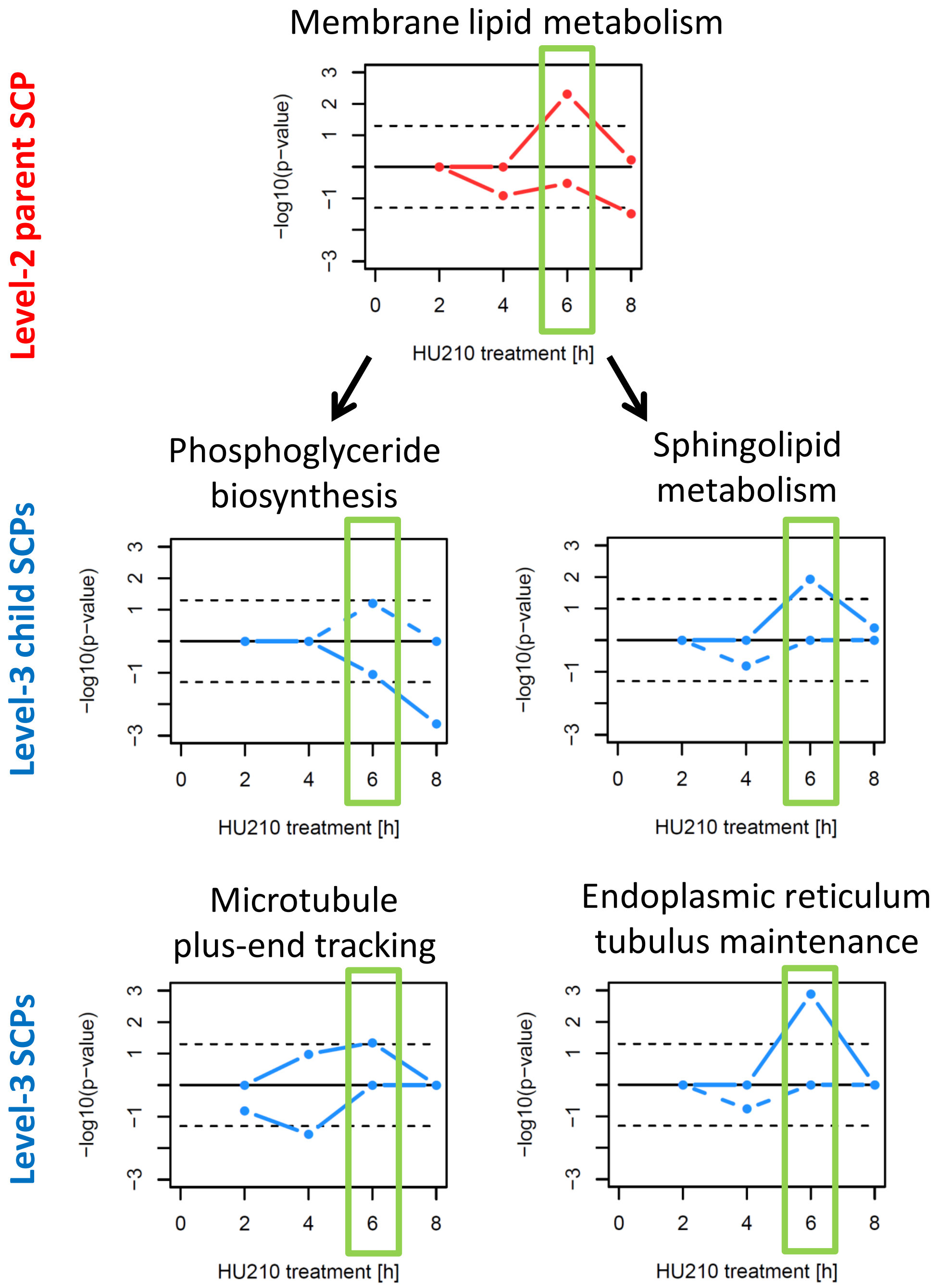


**Supplementary figure 1F: Selected SCPs that were predicted by standard enrichment analysis of DEGs at 6h.**

### Suppl. figure 1G

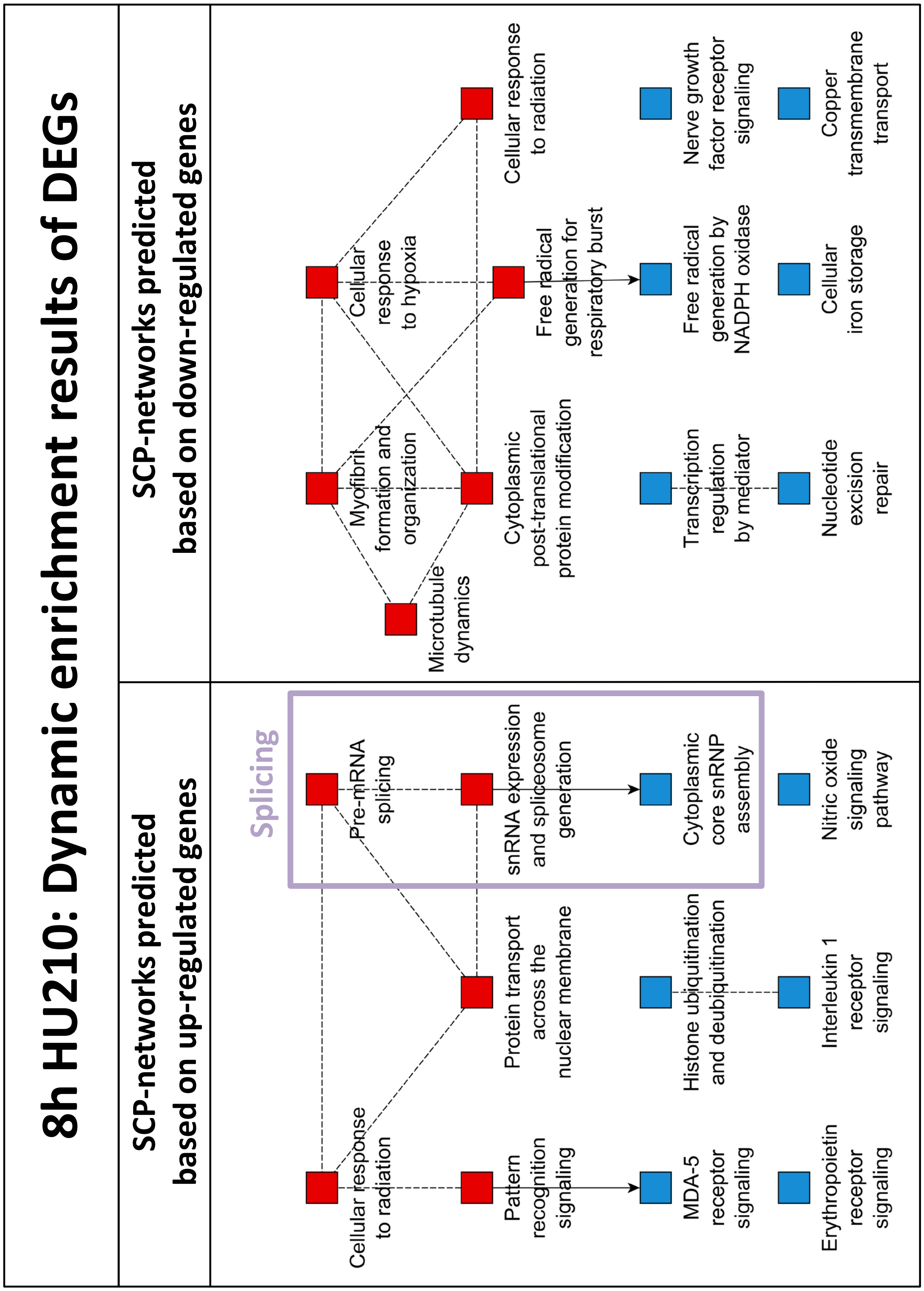


**Supplementary figure 1G: Dynamic enrichment analysis of DEGs at 8h.**

### Suppl. figure 1H

## 8h HU210:
#### Standard enrichment results of DEGs

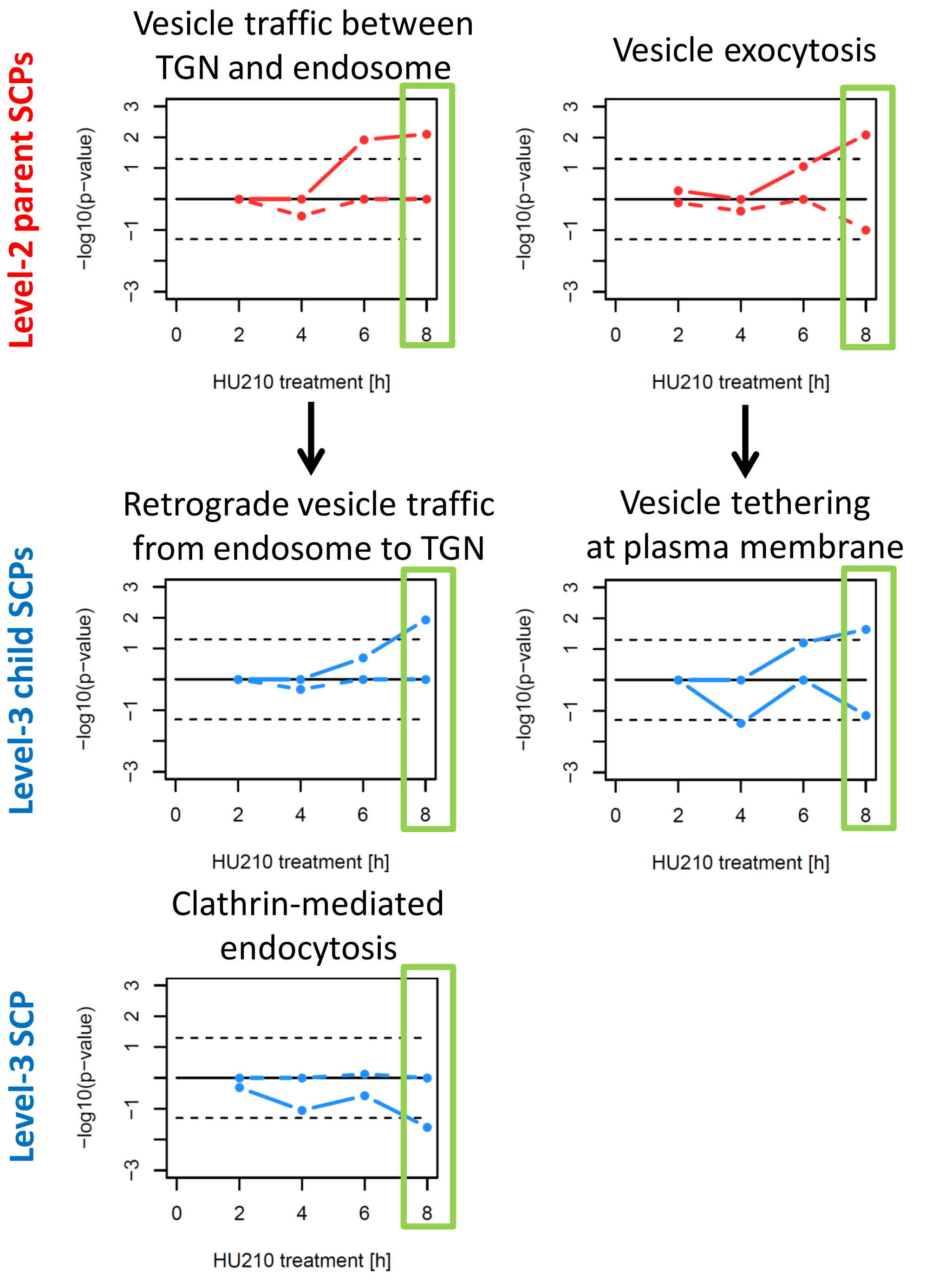


**Supplementary figure 1H: Selected SCPs that were predicted by standard enrichment analysis of DEGs at 8h.**

### Suppl. figure 1I

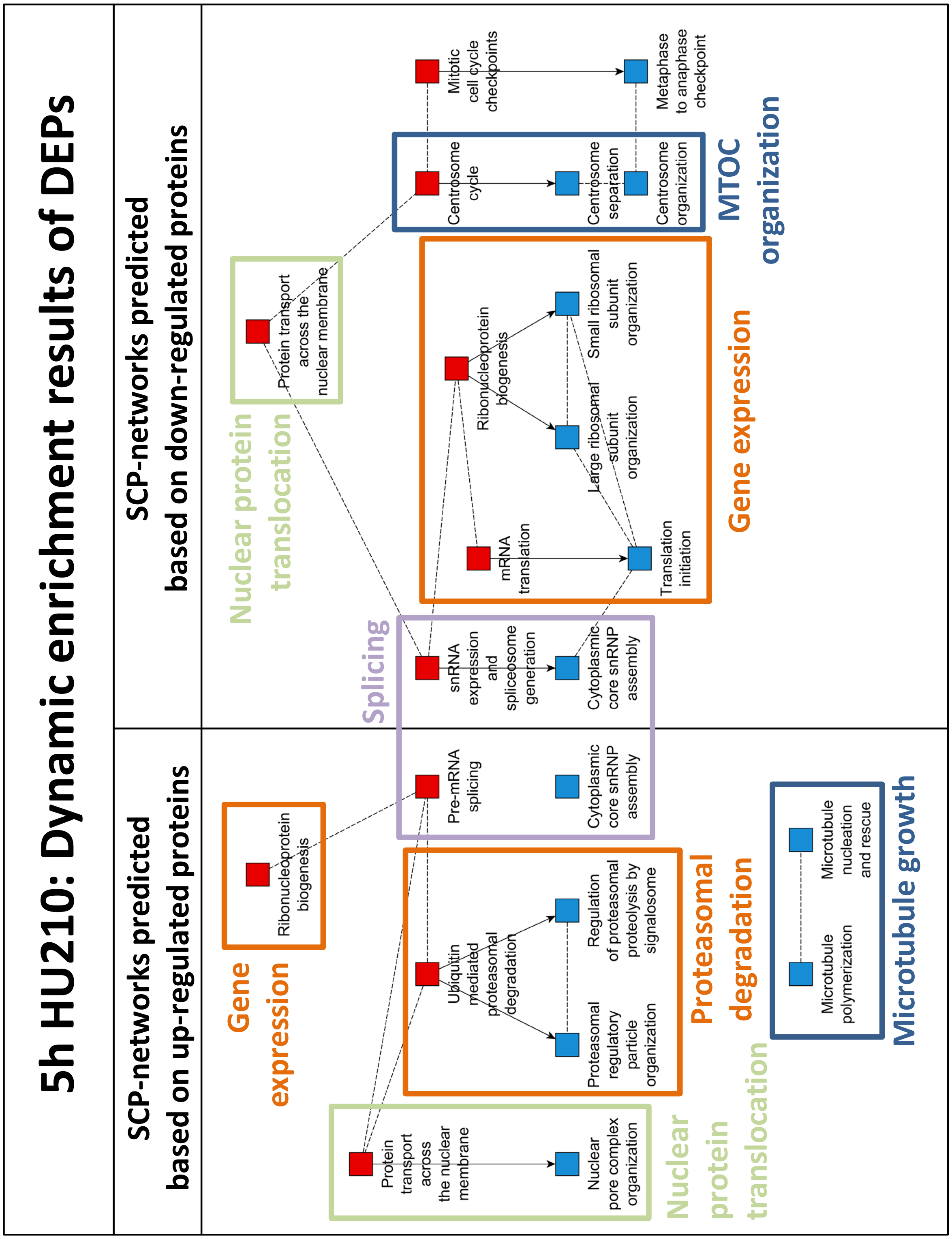


**Suppl. figure 1I: Dynamic enrichment analysis of differentially expressed proteins (DEPs) at 5h.**

### <u>Suppl. figure 1J</u>

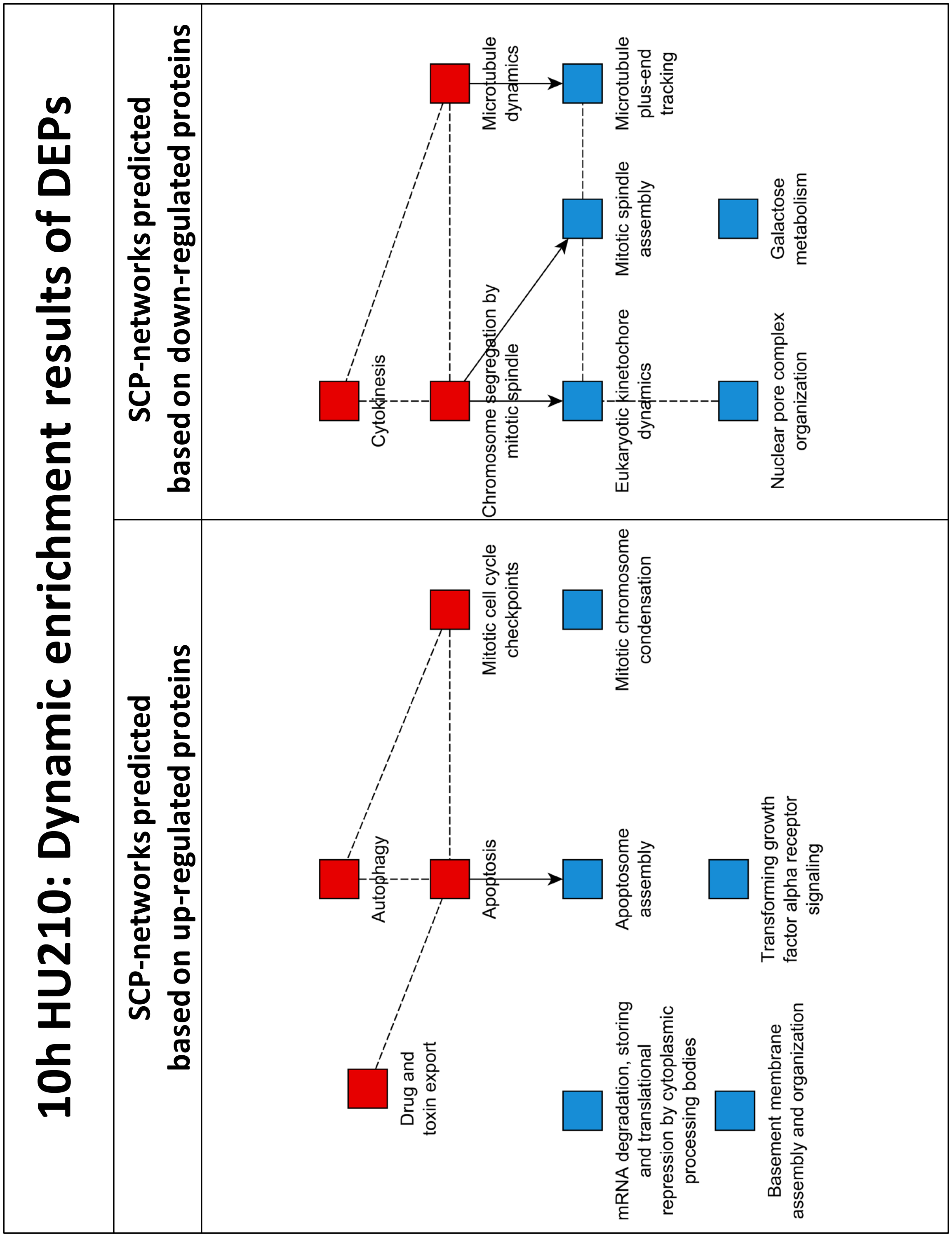


**Suppl. figure 1J: Dynamic enrichment analysis of differentially expressed proteins (DEPs) at 10h.**

### Suppl. figure 1K

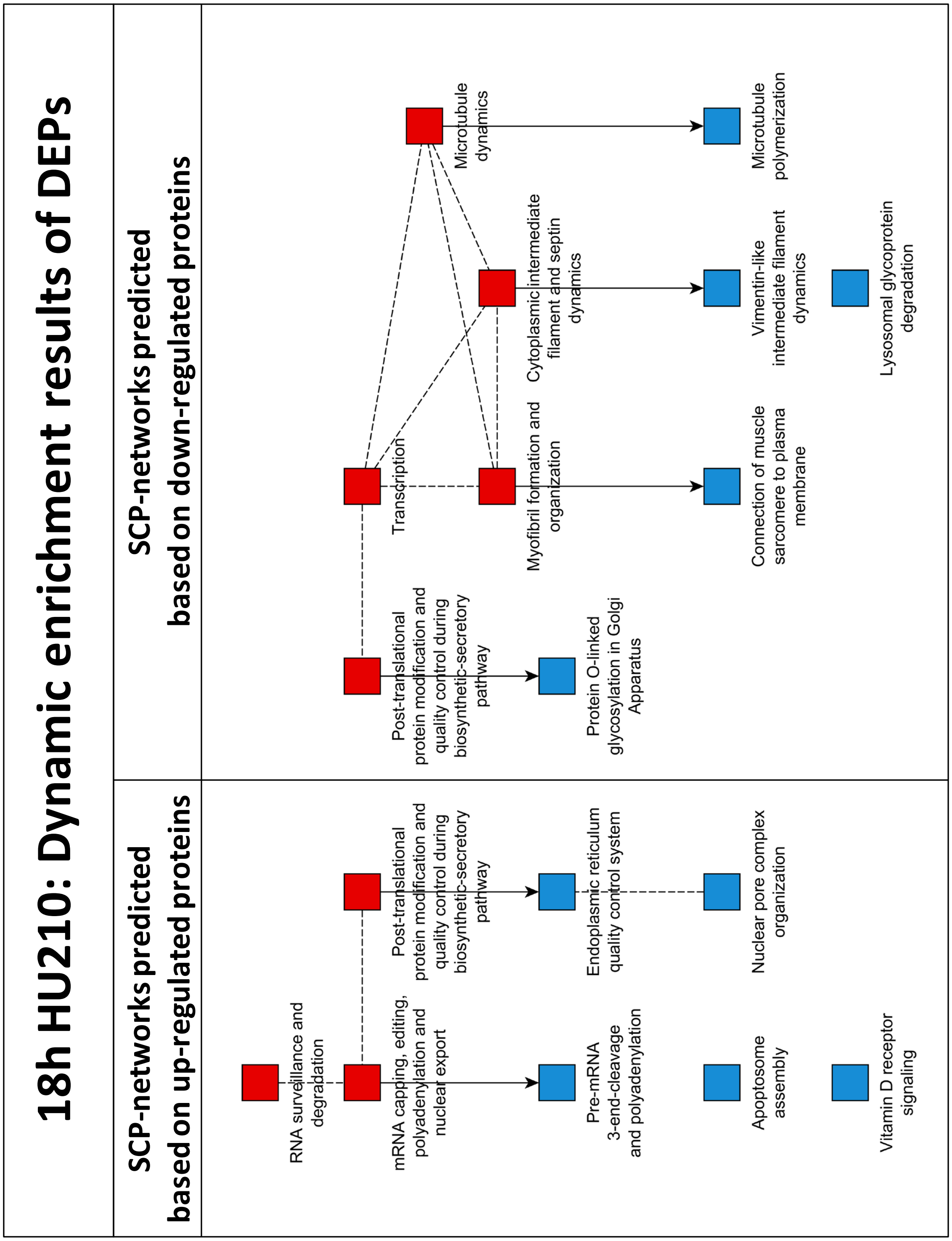


**Suppl. figure 1K: Dynamic enrichment analysis of differentially expressed proteins (DEPs) at 18h.**

### Suppl. figure 2

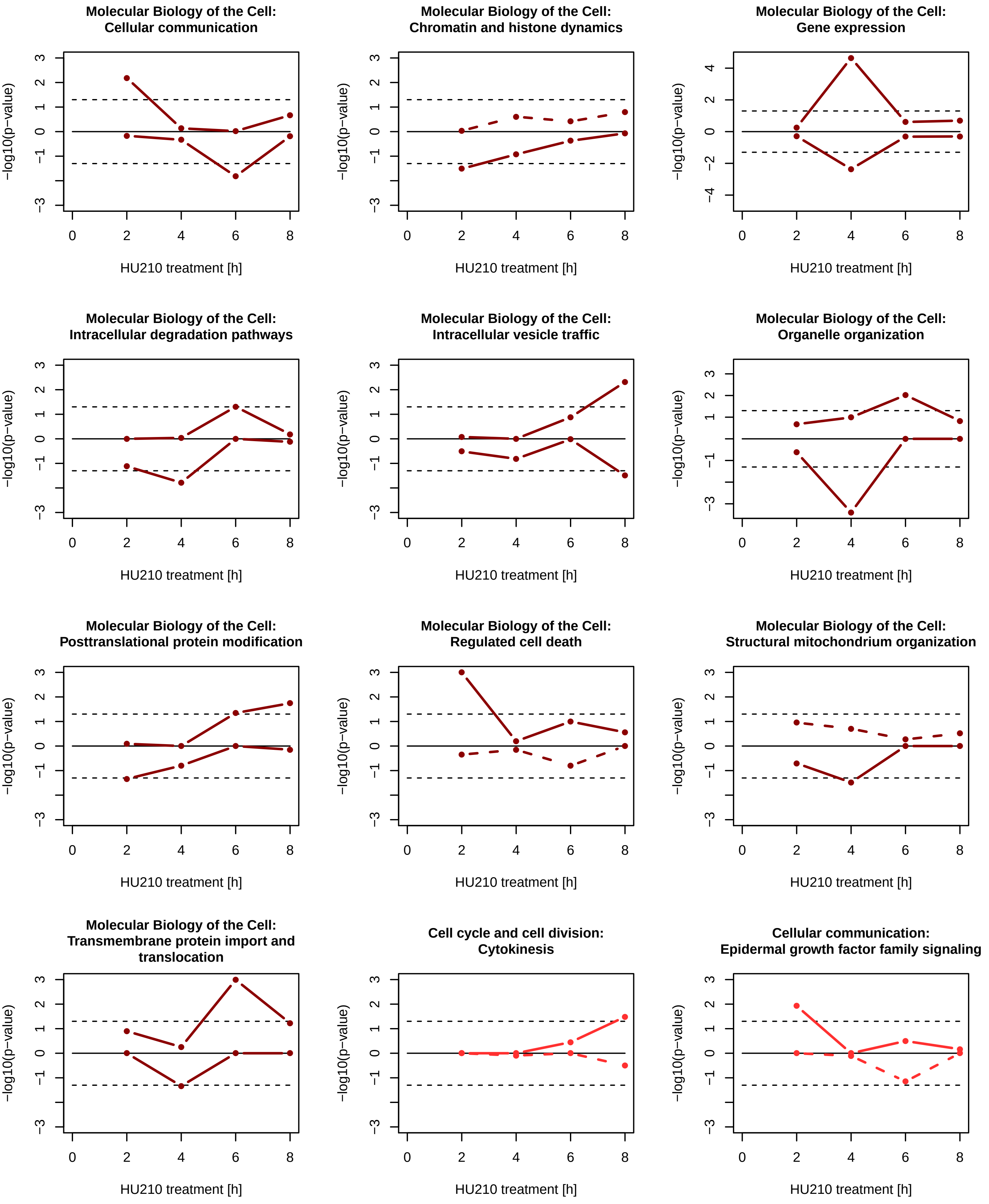

### Suppl. figure 2

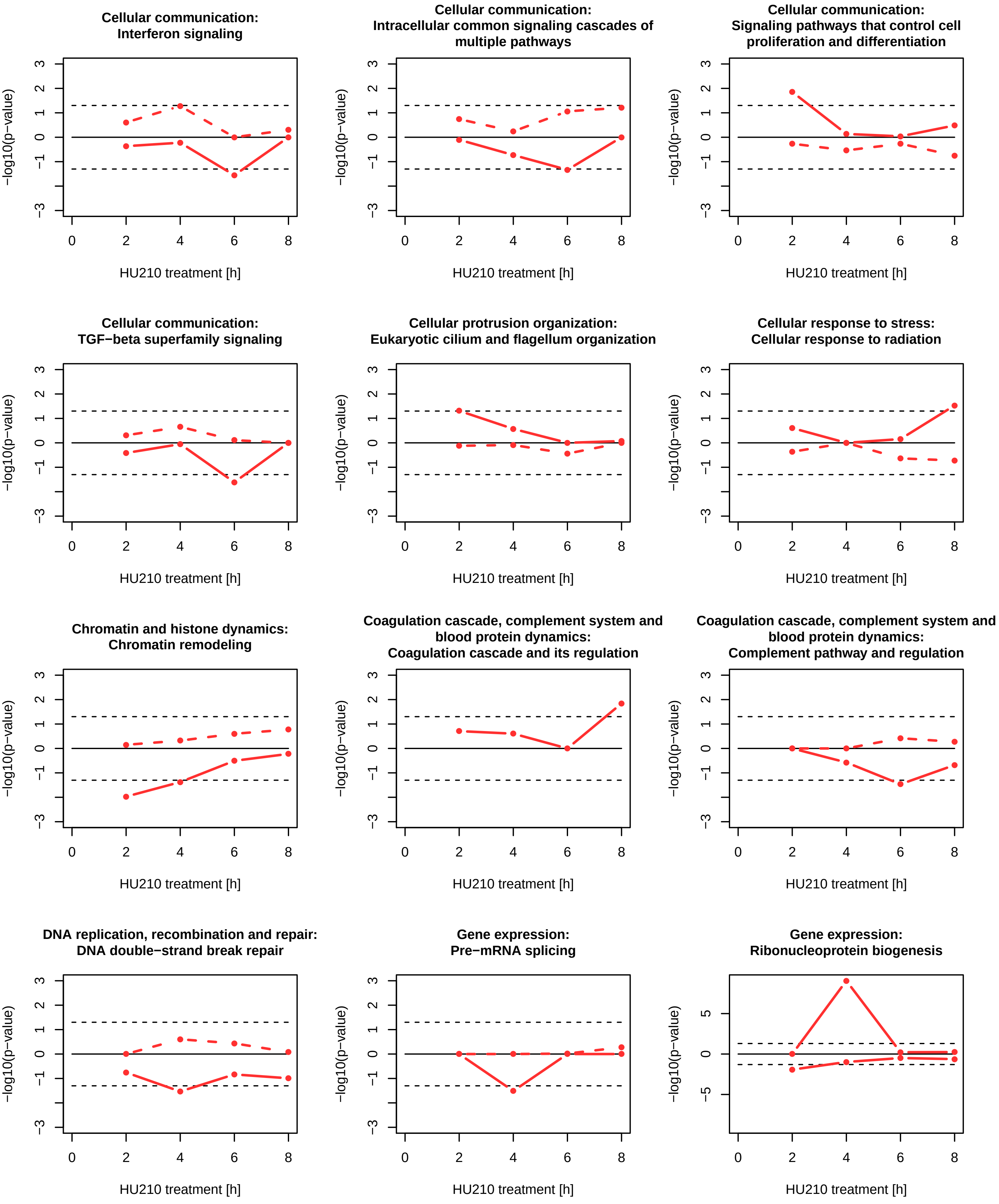

### Suppl. figure 2

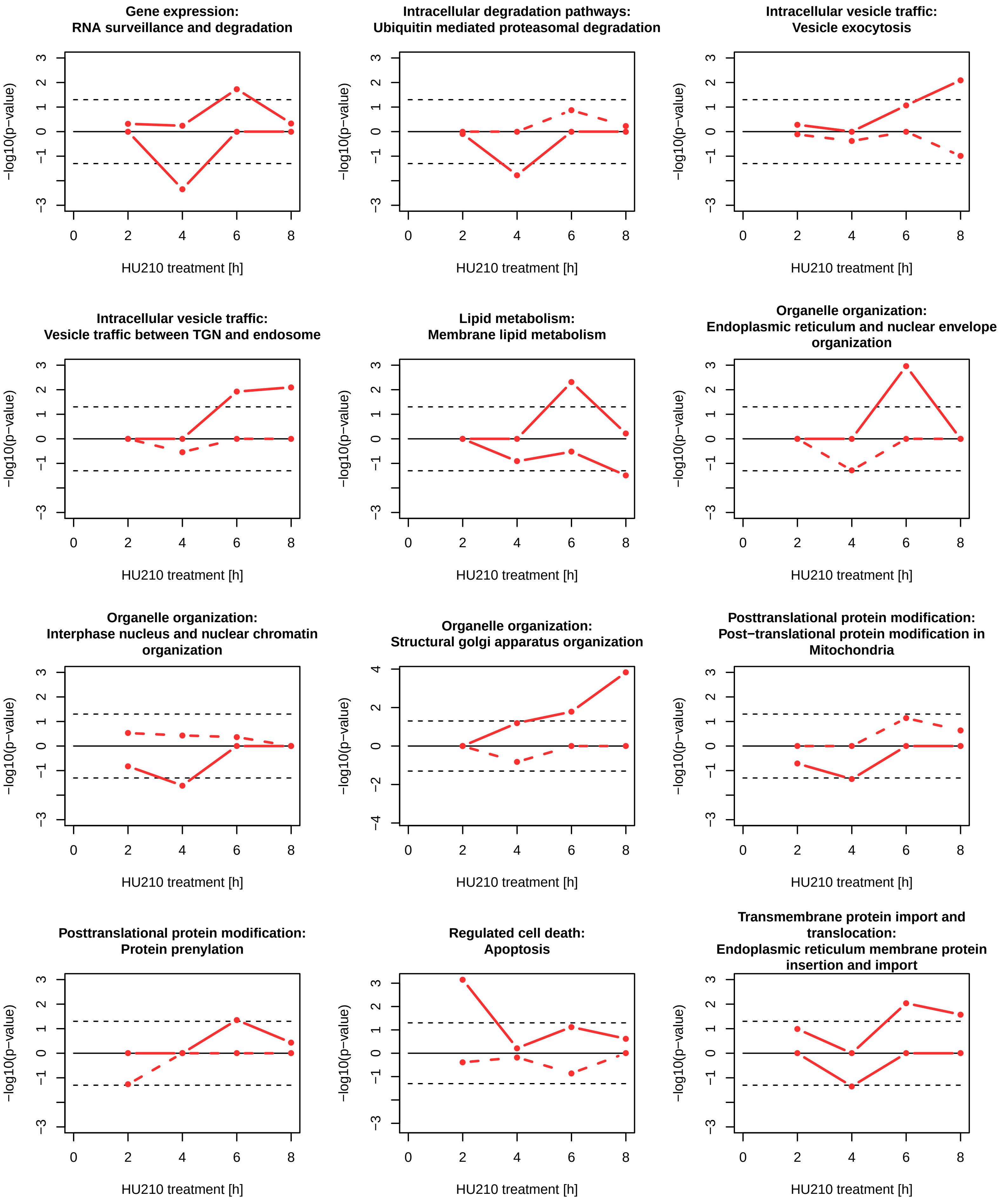

### Suppl. figure 2

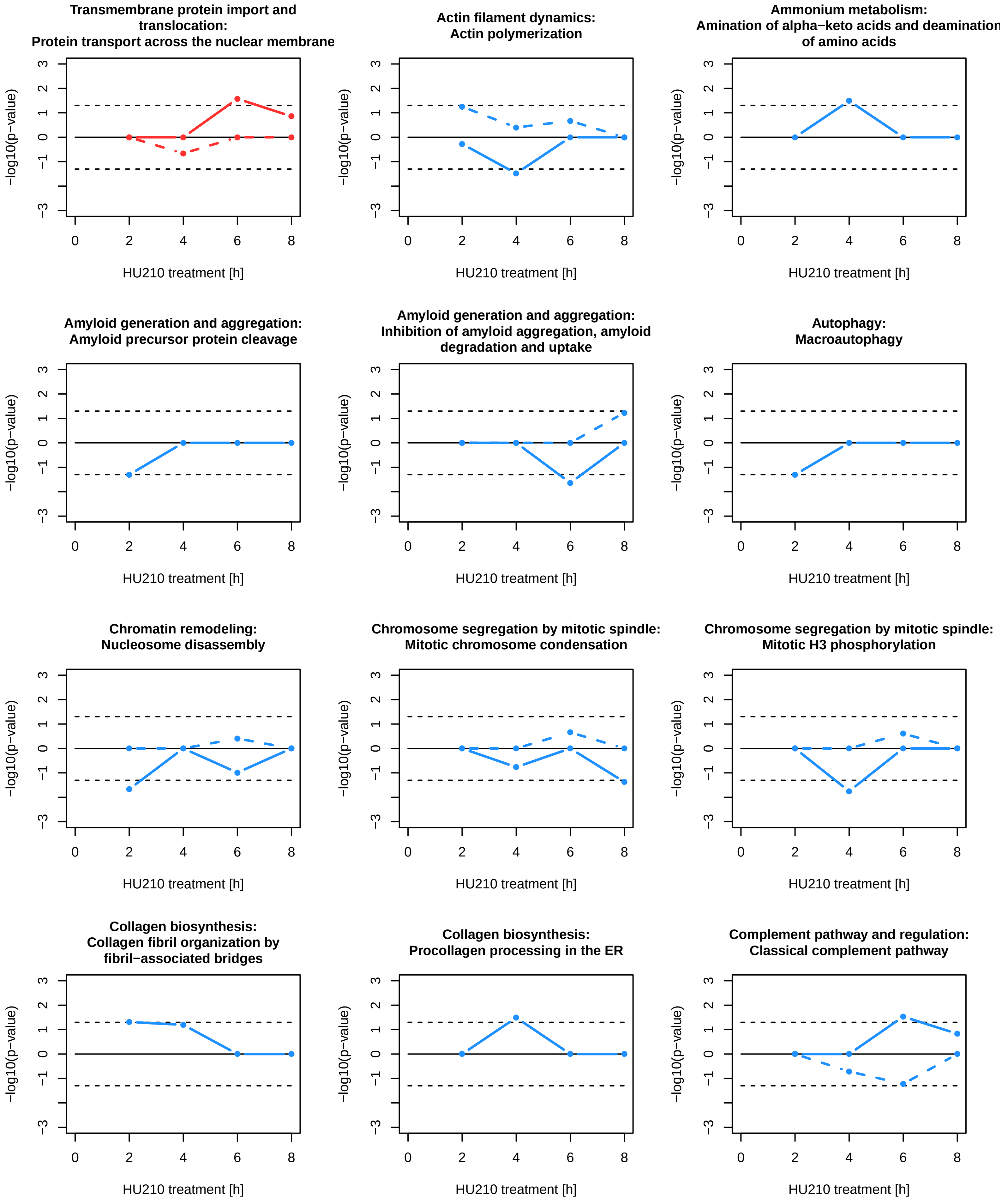

### Suppl. figure 2

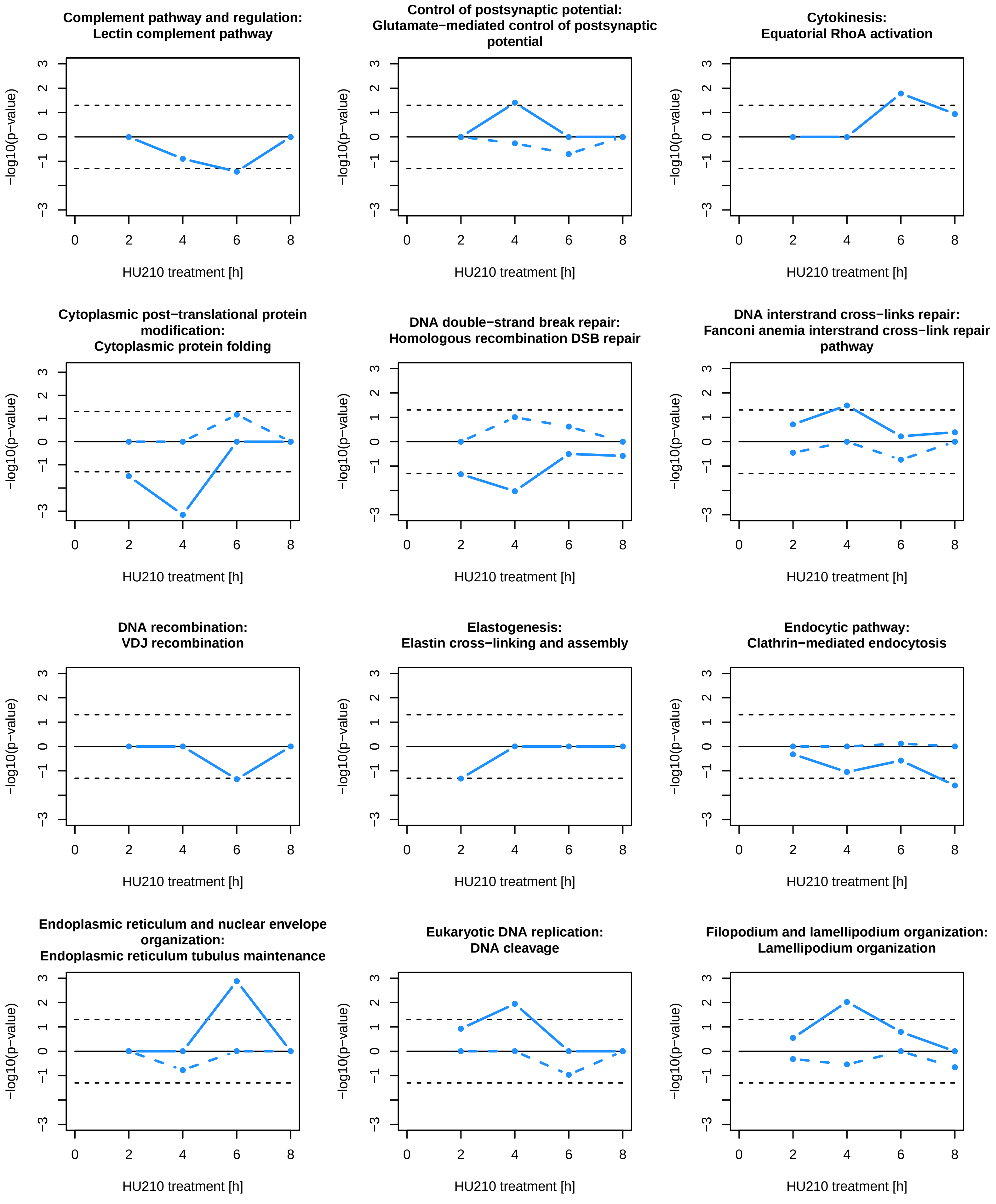

### Suppl. figure 2

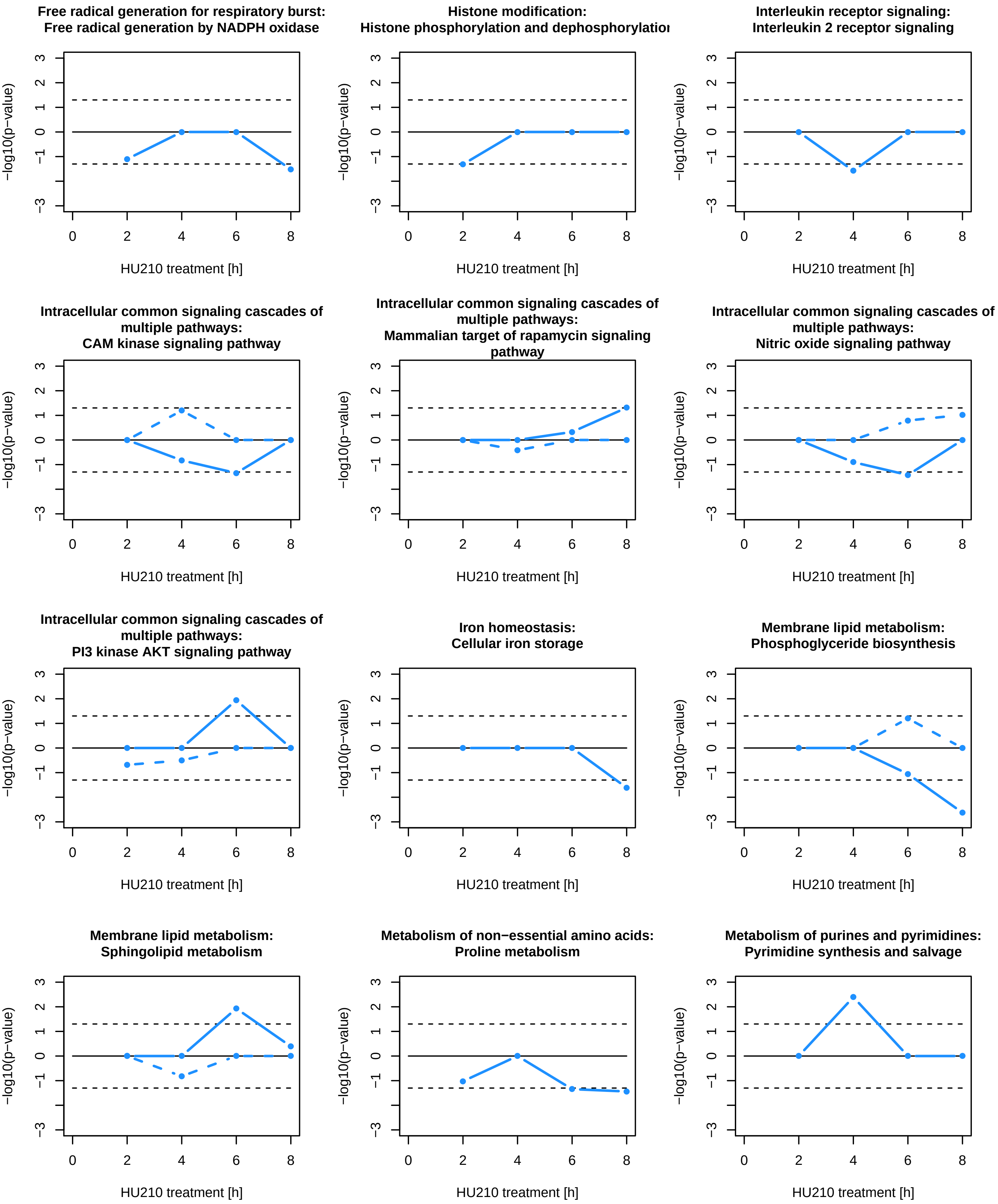

### Suppl. figure 2

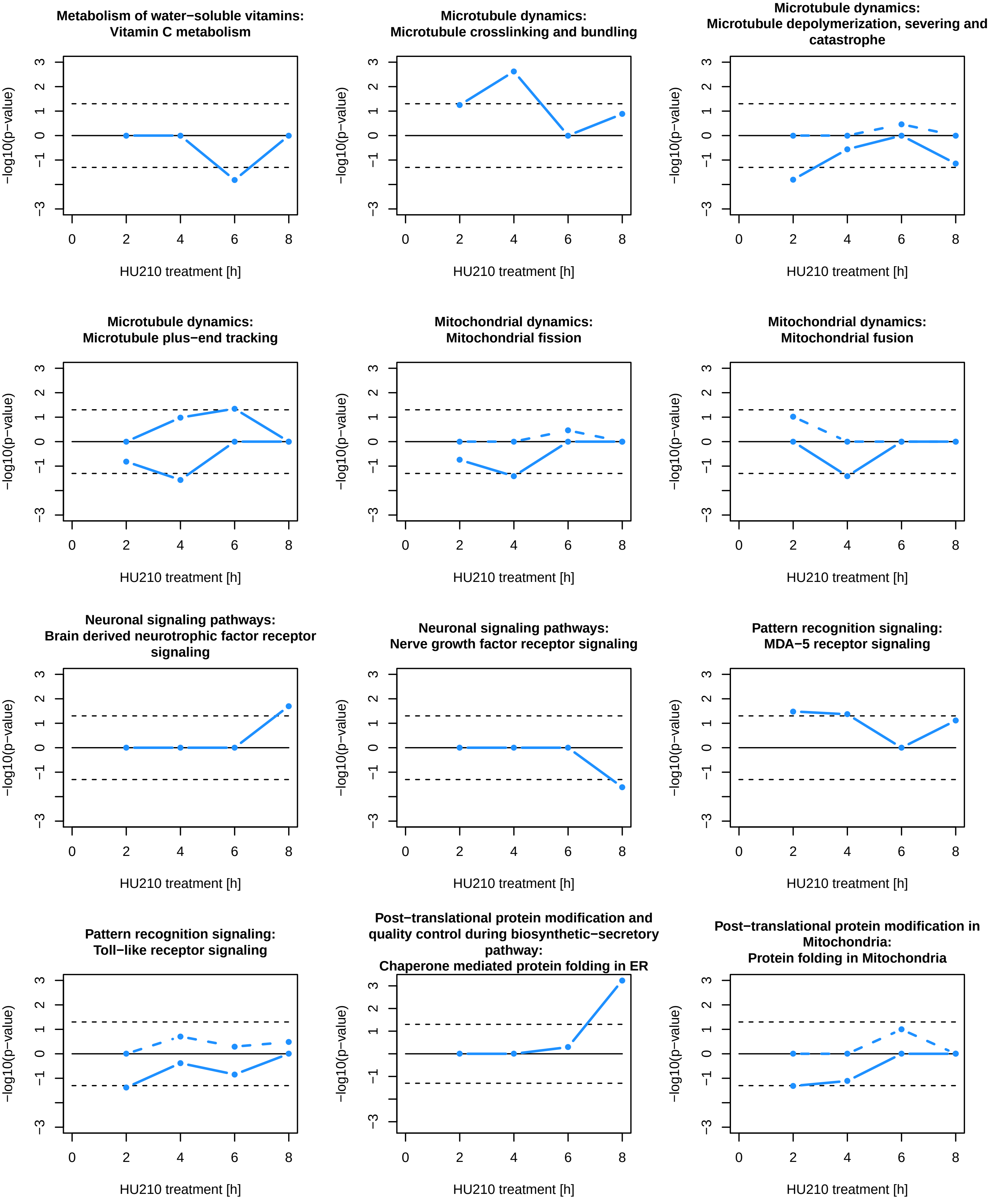

### Suppl. figure 2

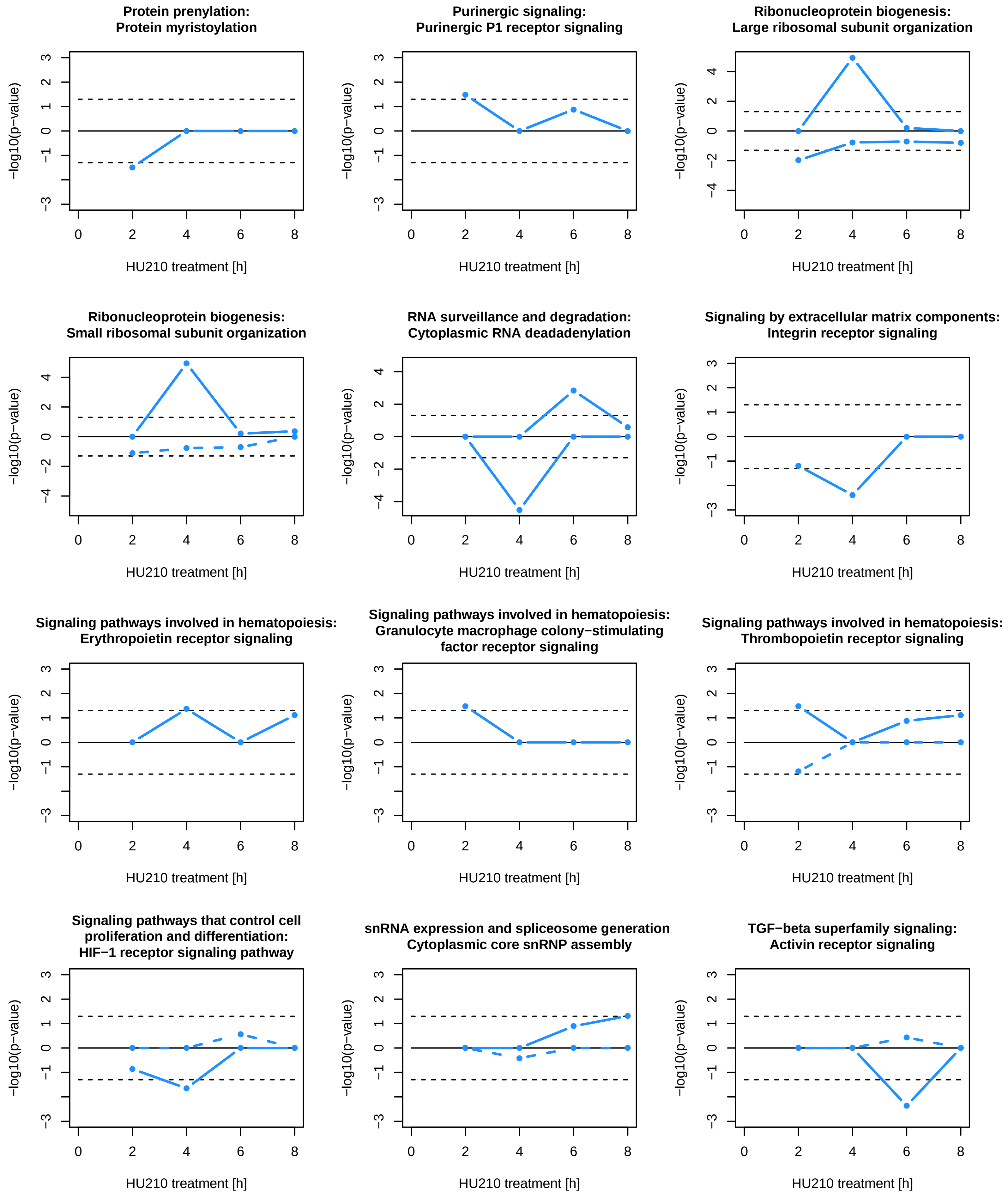

### Suppl. figure 2

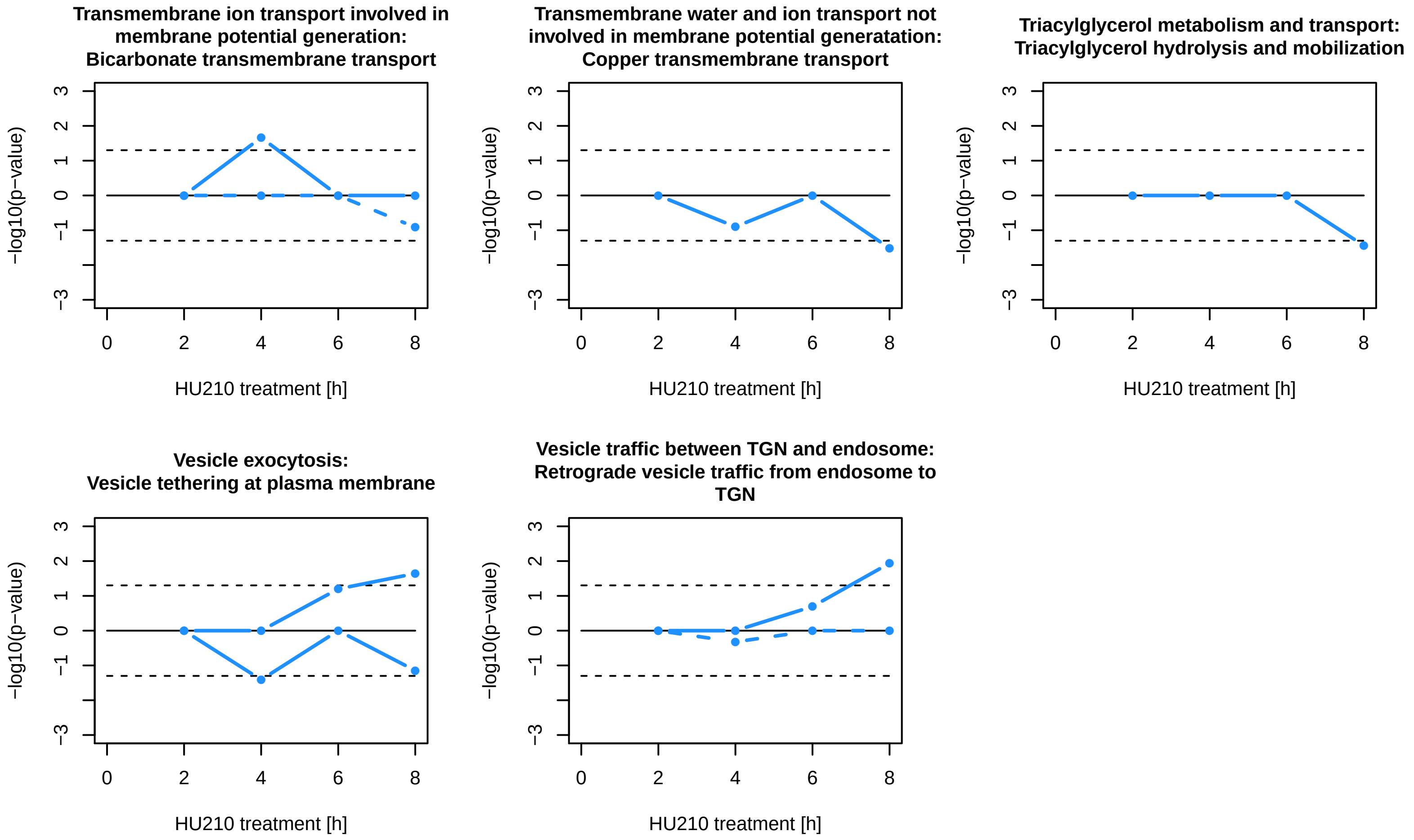

**Suppl. figure 2: Timeline of all SCPs that were predicted based on standard enrichment analysis.** First headline: Parent SCP of predicted SCP, Second headline: predicted SCP. Dark red/light red/blue: predicted SCP is level-1/level-2/level-3 SCP. For details see supplementary figure 2B.

### Suppl. figure 3

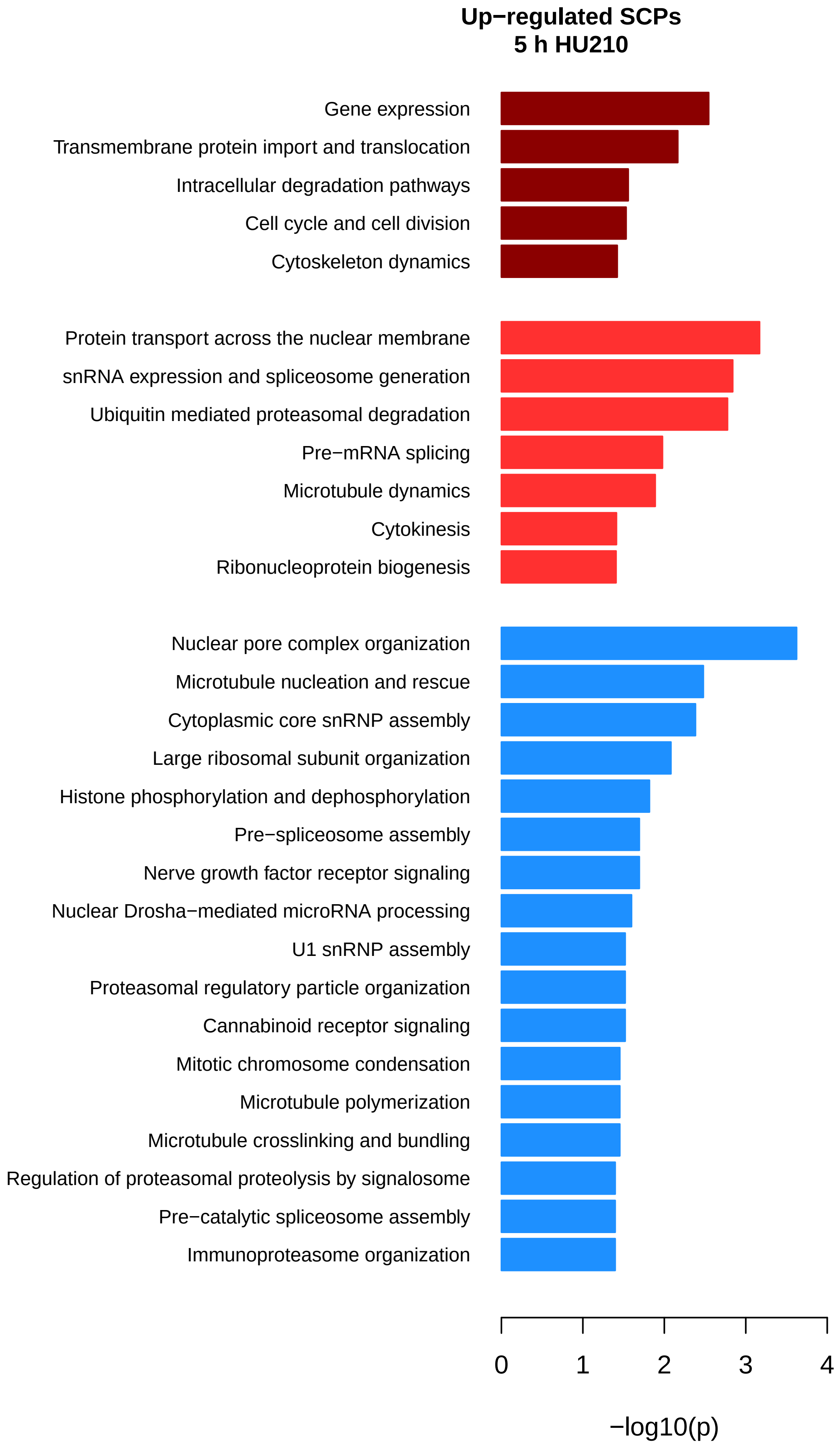

### Suppl. figure 3

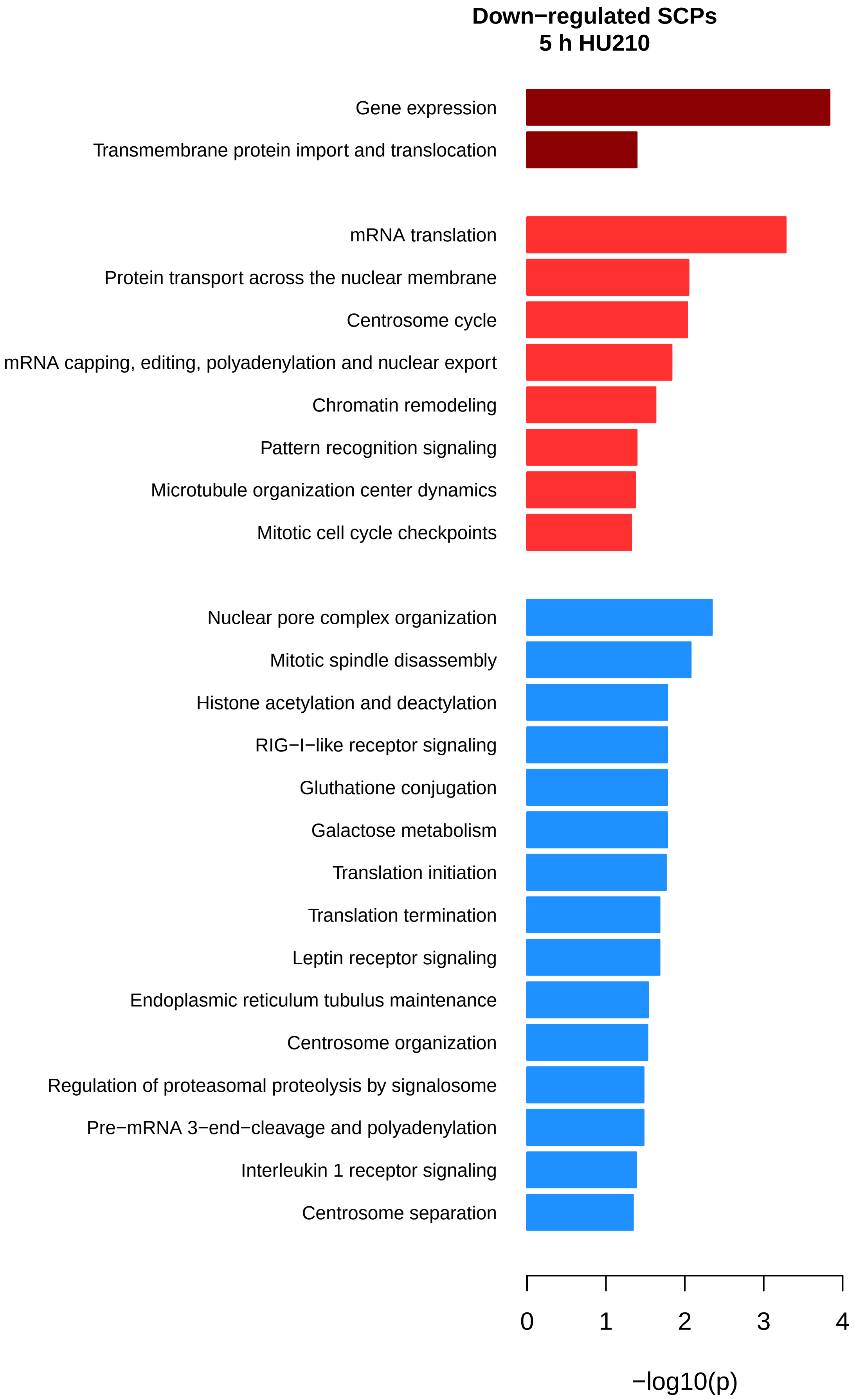

### Suppl. figure 3

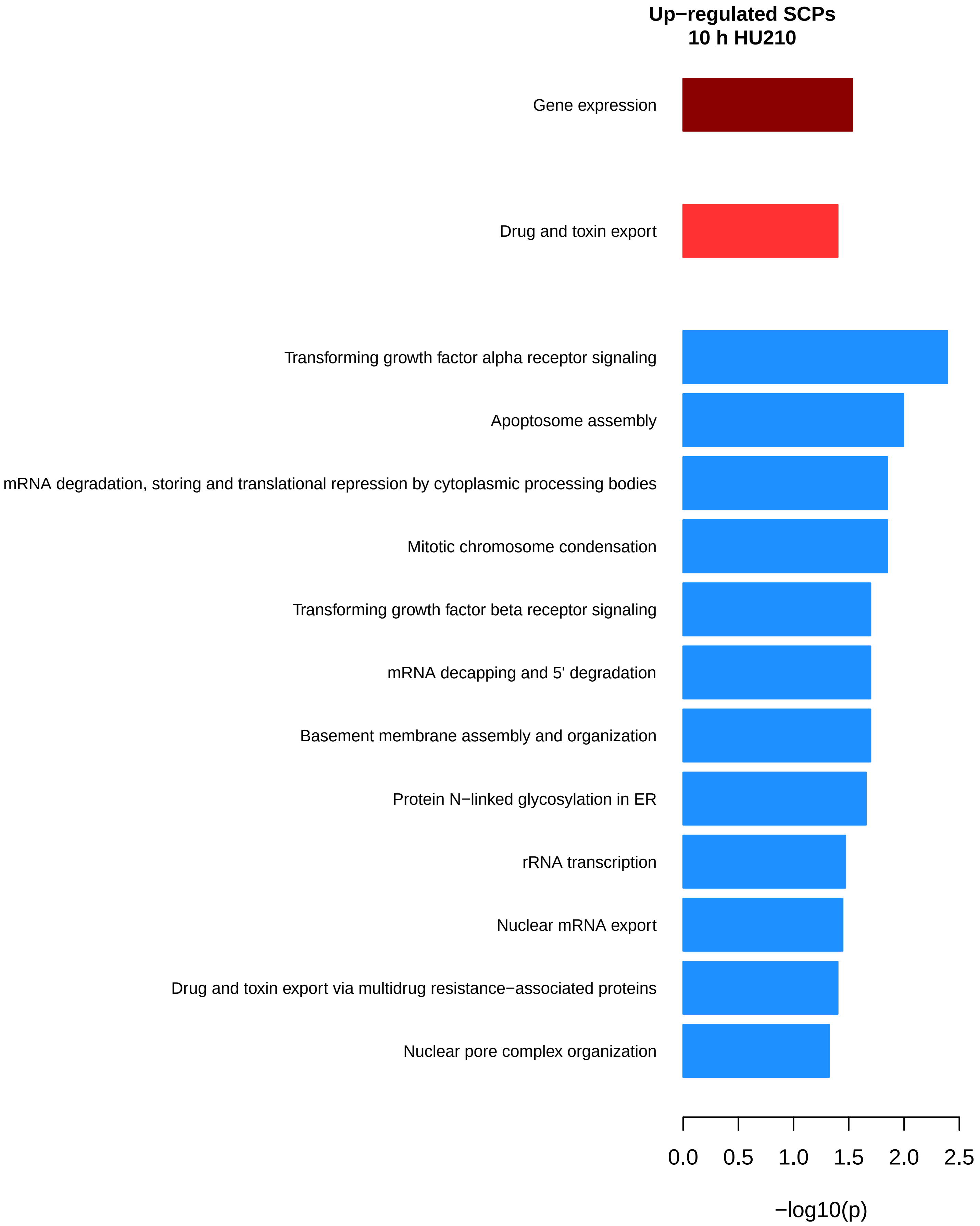

### Suppl. figure 3

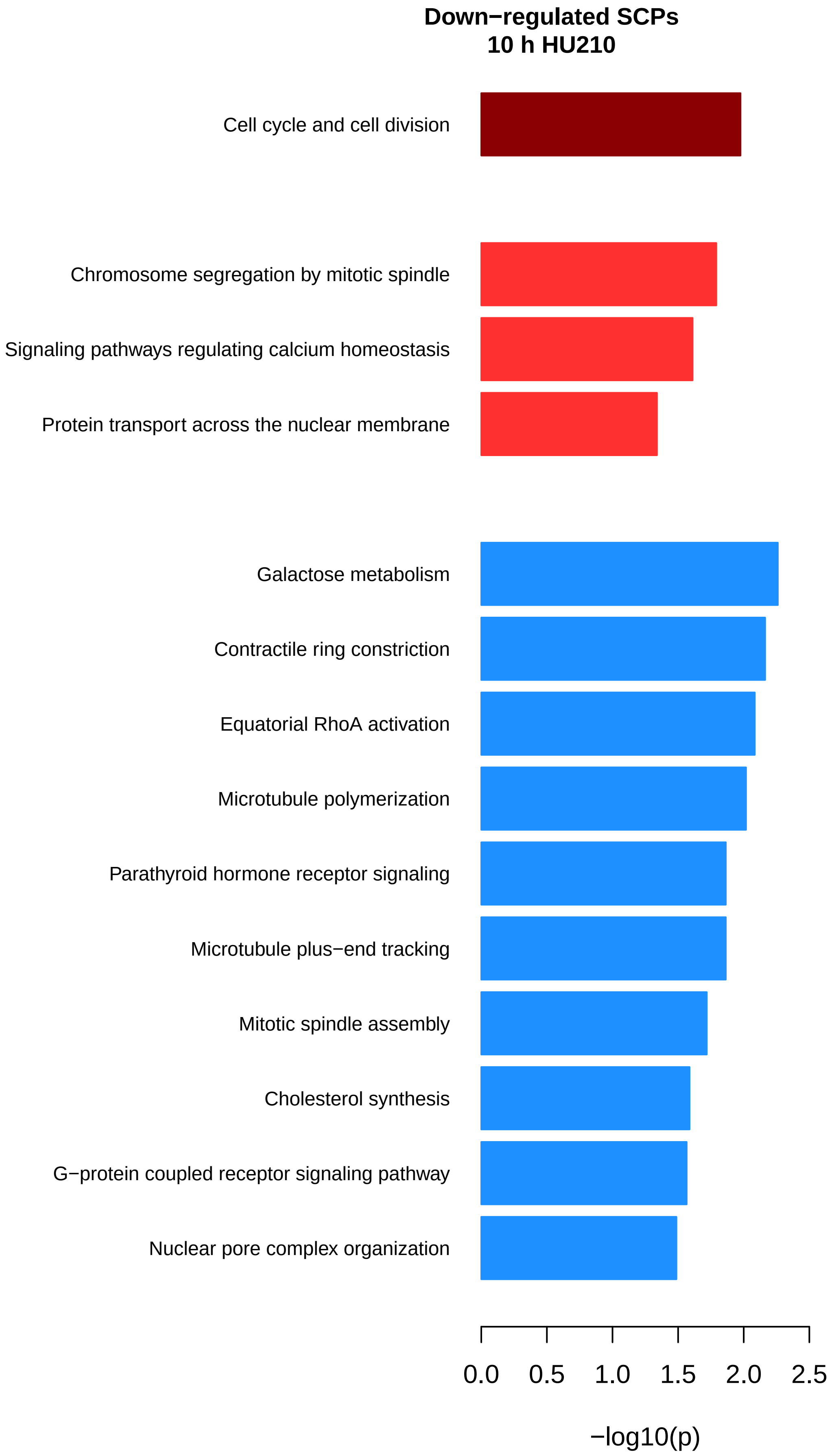

### Suppl. figure 3

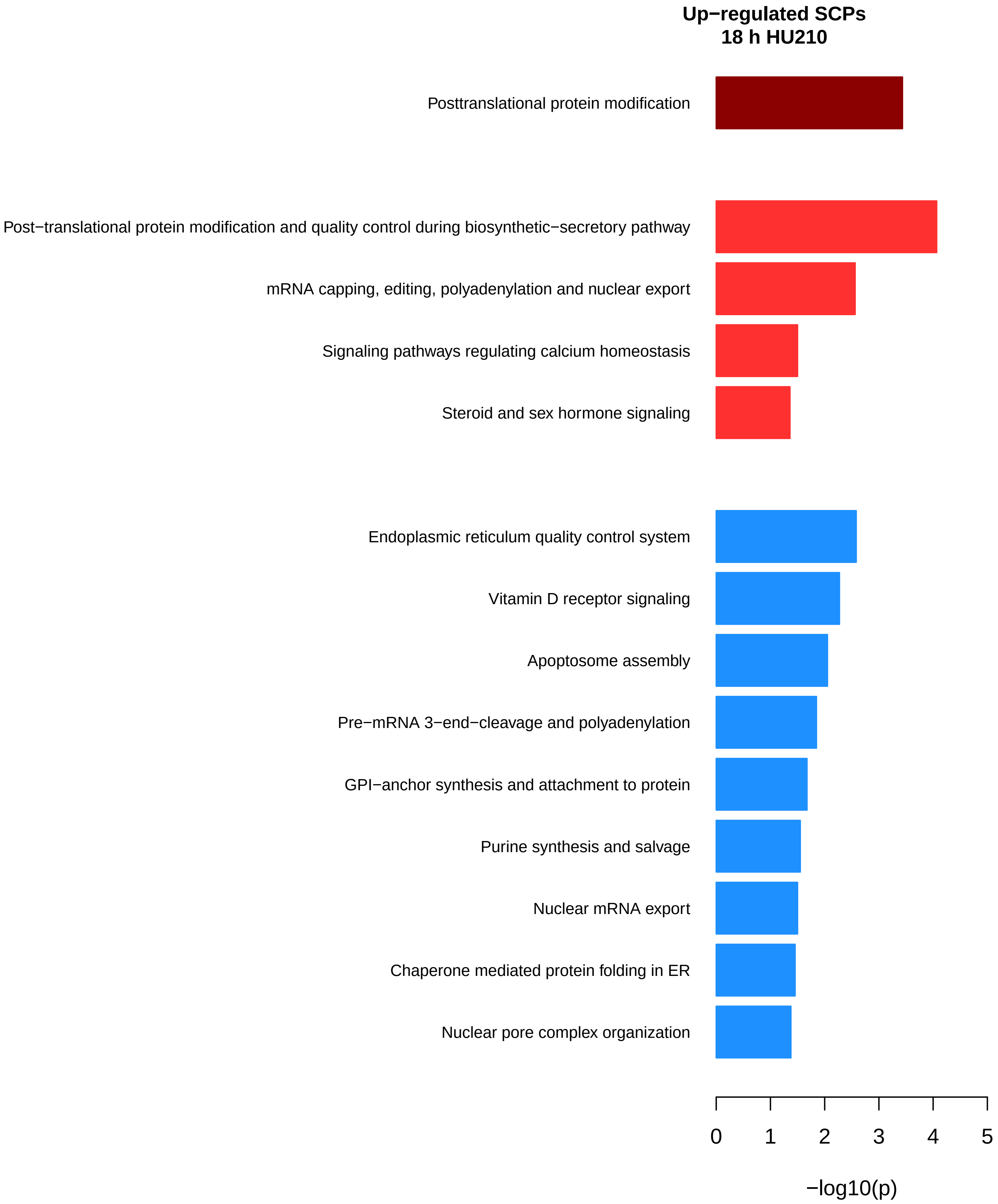

### Suppl. figure 3

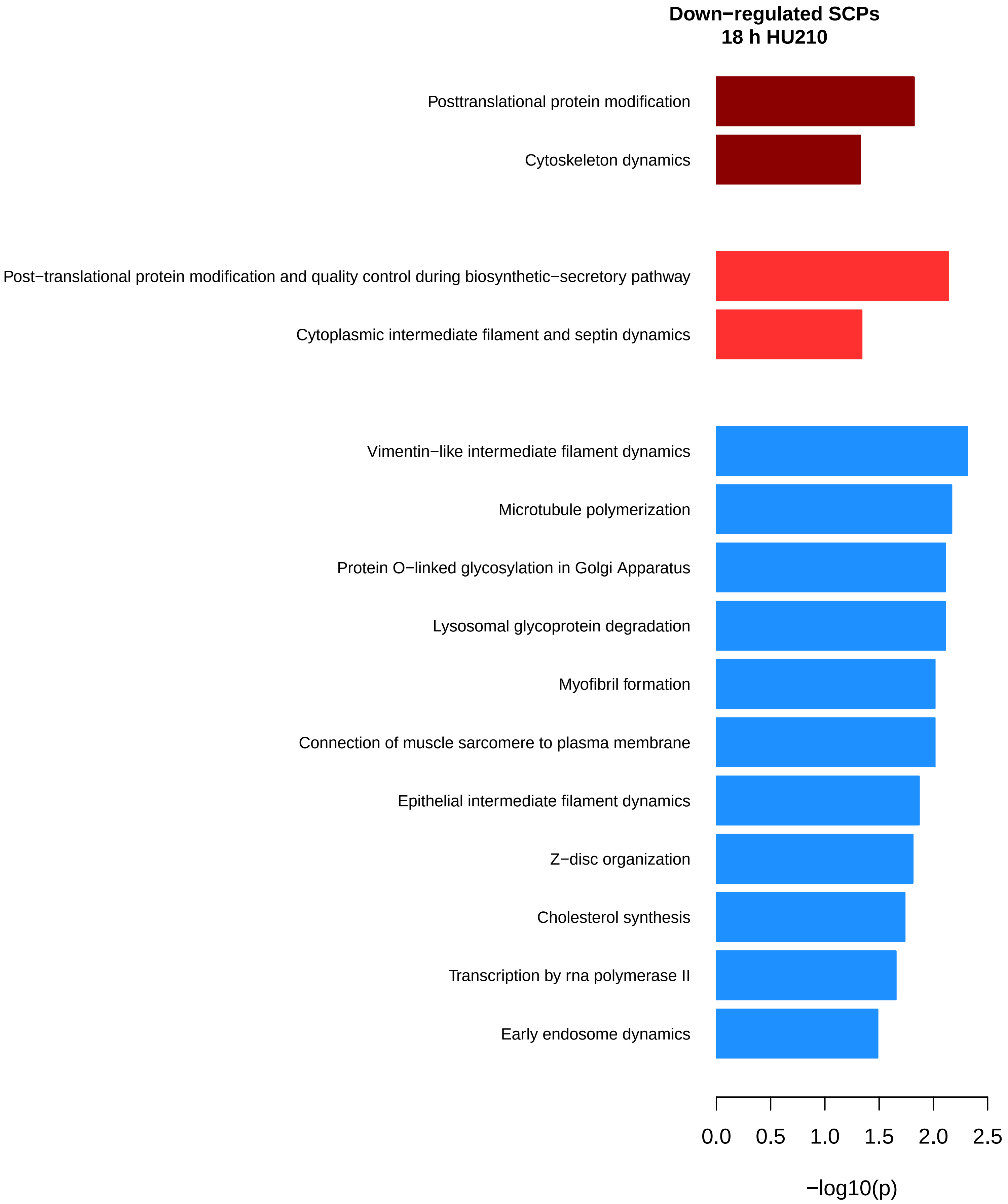

**Suppl. figure 3: Standard enrichment analysis of differentially expressed proteins.** Up- and downregulated proteins were subjected to standard enrichment analysis. Predicted level-1, level-2 and level-3 SCPs with nominal significance (alpha=0.05) are visualized.

### Suppl. figure 4

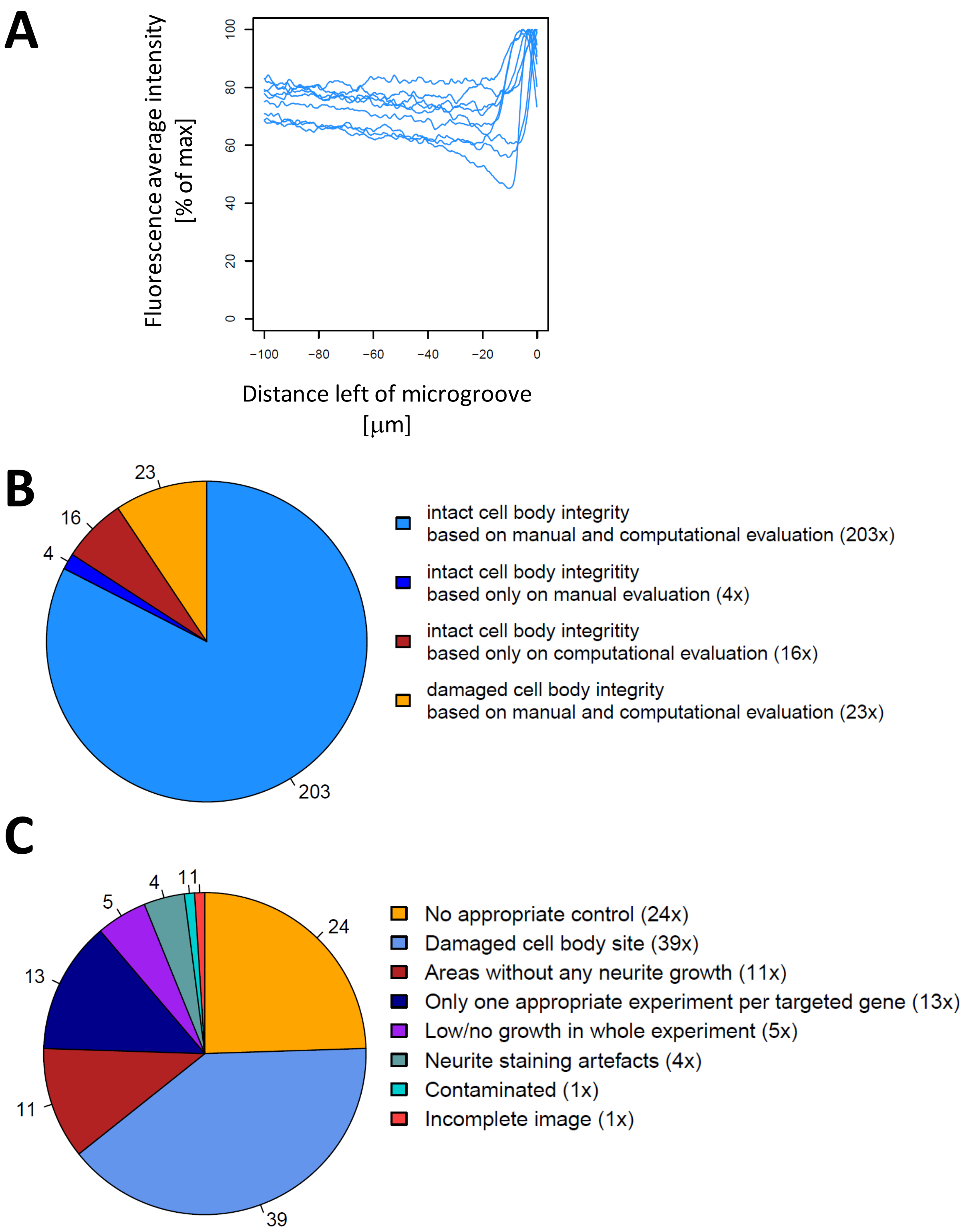

**Suppl. figure 4: Quality control of neurite outgrowth assays.** (A) During plating cells will occupy every space available within the cell body site of the chamber including those right in front of the microgroove. Microscopic analysis will document uniform tubulin staining and an intense line of tubulin staining right in front of the microgroove wall (see Figure 2A). In a few cases the cell body site was damaged during the axotonomy, resulting in a removal of cells in front of the microgroove. Consequently, the staining will be less uniform and the intense line of tubulin staining will appear at an increased distance from the microgroove wall. To identify samples where the cell body side has been damaged during axotonomy, we quantified fluorescence intensities on the cell body site and calculated the average intensity at each distance up to 100 μm left of the microgroove wall. All average intensities for one outgrowth chamber were normalized towards the highest intensity. Intact cell body sites show a fluorescence intensity peak right in front of the microgroove wall as shown for one typical experiment (each blue line represents one targeted or scramble siRNA treated sample). This peak corresponds to the intense line of tubulin staining. If the cell body site was damaged during axotonomy, this peak will be farer away from the microgroove wall. Our algorithm was searching for the highest peak within the first 50 μm left to the microgroove wall (i.e. within a distance that corresponds to 1/3 of the microgroove wall width of 150 μm). If the normalized intensity of that peak was at least 85%, our algorithm labeled the cell body site as intact and as non intact otherwise. (B) Compuational predictions of cell body integrity were manually verified. In 20 out of 246 manually assigned cell body integrity differed from the computationally predicted ones. In these cases we continued with the manual quantifications. (C) Any samples with a damaged cell body site as well as any experiments that did not contain controls with intact cell body sites were removed from any further analysis. Manual investigation also identified further exclusion criteria that lead to the removal of additional samples from the analysis.

### Suppl. figure 5

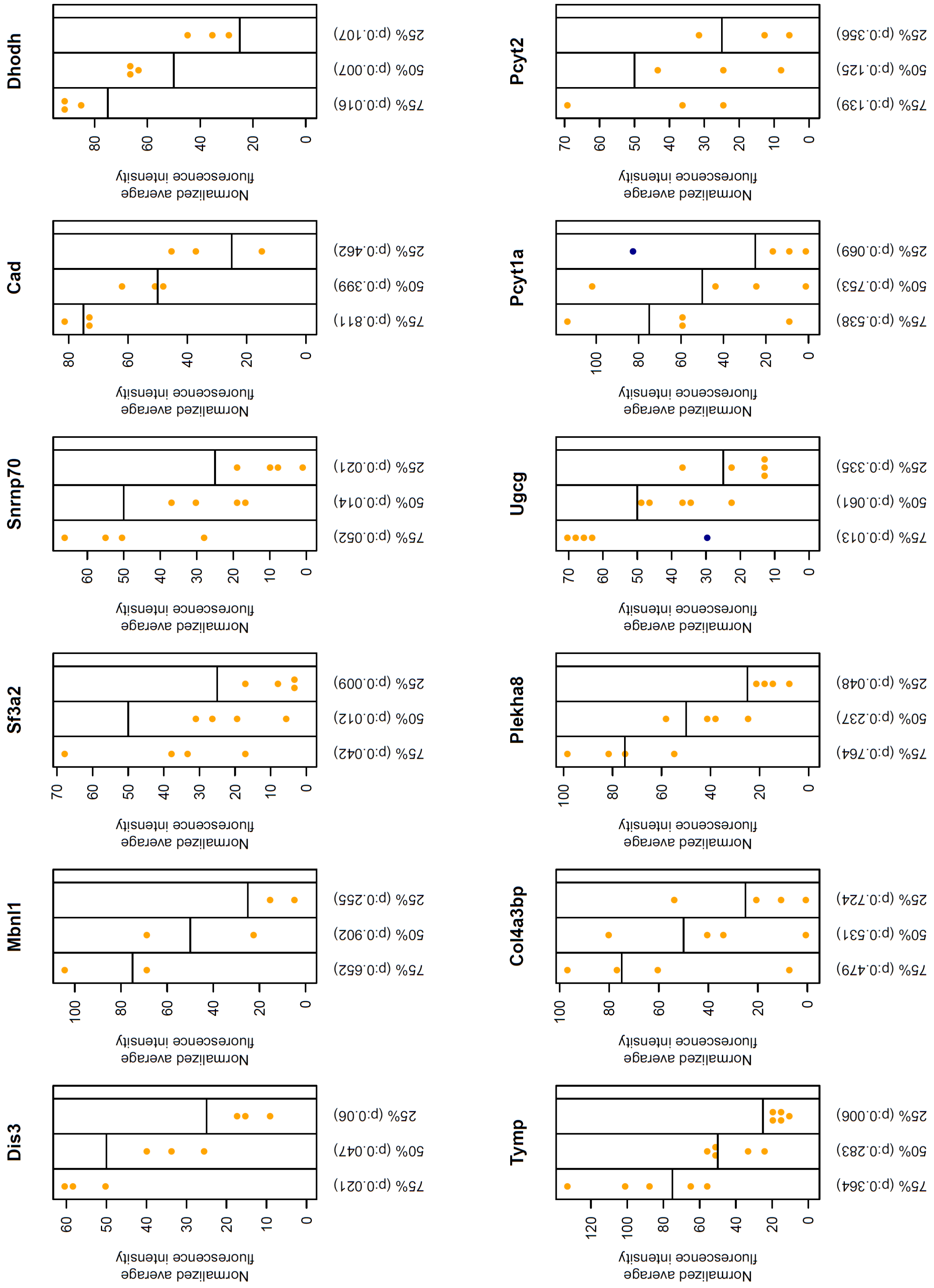

### Suppl. figure 5

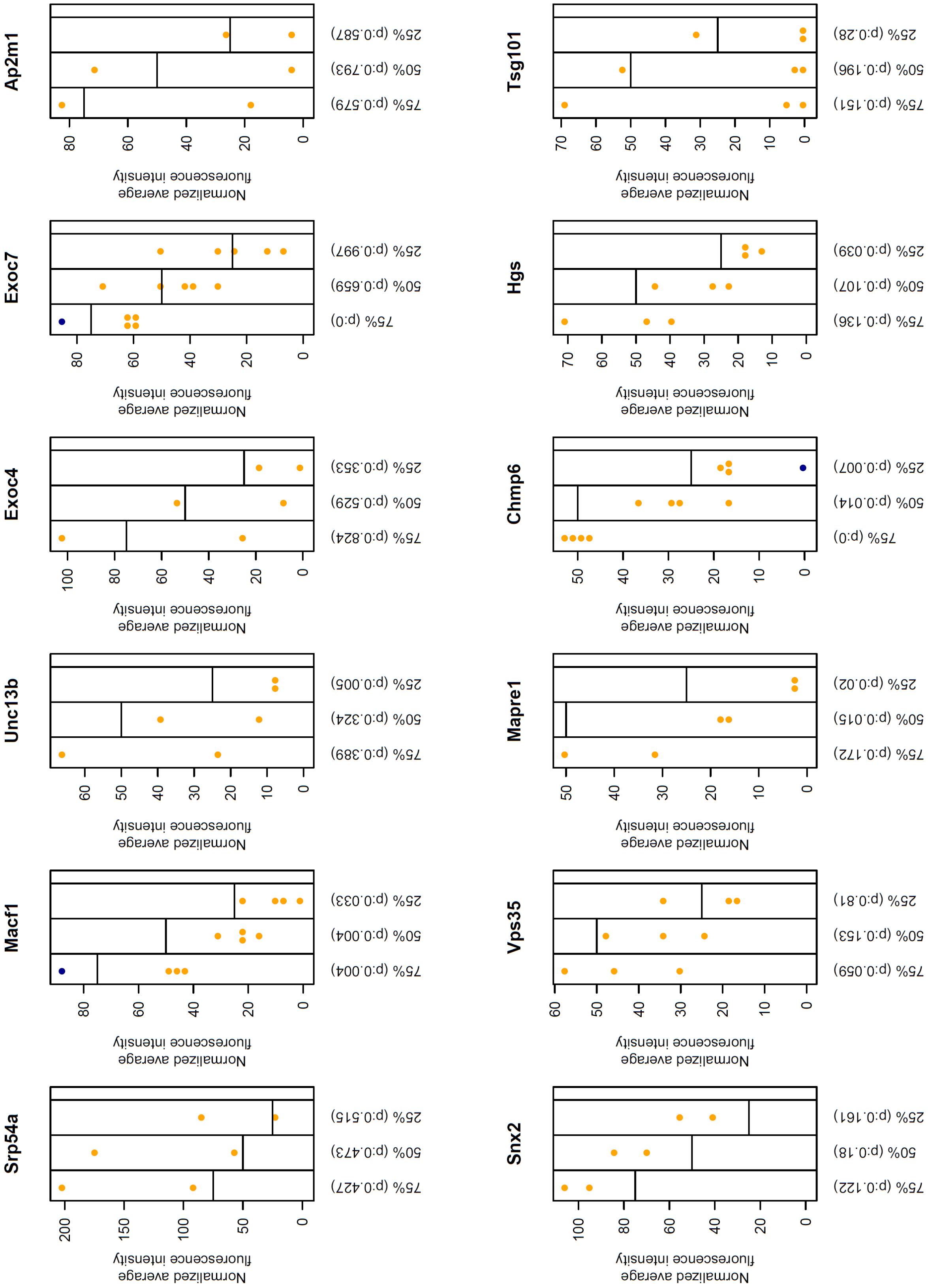

### Suppl. figure 5

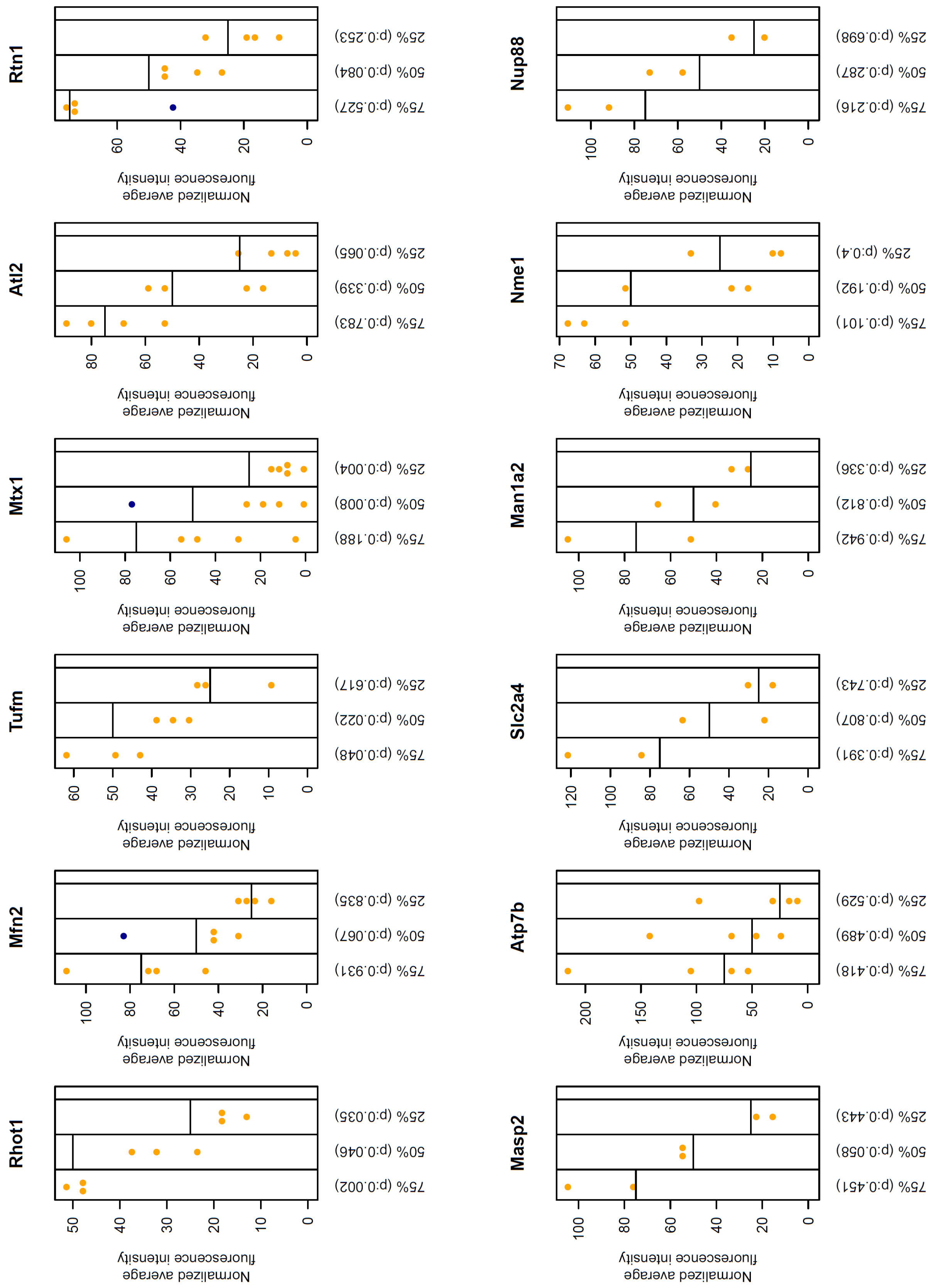

**Suppl. figure 5: Neurite outgrowth results after siRNA mediated gene knock down.** SiRNA knock down results were analyzed as described at figure 2. Orange and blue dots show the normalized outgrowth intensity obtained for the indicated gene in one experiment at the distance that showed indicated control intensity (75%, 50%, 25%). Blue dots are outliers that were identified using Dixon's Q-Test and removed before p-value calculation using one-sample two-tailed ttest. Horizontal lines show reference control intensities.

**Suppl. table 1: Differentially expressed genes and proteins induced by Cb1r sitmulation in Neuro 2A cells.** FDR = 5%, minimum $log_2$(fold change) = +/- $log_2$(1.3) for differentially expressed genes. Nominal p-value = 5% and minimum $log_2$(fold change) = +/- $log_2$(1.1) for differentially expressed proteins.

**Suppl. table 2: Dynamic enrichment results of differentially expressed genes.** Differentially expressed genes (FDR = 5%, minimum $log_2$(fold change) = +/- $log_2$(1.5)) were submitted to dynamic enrichment analysis using MBCO. Seed supplementary figure 1A for details. The column 'Overlap symbols' lists those symbols that are part of the input gene list and the SCP. Numbers in brackets behind each symbols indicate $log_2$(fold change) of that gene.

**Suppl. table 3: Standard enrichment results of differentially expressed genes.** Differentially expressed genes (FDR = 5%, minimum $log_2$(fold change) = +/- $log_2$(1.5)) were submitted to standard enrichment analysis. Seed supplementary figure 1B for details. The column 'Overlap symbols' lists those symbols that are part of the input gene list and the SCP. Numbers in brackets behind each symbols indicate $log_2$(fold change) of that gene.

**Suppl. table 4: Dynamic enrichment results of differentially expressed proteins.**

**Suppl. table 5: Standard enrichment results of differentially expressed proteins.**
